## Supplemental information Text and Figures for "On the relationship between protist metabarcoding and protist metagenome-assembled genomes"

### S1.1 Data structure, preprocessing and analysis

The data on SMAG abundance and SMAG taxonomic assignment was taken from (Delmont et al., 2022) Supplementary data. SMAG abundances in the original data are estimated as a number of mapped reads normalised by the estimated length of the genome (in this case represented by the size of the SMAG assembly augmented using the BUSCO completion score) (Delmont et al., 2022). V9 raw abundance dataset for V9 metabarcodes organised at OTU level was obtained from (Mahé et al., 2022). All the intermediate and accessory datasets generated and used in the process, along with the code and plots are available at `https://github.com/beaplab/Protist_barcode_MAG_correspondence.git`.

The original SMAG dataset contained data on the abundance of 714 non-redundant SMAGs across 939 samples; the original V9 OTU dataset contained data on the abundance of 262722 V9 OTUs across 1084 samples. From the total of 714 SMAGs present in the dataset, we focused on 286 SMAGs that had non-animal and non-NA "genre" taxonomic assignment according to (Delmont et al., 2022) Table S04.

To make dataset processing and analysis computationally feasible, as well as to filter out noise and spurious correspondence, V9 OTU dataset and SMAG datasets are subsetted according to the taxonomic groups into which V9 OTUs and SMAGs are classified. (An all versus all analysis exceeded the memory available on the computational servers that we used.) For this, a subset of both SMAGs and V9 OTUs belonging to the same taxonomic group should be obtained. The V9 OTU taxonomic classification is based on the UniEuk taxonomy. It is hierarchical, i.e. it contains different ranks. The number of ranks is not fixed for each terminal branch. We made an attempt to split V9 OTU taxonomic assignments into 5 ranks which should represent taxonomic groups of differing sizes and genetic affinities to each other. They are termed "supergroup", "taxogroup 1", "taxogroup 2", "taxogroup 3" and "genus", listed here from the highest rank to the lowest. The SMAG and V9 OTU datasets lacked a unified taxonomic system. Although the precision of taxonomic inference is an unresolved issue the SMAGs analysed, we made an attempt to find the best analogies between SMAG and V9 OTU taxonomy. For the SMAGs the "genre" taxonomic assignment was taken as a base, and for the 44 unique non-metazoan taxonomy genres the best possible match to V9 OTU taxonomy was found.

The final system used to subset SMAGs and V9 OTUs by taxonomy is summarised in Figure S4.4.

Prior to the correspondence estimation, we performed zero substitution by smallest count. Next, we computed relative abundance separately for SMAG and V9 abundance datasets. To calculate relative abundance, we used interquartile-log-ratio (iqlr) transformation, as suggested by the propr package developers. Briefly, iqlr transformation is considered a suitable option for our analyses as it is sub-compositionally coherent (i.e. allows subsequent subsetting of datasets without affecting the outcome of the analysis) ((Quinn et al., 2019, 2017) - "propr" package vignettes, FAQ section). Relative abundance calculation and 0 substitution were carried out by an algorithm adapted from the "propr" function of the corresponding package (the adapted version is available at `https://github.com/beaplab/Protist_barcode_MAG_correspondence/tree/main/R_scripts/calculation`).

After replacing zeros and obtaining relative abundance counts of SMAGs and V9 OTUs, we split both datasets into taxonomic subsets. The resulting subsets contained relative abundance data for SMAGs of a given genre and V9 OTUs within the taxogroup that matches SMAG genre at a particular taxonomic level (see Figure S4.4, Figure S1). At this point, we removed V9 OTUs with total raw abundance count $\geq 100$ from subsequent calculations.

Two subsets of SMAG and V9 OTU relative abundances for a particular taxonomic group of a given taxonomic rank were merged into a single dataset, containing relative abundances of both SMAGs and V9 OTUs across *Tara* samples. For the dataset combining both SMAG and V9 OTU relative abundances, pairwise proportionalities/correlations were estimated; the values obtained for all SMAG-V9 OTU pairs were retained.

We estimated proportionality and correlation with algorithms adapted from the "propr" package. We tested five different metrics to assess the correspondence between V9 OTU and SMAG relative abundances: Spearman's correlation and all four different proportionality metrics implemented in the propr package *rho*, *phi*, *phs* and *vlr*. The formulas for each metric used are provided elsewhere (Quinn et al., 2019, 2017), also see the "propr" package vignette "An Introduction to Proportionality" section for more detail. A long discussion that accompanied the development and testing of different methods of proportionality calculation was conducted elsewhere, with the advocation of *rho* metrics as the most appropriate (Erb and Notredame, 2016).

Spearman's correlation and *rho* proportionality outperformed other correspondence metrics according to the result of SMAG controls and simulations (see Section S2.2, Section S4.4). Hence, the results for *rho* metrics are the ones shown in the main text, and the correspondence calculated by using all metrics is presented in the supplement.

After obtaining correspondence estimates with all metrics for all SMAG-V9 OTU pairs within each taxonomic group of each taxonomic rank, the V9 OTUs that match every given SMAG were ranked by correspondence value from lowest to highest: the V9 OTU with the highest correspondence being ranked as "top-1 match", the second highest "top-2 match", and so forth.

All SMAG-V9 OTU matches obtained for *rho* proportionality and Spearman's correlation were automatically filtered to match the following criteria: a) the difference between the mean correspondence estimate of top-1 match from the shuffled subset + 1 standard deviation and the correspondence estimate of a given pair is > 0.05; b) the difference between the correspondence estimate of a given top-nth ranked pair and the next top-n+1-ranked pair is > 0.05.

When the proportionality/correlation estimates for the positive controls from simulations fell within the interval of the mean proportionality/correlation estimate of top-1 match from the shuffled subset + 1 standard deviation (and therefore failed to meet the first candidate match selection criterion), there was no need to apply an additional filter for the match proportionality/correlation estimates to be higher or equal to simulated positive control matches, because the negative control values of the simulation were predominantly lower than that of shuffling baselines for Spearman's correlation and *rho* metrics Section S4.4.

For each SMAG genre, we filtered candidate V9 OTU matches from the two lowest rank taxonomic subsets, from 2 correspondence metrics (*rho* and Spearman's correlation), which resulted in a combination of candidate matches for 4 different settings. For each match, we assessed consistency between the taxonomic subsets and metrics using a 0 to 4 scoring system, 0 being the least consistent. If a V9 OTU for a given SMAG is recovered as the match of the same rank across two taxonomic subsets, the match scores 2; if the OTU is recovered in both subsets, but the ranks of the match do not match, it scores 1; if the V9 OTU is recovered as candidate match in only 1 taxonomic subset, it scores 0. As there are two different metrics, we estimated the consistency between taxonomic subsets for both, with the resulting maximum score of 2+2=4. The same system was applied to estimate the consistency of V9 OTU being recovered across each of the two metrics. We also counted the number of times a given V9 OTU was recovered as a candidate match for other SMAGs within the same genre as the SMAG examined, and outside of the given SMAG taxonomy genre.

Using the automated procedure described above, we manually assessed candidate matches to produce a list of matches which can be potentially informative relations of V9 OTUs and SMAGs, although not necessarily implying a direct correspondence of a SMAG and a matched V9 OTU to a single biological entity. Depending on the consistency between taxonomic subsets and metrics, and on the uniqueness of V9 OTU match within/outside of a SMAG taxonomic genre, different categories reflecting the quality of the SMAG-V9 OTU match were assigned to each pair: "Less Probable" (very low confidence) < "Probable" (low confidence) < "More Probable" (medium confidence) < "Yes" (high confidence).

We note that the final list of potentially matching pairs was assessed manually and is not meant to be exhaustive. We also make available the initial list of candidate matches which did not undergo manual filtering to permit further exploration of particular taxonomic groups or SMAG-V9 OTU pairs.

#### S2.2 Simulated datasets control

To test the performance of different proportionality/correlation estimation methods, we created artificial datasets of V9 OTU and SMAG abundances, where abundance counts are randomly distributed (i.e., no intentional correspondence between features in two datasets). To those mock datasets with randomly distributed abundances, we added a single control pair of SMAG and V9 OTU with abundances that exactly correspond to each other.

We created 46 independent pairs of SMAG and V9 OTU simulated datasets using the "metaSPARSim" package (Patuzzi et al., 2019) function metaSPARSim and the input parameters described below.

We estimated the input parameters for simulations from the real SMAG and V9 OTU datasets. The "Library size" parameter for V9 OTU dataset simulation was the sum of raw V9 OTU abundances across each sample; the "Intensity" input parameter was estimated as the mean raw abundance for each V9 OTU; finally, the "Variability" parameter contained variances of raw abundances estimated for each V9 OTU using "var" function from R base (R Core Team, 2020).

We converted the input SMAG abundance dataset to raw abundance counts prior to estimating the input parameters for simulation based on it. For this, we first calculated the total number of mapped raw reads for each sample from the total number of reads per station multiplied by the fraction of mapped reads (corrected by length); Then the relative abundance of each SMAG in each sample was calculated by dividing percent of mapped reads normalised by SMAG's BUSCO by the total percent of mapped reads per sample and subsequently multiplying by the number of raw reads per sample.

The obtained matrix of raw SMAG counts, normalised with the GMPRX function from metaSPARsim package, as suggested by developers, was used as a template for metaSPARsim "estimate_parameter_from_data" function, with the "perc_not_zeros" parameter lowered to 0.01 (due to the high fraction of zero counts in the dataset). "Intensity" and "Library size" input parameters obtained from the "estimate_parameter_from_data" function outputs was later divided by 10000, and "variability" parameter was correspondingly divided by $10000^2$ to lower the actual raw read counts that otherwise are too high for computationally feasible simulation process. (As the proportionality/correlation calculation algorithm operates with relative abundances, such artificial reduction of input parameters by several magnitudes does not affect the final input dataset for proportionality/correlation calculation.) The obtained input parameters were used to simulate the SMAG count matrix. The dimensions of the simulated datasets were the same as that of the real data.

The abundances for the control V9 OTU were generated by random sampling of all abundance counts of V9 OTU simulated dataset, with the 0 count frequency maintained the same as in the rest of the data. Later, the abundance vector of the control V9 OTU was divided by the average abundance count per sample, and multiplied by the average abundance count of SMAGs in the same sample of the simulated SMAG dataset, as follows:

$$abSMAGcontrol_{i,j} = \frac{abV9control_{e,j} * \sum_{x=0}^{i-1} abSMAG_{x,j}/i}{\sum_{x=0}^{e-1} abV9_{x,j}/e} \qquad (1)$$

where $abSMAGcontrol_{i,j}$ is the abundance count of the control SMAG in the sample $j$ with $i$ number of features; $abV9control_{e,j}$ is the abundance of the control V9 OTU in the sample $j$ (i.e. the sample corresponding to the sample $j$ from SMAGs dataset) with $e$ number of features; $abSMAG_{x,j}$ and $abV9_{x,j}$ are the abundances of individual feature (SMAG or V9 correspondingly) number $x$ of a sample $j$.

As a result, the control SMAG relative abundances across samples completely corresponded with the relative abundances of the control V9 OTU. The control abundance vectors were then merged into the corresponding simulated datasets (with trackable identifiers).

Proportionalities/correlations for each of 23 pairs of simulated datasets were calculated by the algorithm explained above Section S1.1. A control V9 OTU - SMAG pair was included in each taxonomic subset (taxonomic subsets, in this case, representing the total number of SMAGs and V9 OTUs in a real data subset Figure S4.4). To save computational time, the number of V9 OTUs in the subset was set not to exceed 20000, so every subset larger than that was artificially limited to 20000.

To verify that 23 replicates of simulation controls were sufficient, we analysed how the standard deviation of proportionality/correlation for the top-1st ranked matches of the positive control SMAG with the positive control V9 OTU changed with increasing the number of replicates Figure S5. In the majority of cases, the standard deviation reached a plateau before the 23rd replicate.

After calculating proportionality/correlation on the simulated datasets, the matches obtained for the features mimicking SMAGs and V9 OTUs could be classified into 4 categories: 1) "Both": the positive control SMAG is matched to the positive control V9 OTU, i.e. the two features designed to be proportional are recovered as a match (indicating that the method produced the "true positive" match) 2) "None": a random (negative control) SMAG is matched to a random (negative control) V9 OTU, - i.e. two features created with random relative abundance patterns are recovered as a true match. This category can be used as a negative control, representing the "baseline" values obtained just by chance. 3) "Only V9, not SMAG": a random (negative control) SMAG is matched to the positive control V9 OTU, i.e. the V9 OTU feature that is expected to match the positive control SMAG is matched with a random feature instead, indicating a false positive match. 4) "Only SMAG, not V9": the positive control SMAG is matched with a random (negative control) V9 OTU, i.e. the V9 OTU feature that is expected to match the positive control SMAG is matched with a random feature instead, indicating a false-positive match..

We estimated the number of matches from each of the 4 categories ("None", "Both", "Only V9, not SMAG", and "Only SMAG, not V9") for each of the top-10 best ranked pairs in simulated datasets. The results are summarised in Figure S6. The higher the number of replicates for which true-positive "Both" category matches are recovered as the top-1st rank, the lower the amount of false-positive match categories ("Only V9, not SMAG", "Only SMAG, not V9") and the bigger the difference in proportionality/correlation values between the true positive and false positive/true negative matches, the better is the performance of a given correspondence metric. From Figure S6 it is clear that the metrics *phi*, *phs* and *vlr* failed to recover the true positive matches as high-rank pairs in the absolute majority of cases. This result suggests that *rho* and Spearman's correlation metrics are the most appropriate ones, at least for simulated data that mimics the real dataset we analysed.

Simulation outcomes can be used as a control for non-unique V9 OTU matches as well. Scenarios in which a single V9 OTU matches multiple SMAGs in the taxonomic subset are common Figure S3. In cases of a single V9 OTU matching several SMAGs (a redundant match), we need to know if there is a way to discriminate the actual match from the false-positive SMAG-V9-OTU match. For this, we can investigate the correspondence values and ranks of the false-positive matches ("Only V9, not SMAG") obtained from simulation, and compare them to those of redundant V9 OTU matches from real data. The closer are the correspondence values and ranks of real redundant matches to that of simulated ones, the more likely the match is truly redundant Figure S4 Figure 2(c).

#### S3.3 Shuffling

As a negative control, we calculated the proportionality/correlation values that would be obtained by chance for each taxonomic subset from datasets of similar sizes and counts.

After iqlr transformation (i.e. after obtaining relative abundances) of the V9 OTU dataset, the columns and rows of the relative abundance matrix were shuffled (1000 iterations) using "randomizeMatrix" function from the R package "picante" (Kembel et al., 2010). Although the relative abundance values were shuffled, the taxonomic assignments of V9 OTUs were kept the same (as they were to be used for creating "mock" taxonomic subsets of the same size as the original ones). Next, proportionality/correlation was calculated for the SMAG and V9 OTU shuffled relative abundance datasets, via the same algorithm as described in Section S1.1.

We performed the procedure of shuffling followed by correspondence estimation in 132 independent replicates, i.e. on the 132 independently randomly shuffled V9 OTU relative abundance datasets. For each taxonomic subset, we calculated the mean, maximum/minimum and standard deviation of correspondence values for each rank of SMAG-V9 OTU pairs, over 132 replicates.

The +/- standard deviations from the mean for each SMAG-V9 OTU pair rank are shown as ribbons in the plot Figure 2, Figure S4.4 .

As Figure S4.4 illustrates, the standard deviation values of the top-1 ranked SMAG-V9 OTU pairs stopped fluctuating before the number of replicates reached 100, indicating that the number of replicates was sufficient and its increase would be unlikely to improve error rates.

#### S4.4 Single-cell amplified genome/reference genome controls

We extracted 18S sequences from genome assemblies of single-cell amplified genome (SAG) assemblies from (Delmont et al., 2022) and from 4 reference genomes with ANI score match $\geq$99% to MAGs from (Delmont et al., 2022) by performing a BLAST search with 18S sequences with corresponding taxonomic assignments from the EukRibo database used as queries Table S2. 18S sequences obtained from SAG and reference genome assemblies were used as queries to search against the V9 OTU Tara dataset (Mahé et al., 2022); barcodes matching with percent identity 99.2 were obtained. Afterwards, we compared the barcodes corresponding to the 18S sequences found in the SAGs/reference genomes assemblies to the ones that were matched with SAGs/reference genomes based on the relative abundance across samples (by the algorithm described in Section S1.1).

As seen from Figure S4.4 (orange circles and purple asterisks in the subplots for corresponding taxonomic subsets), the proportionality metrics *rho* and Spearman's correlation (Spearman's correlation) recovered the V9 OTU corresponding to the V9 sequence present in the genome assembly more frequently and with higher ranks when compared to the *phi*, *phs* and *vlr* proportionality metrics (in particular, for SMAGs of "unidentified Chrysophyceae", "unidentified Bicosoecida", "Ostreococcus", "Micromonas" and "unidentified Bacillariaceae 1" genres). These results support *rho* proportionality and Spearman's correlation as the most appropriate metrics for our datasets.

However, even when applying *rho* proportionality and Spearman's correlation, V9 OTUs which are not present in control SMAGs are recovered as highly proportional to control SMAGs, in taxa such as "unidentified Bicosoecida", "unidentified Bacillariaceae 1", unidentified Bacillariaceae 3", "unidentified Chrysophyceae". Such false positive correspondence can either arise as an artefact of data preprocessing/proportionality calculation or represent an actual feature of the initial dataset. In the latter case, this means that in the raw data, there is more than a single V9 OTU within the taxonomic subset which has an abundance pattern matching those of the SMAG of interest.

Figure S8 illustrates that more than one V9 OTU shows an abundance pattern which matches that of one SMAG. Hence, in cases where the V9 OTU present in the SMAG sequence was not recovered as the best-corresponding one, this is likely due to the presence of another V9 OTU with a similar abundance pattern in the raw data or due to the small number of stations with non-zero SMAG abundance, and not due to the artefacts of data processing and analysis.

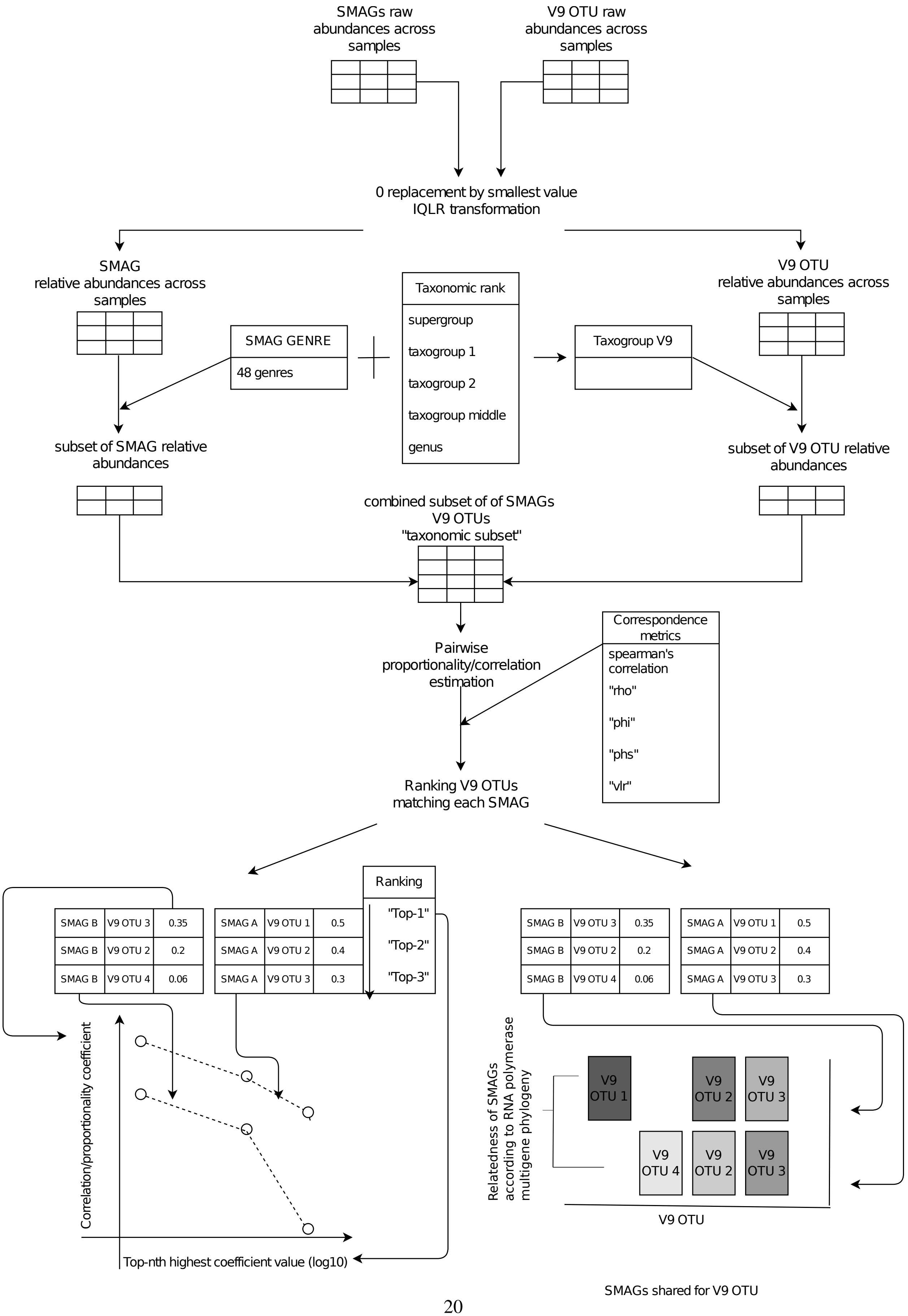




Figure S1: **Schematic outline of the algorithm used for data preprocessing, correspondence estimation and data visualization.**

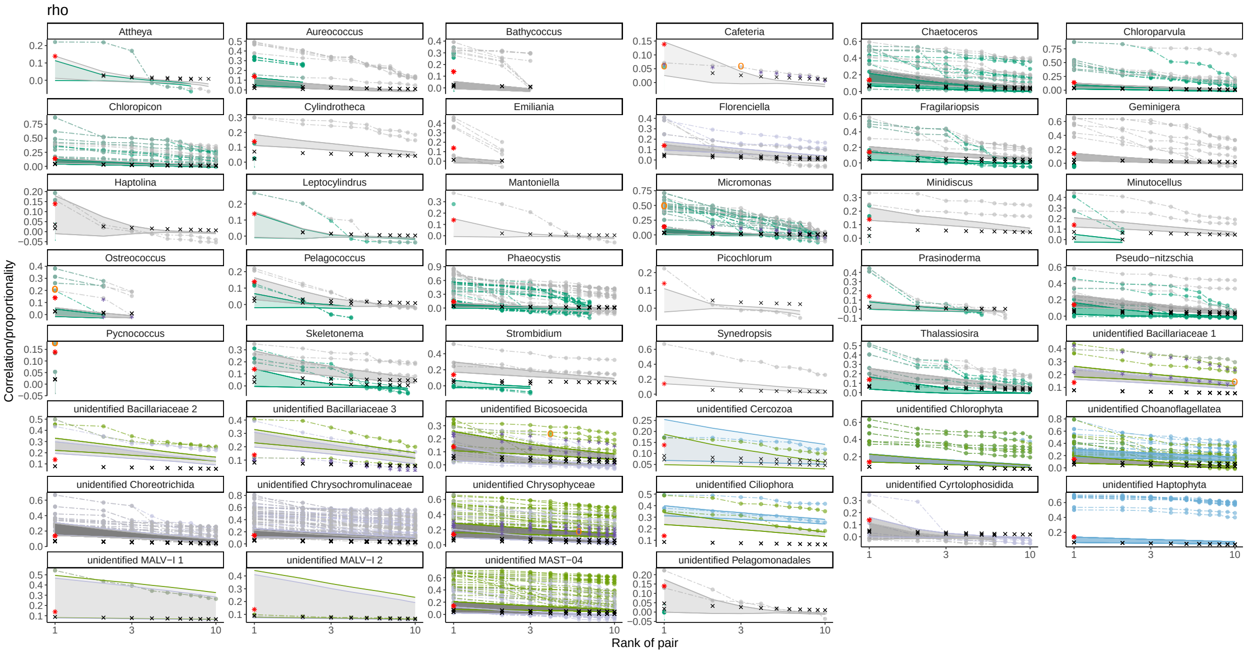


Figure S2: a

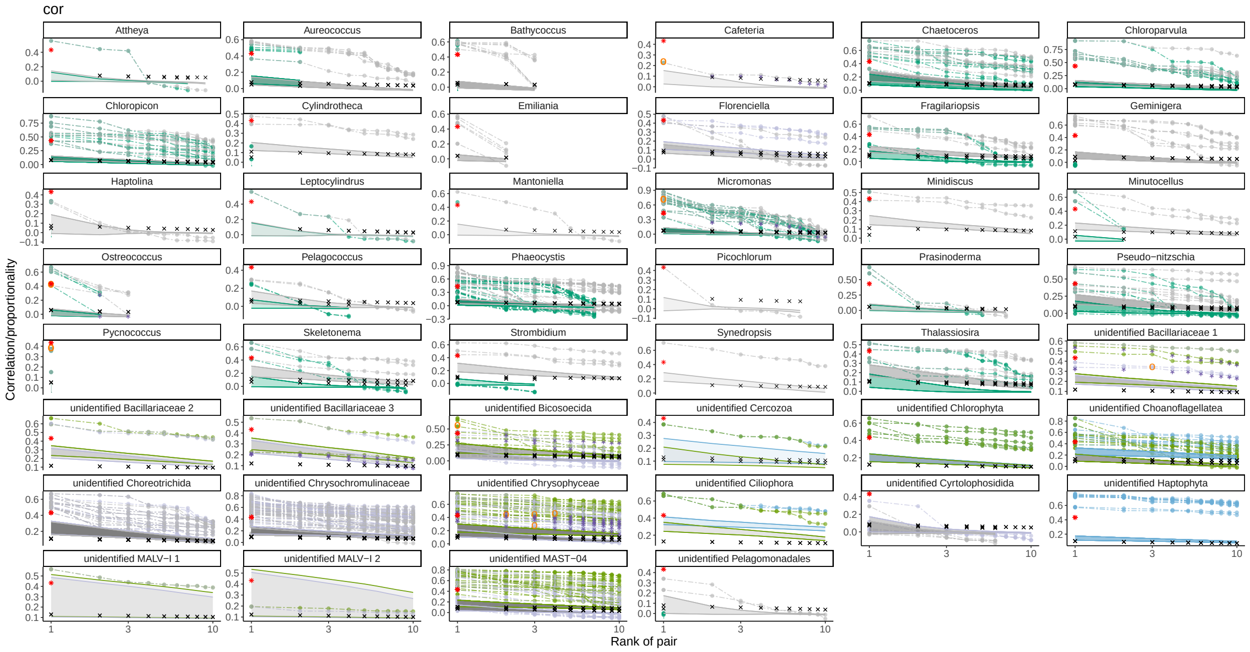


Figure S2: b

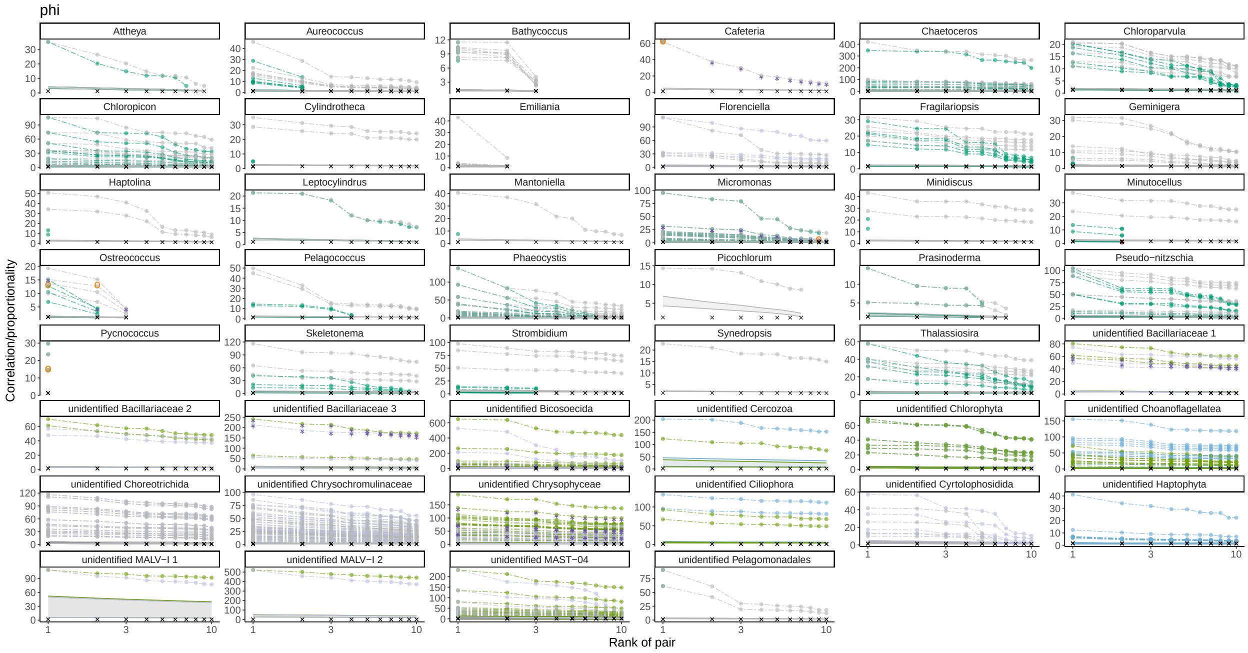


Figure S2: c

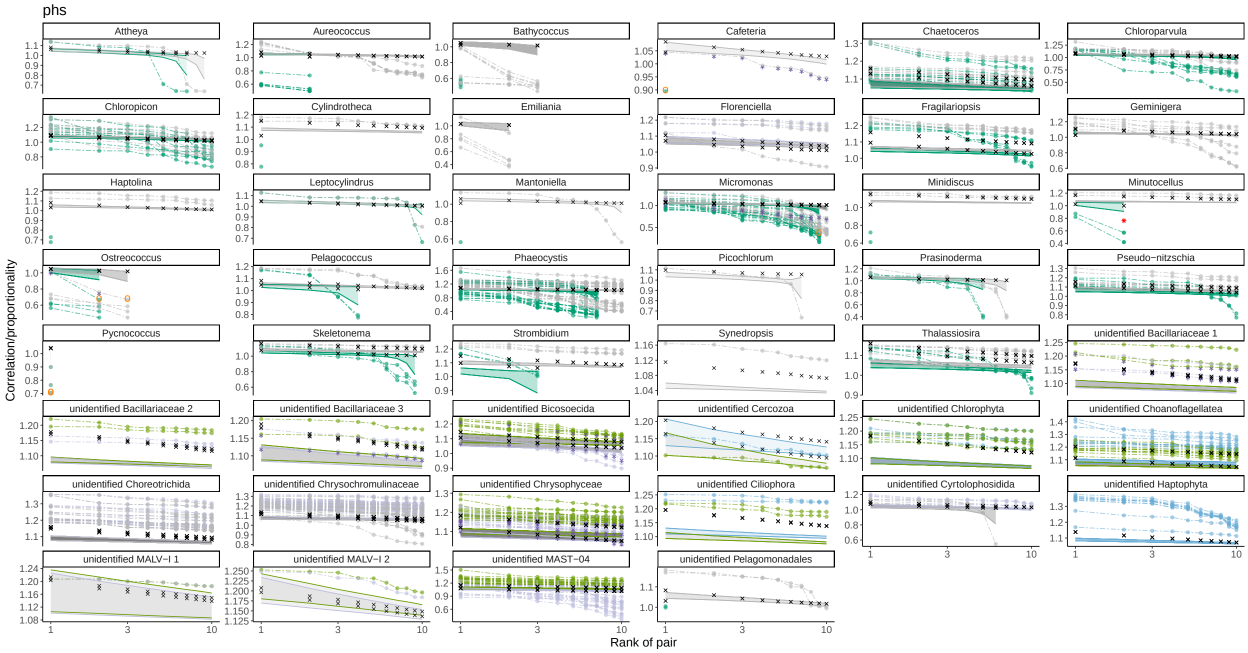


Figure S2: d

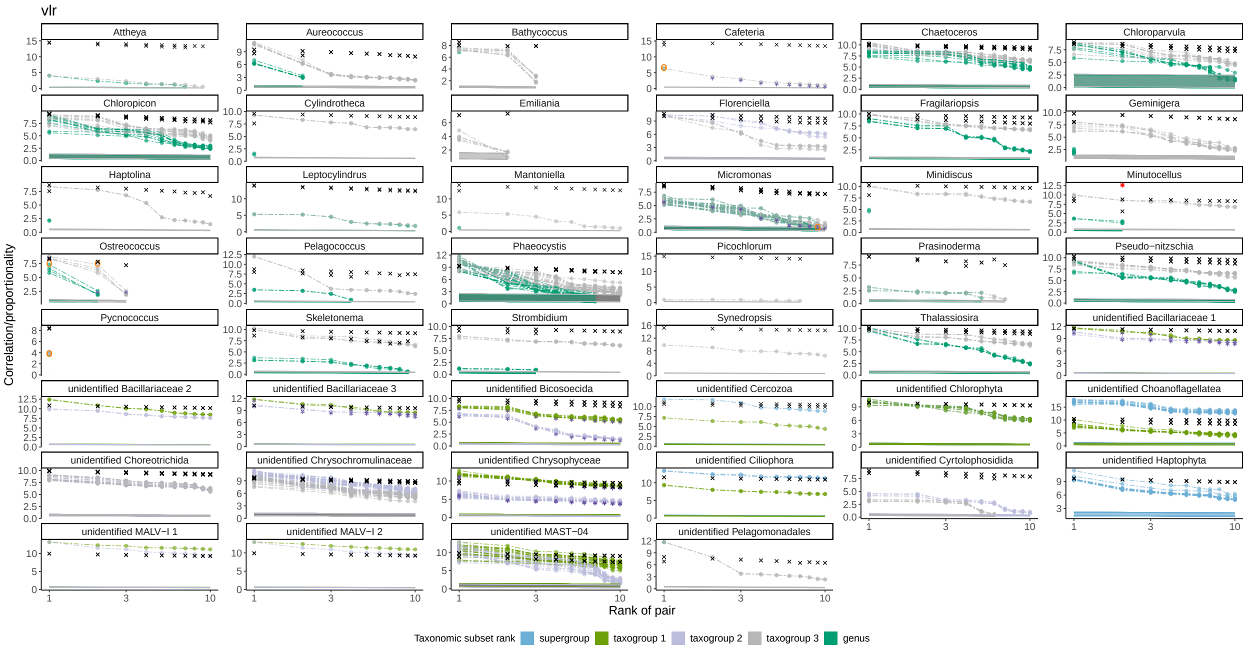


Figure S2: e

Figure S2: **The proportionality of the 10 best V9 OTU matches within two taxonomic subsets, for each SMAG** Proportionality values of V9 OTUs paired with SMAGs are shown on the vertical axis, and the rank of the match on the horizontal axis (log scale). Colours represent the rank of the V9 OTU taxonomic subset (from highest to lowest rank: supergroup, taxogroup 1, taxogroup 2, taxogroup 3, or genus). The filled ribbons represent the mean +/- standard deviation of proportionality obtained for the shuffled datasets of a given taxonomic subset (negative controls). Lines for SMAGs highlighted in a given scenario in Figure 2 are indicated with points. Colour corresponds to the rank of the taxonomic subset shown. Red asterisks represent the mean proportionality/correlation obtained for the true-positive matches from artificially simulated datasets S2.2; black crosses represent mean values of the true-negative controls from simulations; for taxonomic subsets containing one SMAG, black crosses show the values for false-positive controls in which the target SMAG is matched to the wrong V9 OTU S2.2. Orange circles around some of the points represent the SMAG-V9 OTU pairs that could be verified by using the SAGs control as the true-positive match (i.e. the sequence of V9 OTU matching a SMAG was recovered from the SAG/reference genome assembly). Purple asterisks indicate that the V9 OTU matching control SMAG was not found in SAG/reference genome assembly, and likely is a false-positive match S4.4. Each subfigure represents statistics for different correspondence estimation metrics, with the metrics indicated in the header of the subfigure: (a) *rho* proportionality, (b) Spearman's correlation, (c) *phi* proportionality, (d) *phs* proportionality, (e) *vlr* proportionality.

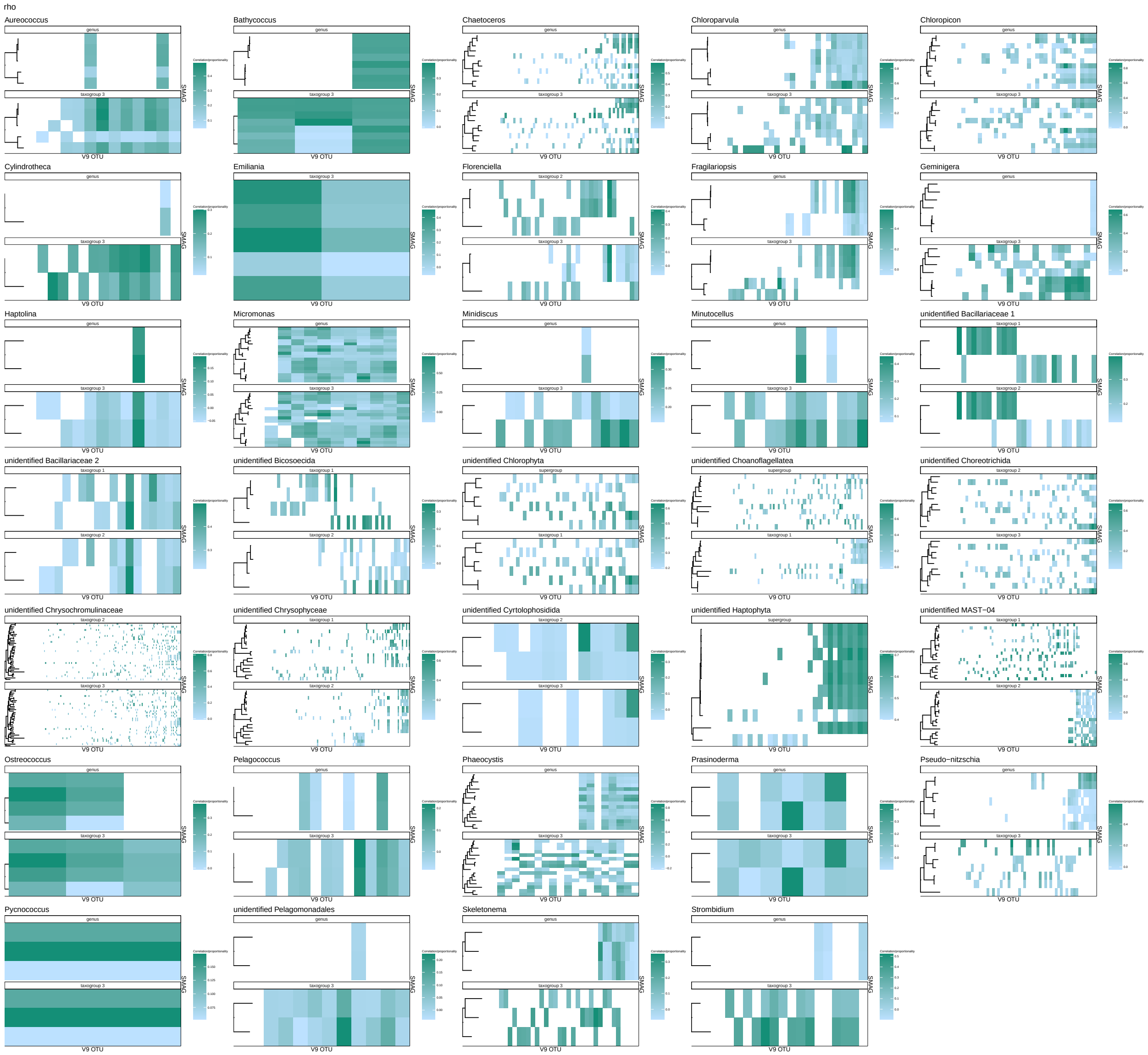
rho
Aureococcus
Bathycoccus
Chaetoceros
Chloroparvula
Chloropicon
Cylindrotheca
Emiliania
Florenciella
Fragilariopsis
Geminigera
Haptolina
Micromonas
Minidiscus
Minutocellus
unidentified Bacillariaceae 1
unidentified Bacillariaceae 2
unidentified Bicosoecida
unidentified Chlorophyta
unidentified Choanoflagellatea
unidentified Choreotrichida
unidentified Chrysochromulinaceae
unidentified Chrysophyceae
unidentified Cyrtolophosidida
unidentified Haptophyta
unidentified MAST−04
Ostreococcus
Pelagococcus
Phaeocystis
Prasinoderma
Pseudo−nitzschia
Pycnococcus
unidentified Pelagomonadales
Skeletonema
Strombidium
V9 OTU
SMAG

Figure S3: a

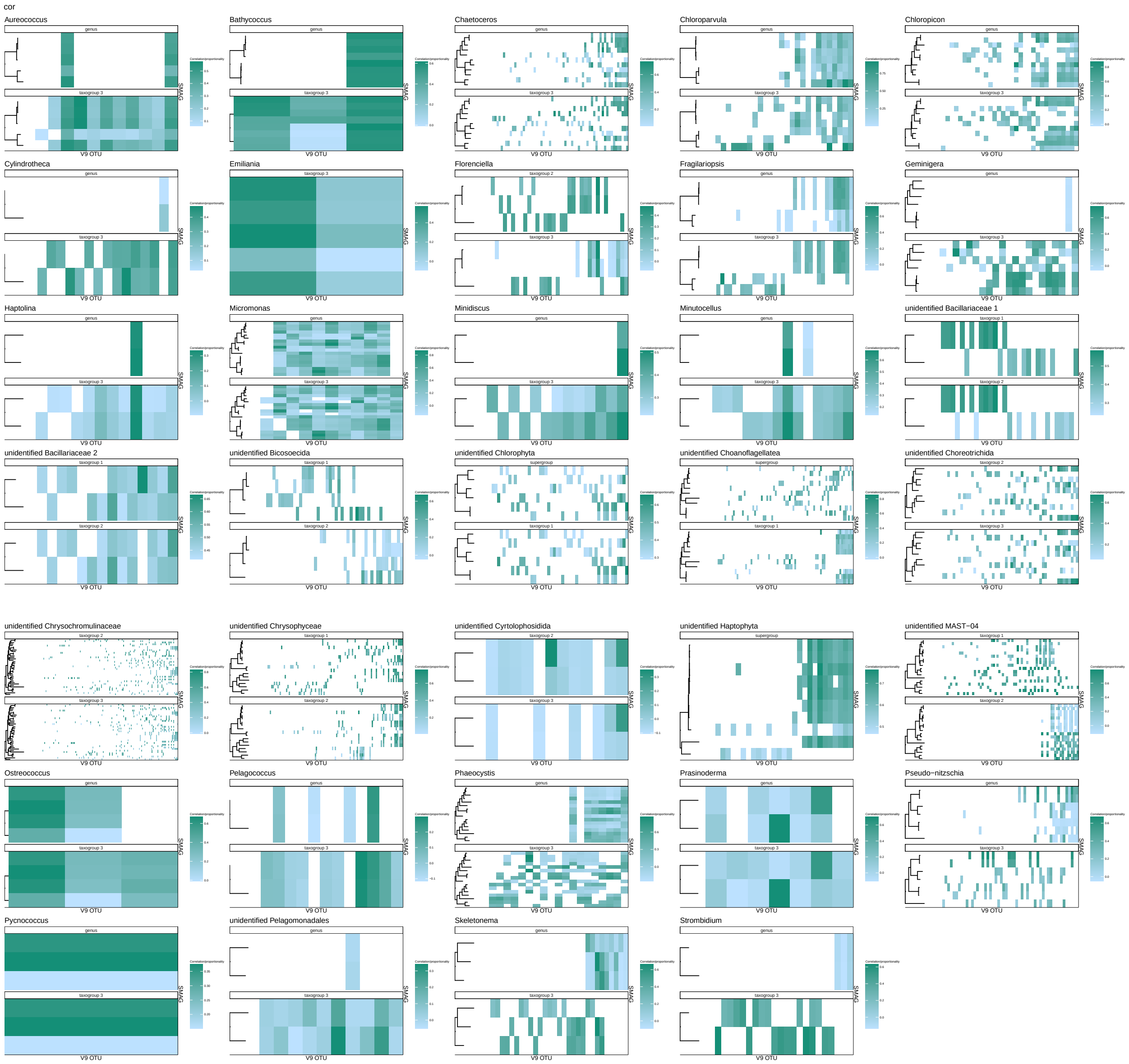
cor
Aureococcus
Bathycoccus
Chaetoceros
Chloroparvula
Chloropicon
Cylindrotheca
Emiliania
Florenciella
Fragilariopsis
Geminigera
Haptolina
Micromonas
Minidiscus
Minutocellus
unidentified Bacillariaceae 1
unidentified Bacillariaceae 2
unidentified Bicosoecida
unidentified Chlorophyta
unidentified Choanoflagellatea
unidentified Choreotrichida
unidentified Chrysochromulinaceae
unidentified Chrysophyceae
unidentified Cyrtolophosidida
unidentified Haptophyta
unidentified MAST−04
Ostreococcus
Pelagococcus
Phaeocystis
Prasinoderma
Pseudo−nitzschia
Pycnococcus
unidentified Pelagomonadales
Skeletonema
Strombidium
V9 OTU
SMAG

Figure S3: b

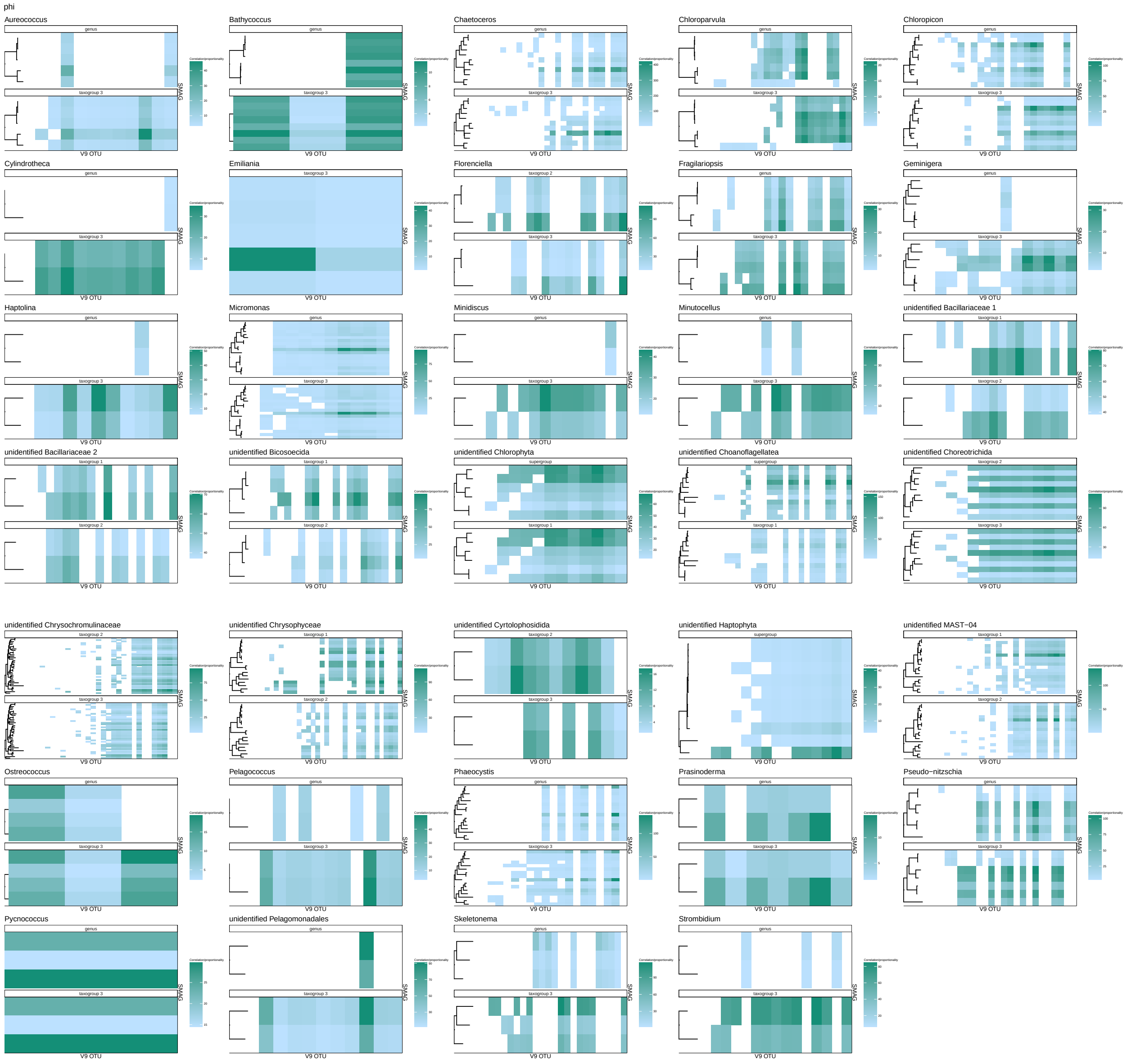
phi
Aureococcus
Bathycoccus
Chaetoceros
Chloroparvula
Chloropicon
Cylindrotheca
Emiliania
Florenciella
Fragilariopsis
Geminigera
Haptolina
Micromonas
Minidiscus
Minutocellus
unidentified Bacillariaceae 1
unidentified Bacillariaceae 2
unidentified Bicosoecida
unidentified Chlorophyta
unidentified Choanoflagellatea
unidentified Choreotrichida
unidentified Chrysochromulinaceae
unidentified Chrysophyceae
unidentified Cyrtolophosidida
unidentified Haptophyta
unidentified MAST−04
Ostreococcus
Pelagococcus
Phaeocystis
Prasinoderma
Pseudo−nitzschia
Pycnococcus
unidentified Pelagomonadales
Skeletonema
Strombidium
V9 OTU
SMAG

Figure S3: c

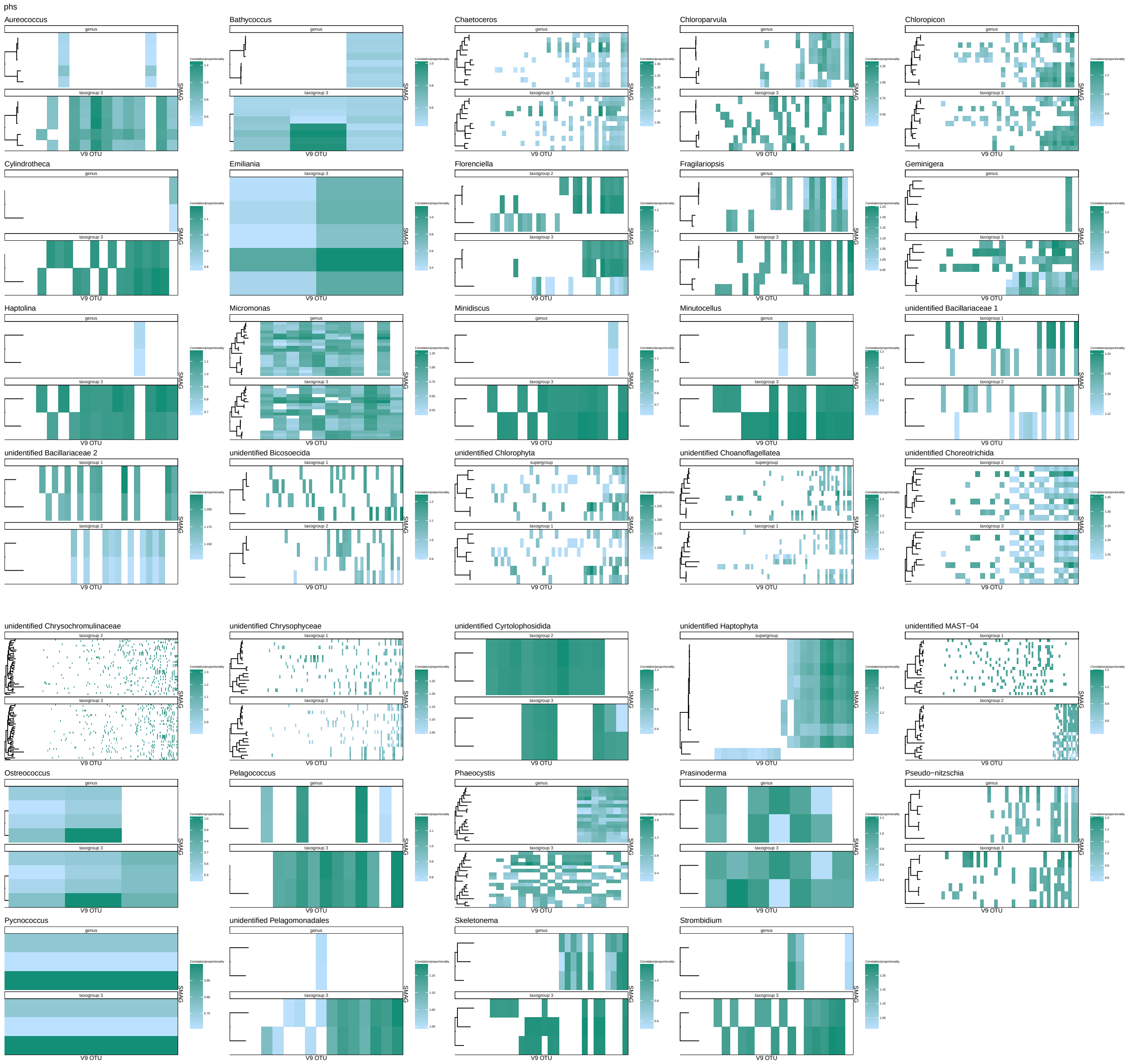
phs
Aureococcus
Bathycoccus
Chaetoceros
Chloroparvula
Chloropicon
Cylindrotheca
Emiliania
Florenciella
Fragilariopsis
Geminigera
Haptolina
Micromonas
Minidiscus
Minutocellus
unidentified Bacillariaceae 1
unidentified Bacillariaceae 2
unidentified Bicosoecida
unidentified Chlorophyta
unidentified Choanoflagellatea
unidentified Choreotrichida
unidentified Chrysochromulinaceae
unidentified Chrysophyceae
unidentified Cyrtolophosidida
unidentified Haptophyta
unidentified MAST−04
Ostreococcus
Pelagococcus
Phaeocystis
Prasinoderma
Pseudo−nitzschia
Pycnococcus
unidentified Pelagomonadales
Skeletonema
Strombidium
V9 OTU
SMAG

Figure S3: d

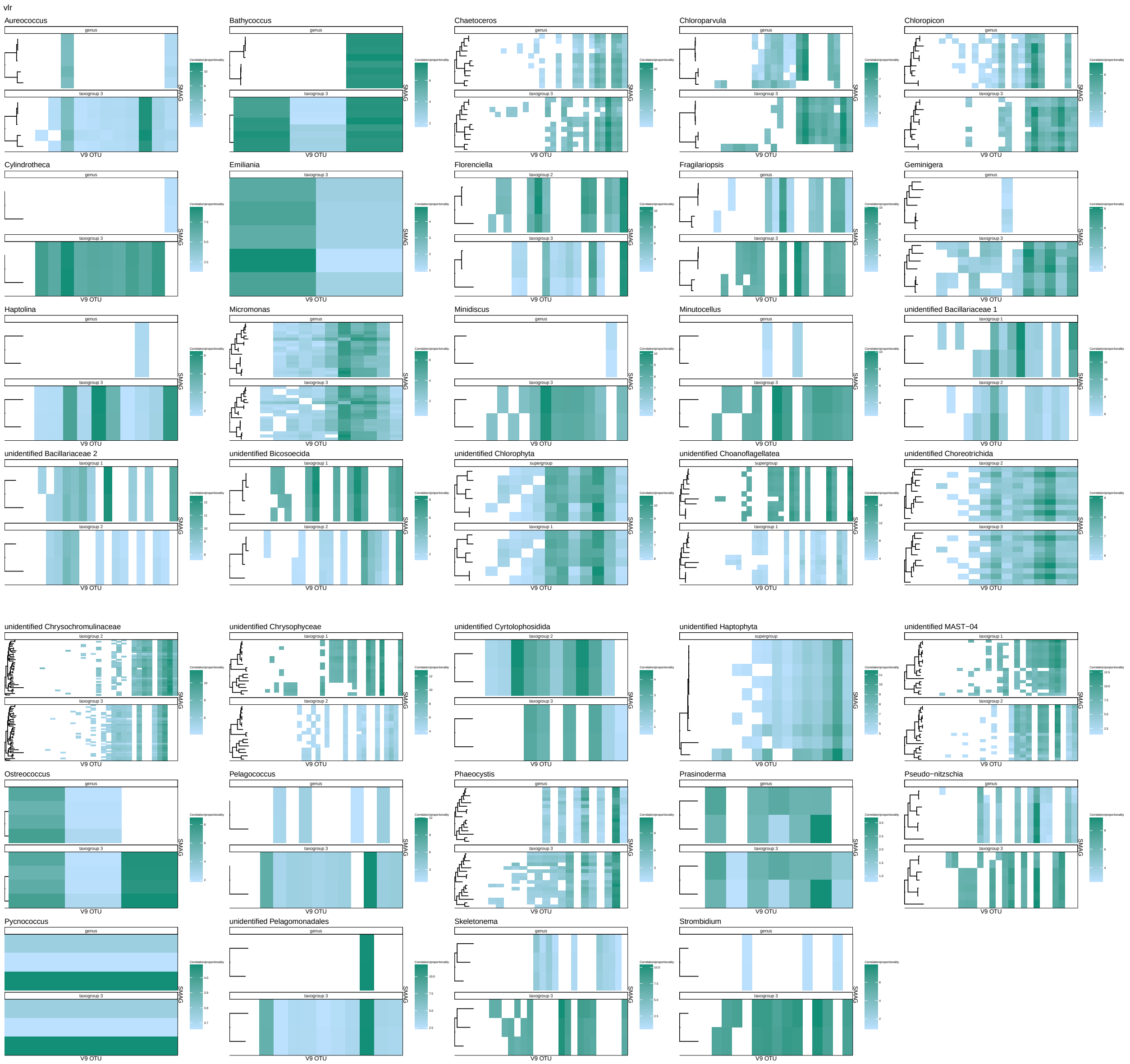


Figure S3: e

Figure S3: **The number of SMAGs potentially matching a given V9 OTU.** The top 10 best-ranked SMAG-V9 OTU matches within the two lowest-ranked taxonomic subsets are represented as tiles. The horizontal axis represents V9 OTUs included in the taxonomic subset, the vertical axis shows SMAGs, and each individual tile thus corresponds to a given SMAG-V9 OTU pair. The fill colour of each tile shows the proportionality value of the match (the scale may differ among subfigures). This plot illustrates the uniqueness of each match: if several SMAGs match the same V9 OTU, several tiles will be present at the same coordinate on the horizontal axis, across different points on the vertical axis and vice versa. The rank of the V9 OTU taxonomic subset presented is indicated at the top of each plot. Phylogenetic trees on the left represent the genetic relatedness of SMAGs in each taxonomic subset, with tree topology and branch lengths from the RNA-polymerase multigene phylogeny of (Delmont et al., 2022). The taxonomic rank for the subset presented is indicated at the top of each subplot. Only those taxonomic subsets with >1 SMAG are presented on the plots. Each subfigure represents statistics for different correspondence estimation metrics, with the metrics indicated in the header of the subfigure: (a) *rho* proportionality, (b) Spearman's correlation, (c) *phi* proportionality, (d) *phs* proportionality, (e) *vlr* proportionality.

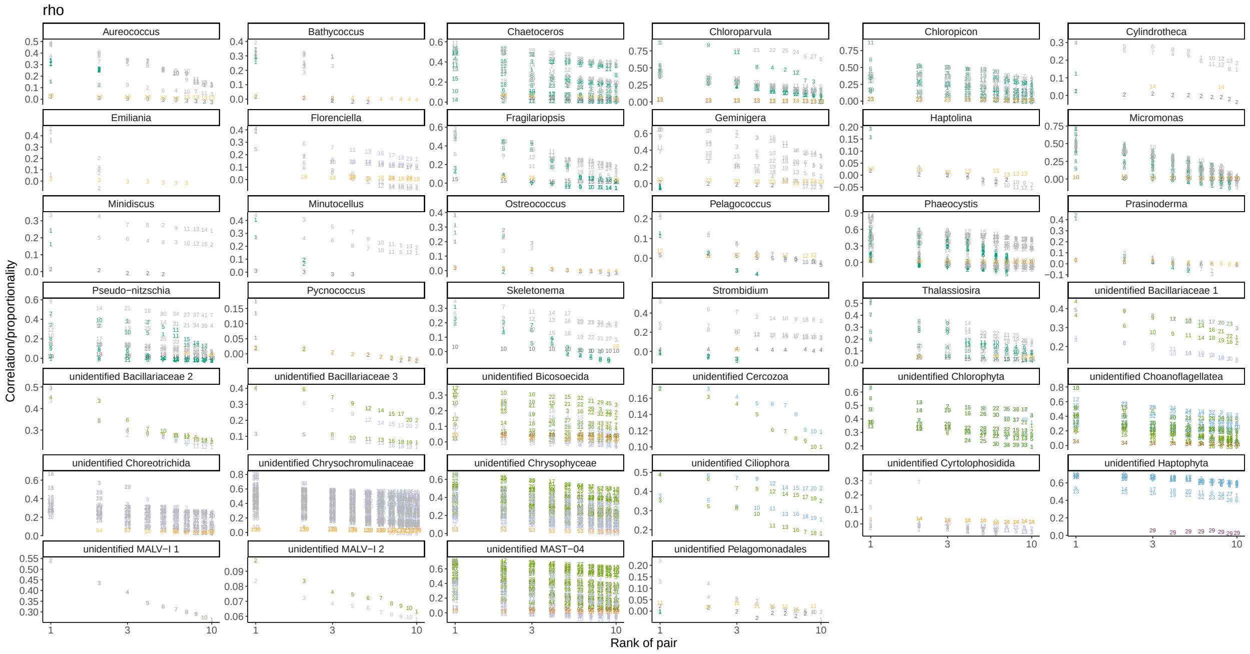


Figure S4: a

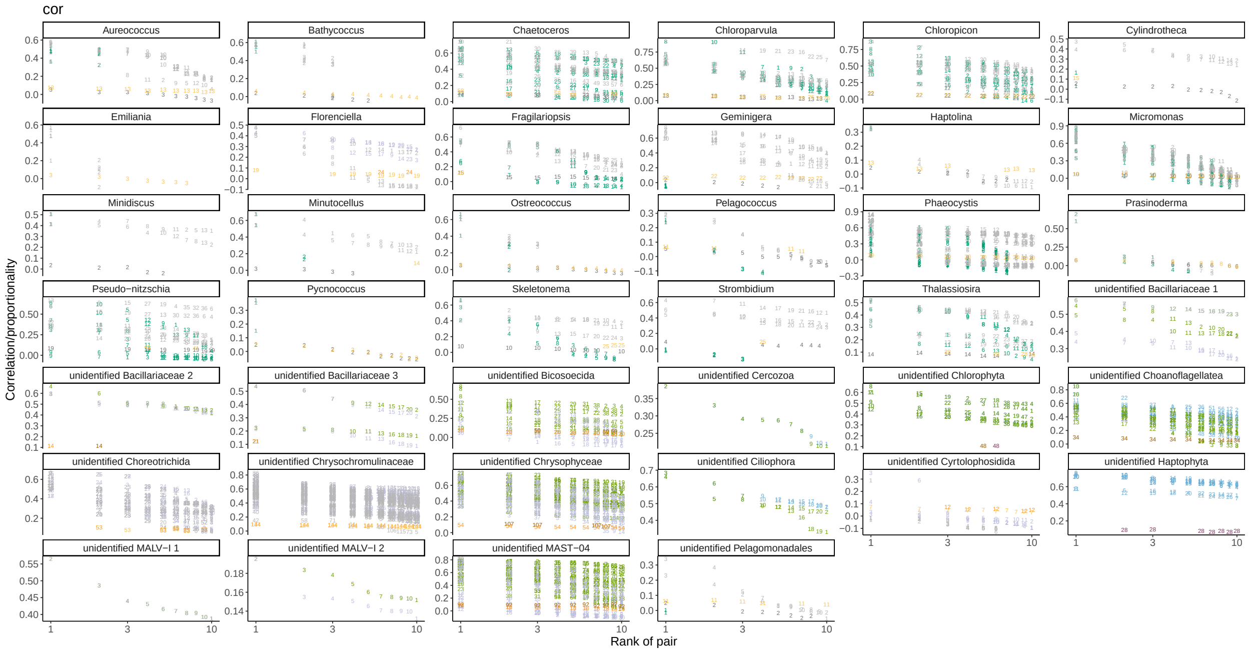


Figure S4: b

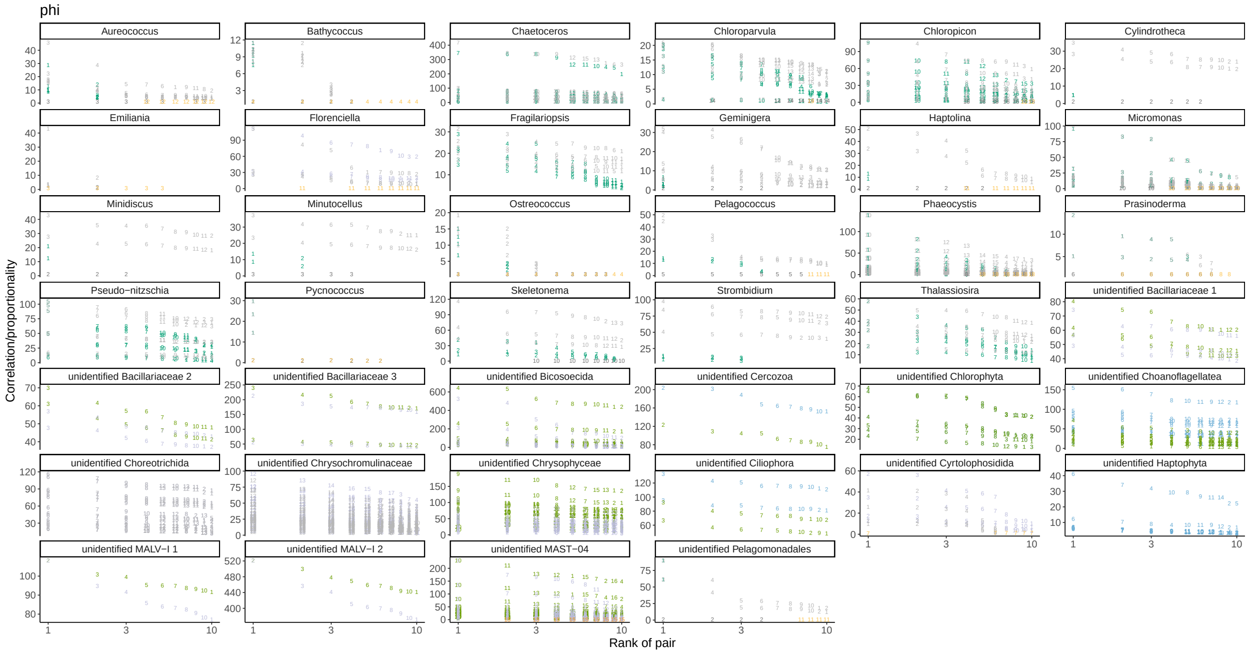


Figure S4: c

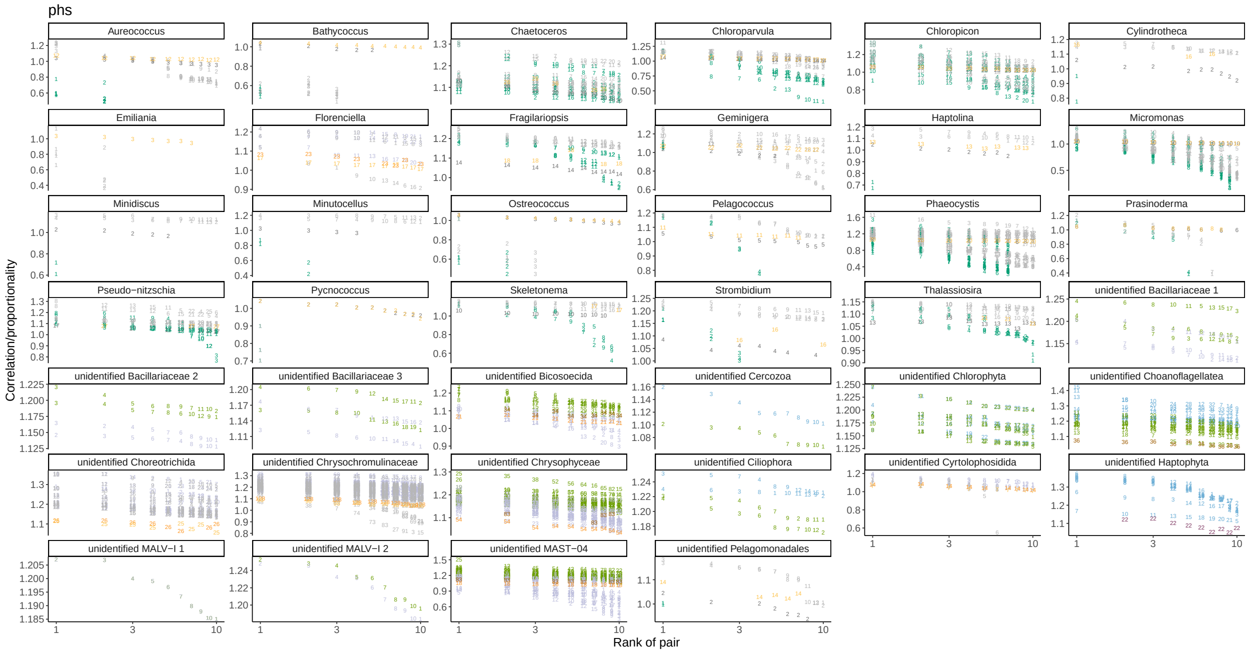


Figure S4: d

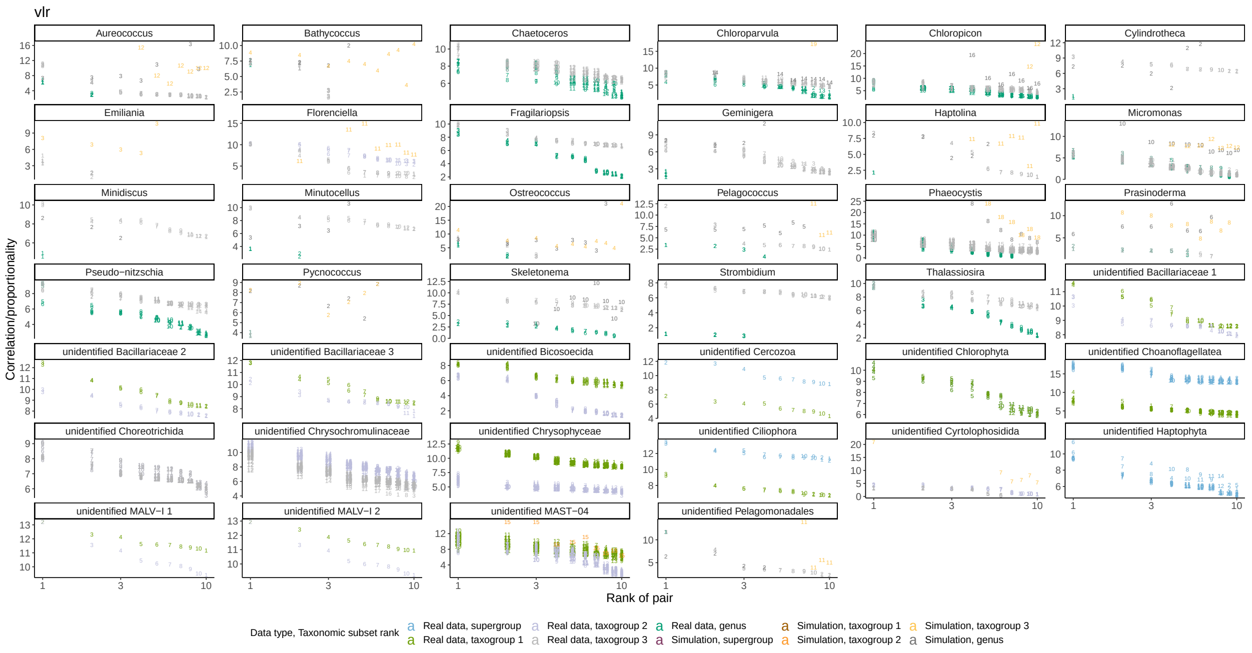


Figure S4: e

Figure S4: **Redundant V9 OTU matches.** V9 OTUs are represented by numbers in each plot, and can occur multiple times, with the same V9 OTUs represented by the same number. The correspondence estimates for V9 OTUs paired with SMAGs are shown on the vertical axis, and the rank of the match is on the horizontal axis. Colour represents the taxonomic subset and whether the matches belong to the real data or to simulations. Each subfigure represents statistics for different correspondence estimation metrics, the metrics indicated in the header of the subfigure: (a) *rho* proportionality, (b) Spearman's correlation, (c) *phi* proportionality, (d) *phs* proportionality, (e) *vlr* proportionality.

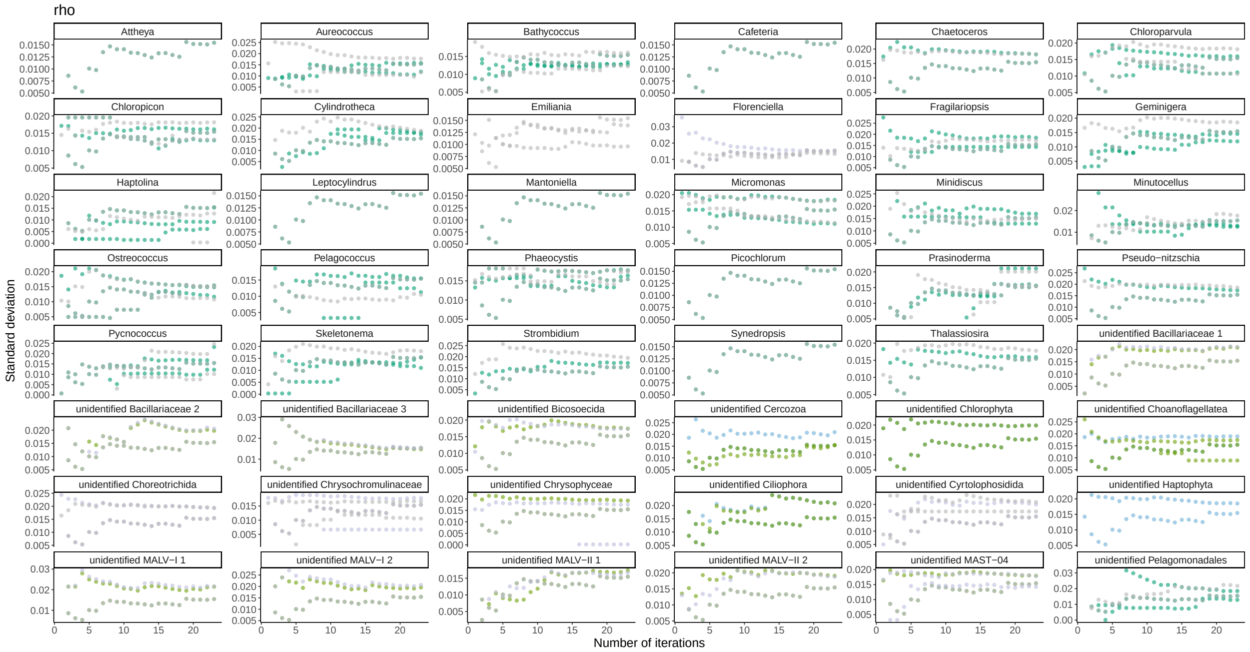


Figure S5: a

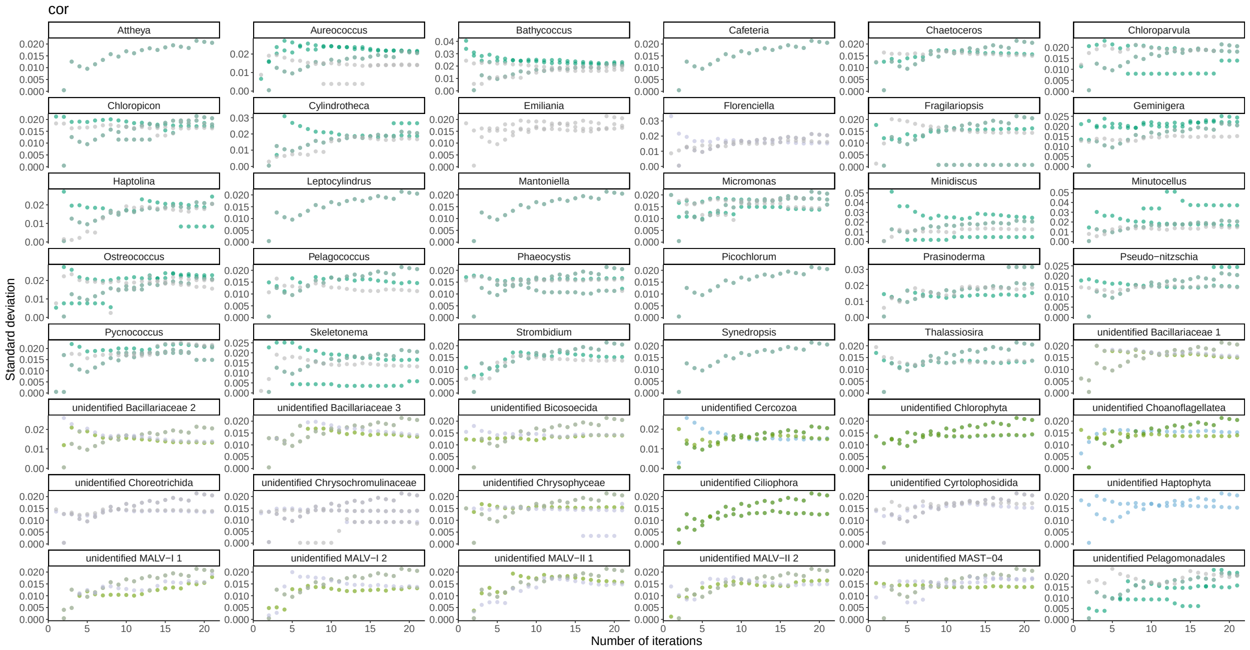


Figure S5: b

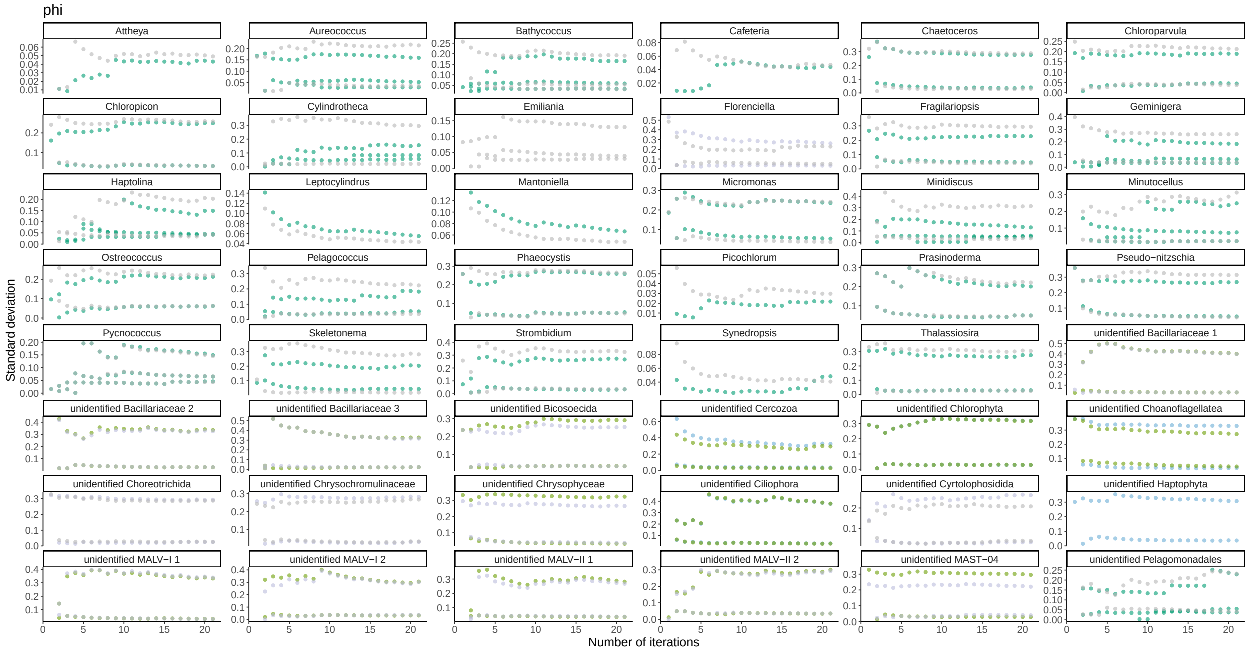


Figure S5: c

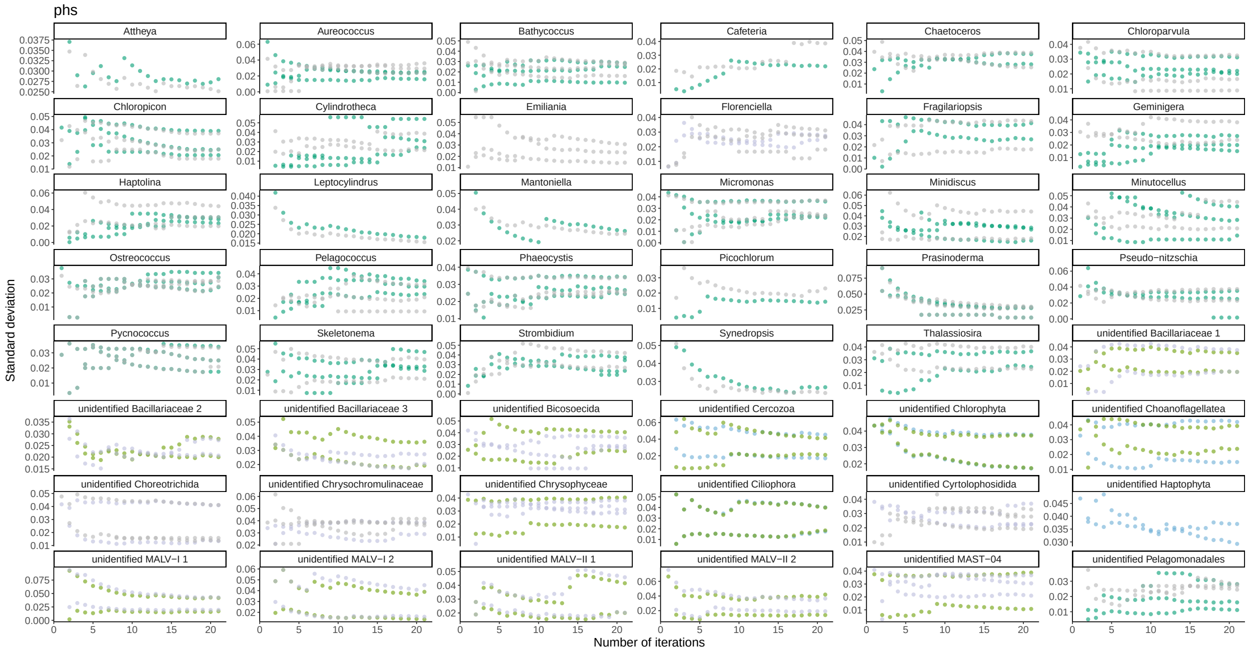


Figure S5: d

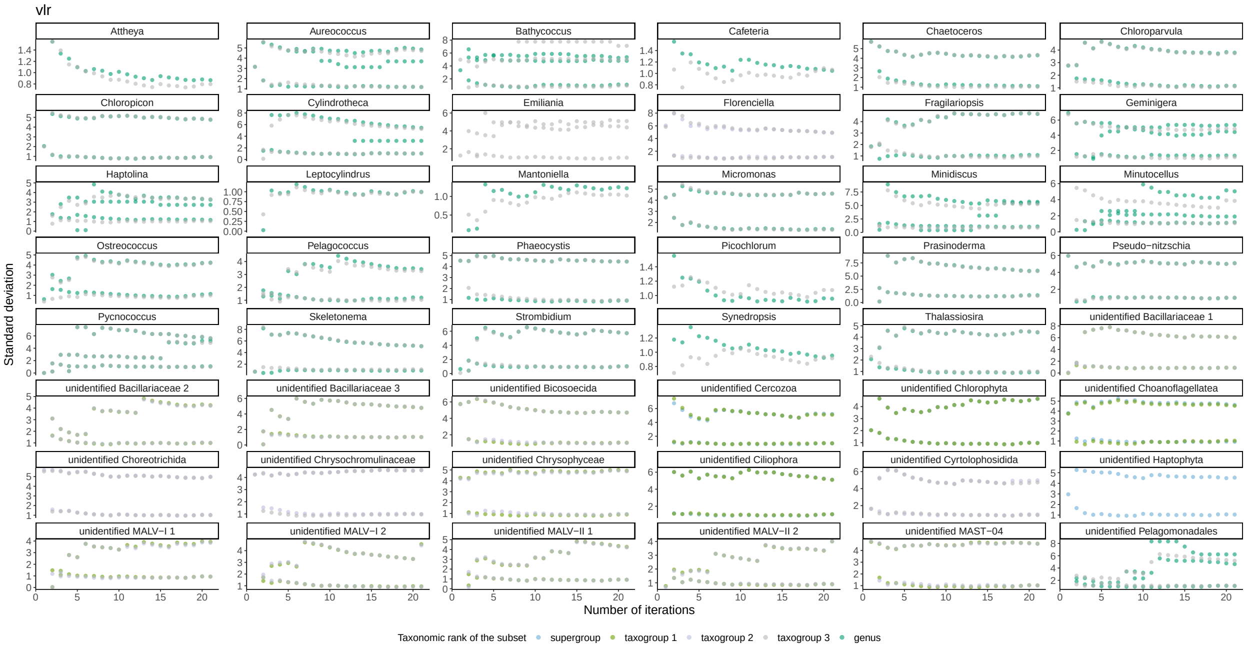


Figure S5: e

Figure S5: **Change of the standard deviation of the correspondence estimates (vertical axis) of simulated datasets depending on the number of simulation replicates (horizontal axis).** Standard deviation is shown for the correspondence value of the top-1st match of a positive control SMAG with the positive control V9 OTU. Colour corresponds to the taxonomic subset shown. Each subfigure represents statistics for different correspondence estimation metrics, with the metrics indicated in the header of the subfigure: (a) *rho* proportionality, (b) Spearman's correlation, (c) *phi* proportionality, (d) *phs* proportionality, (e) *vlr* proportionality.

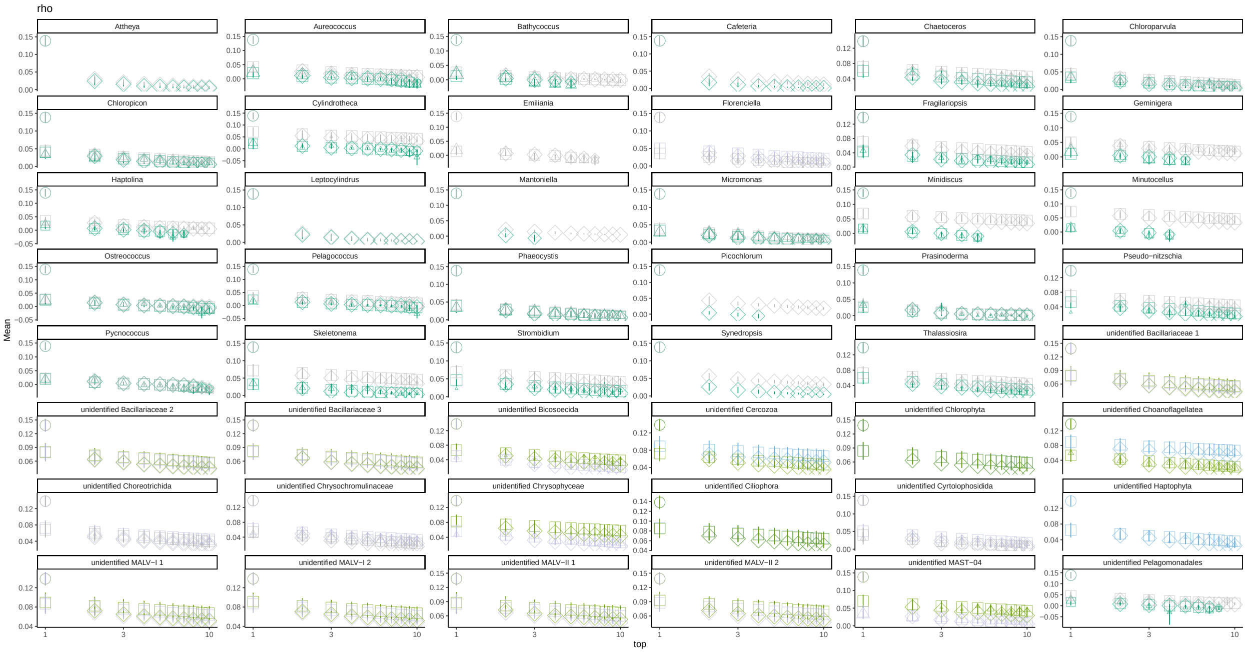


Figure S6: a

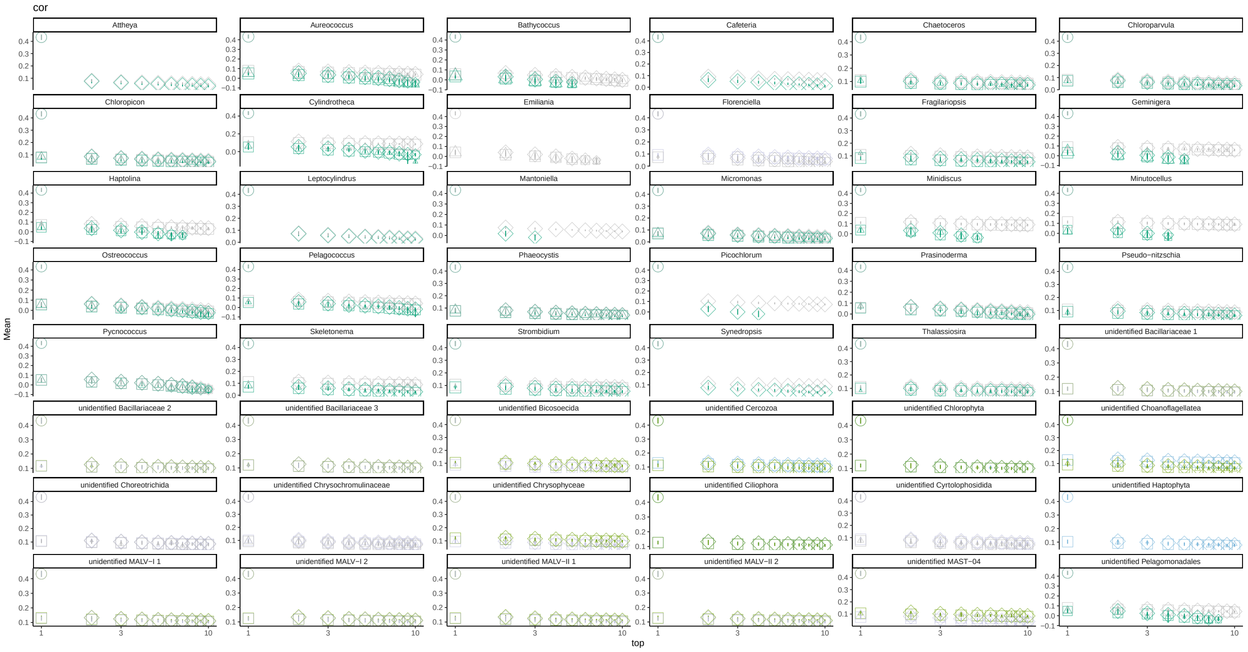


Figure S6: b

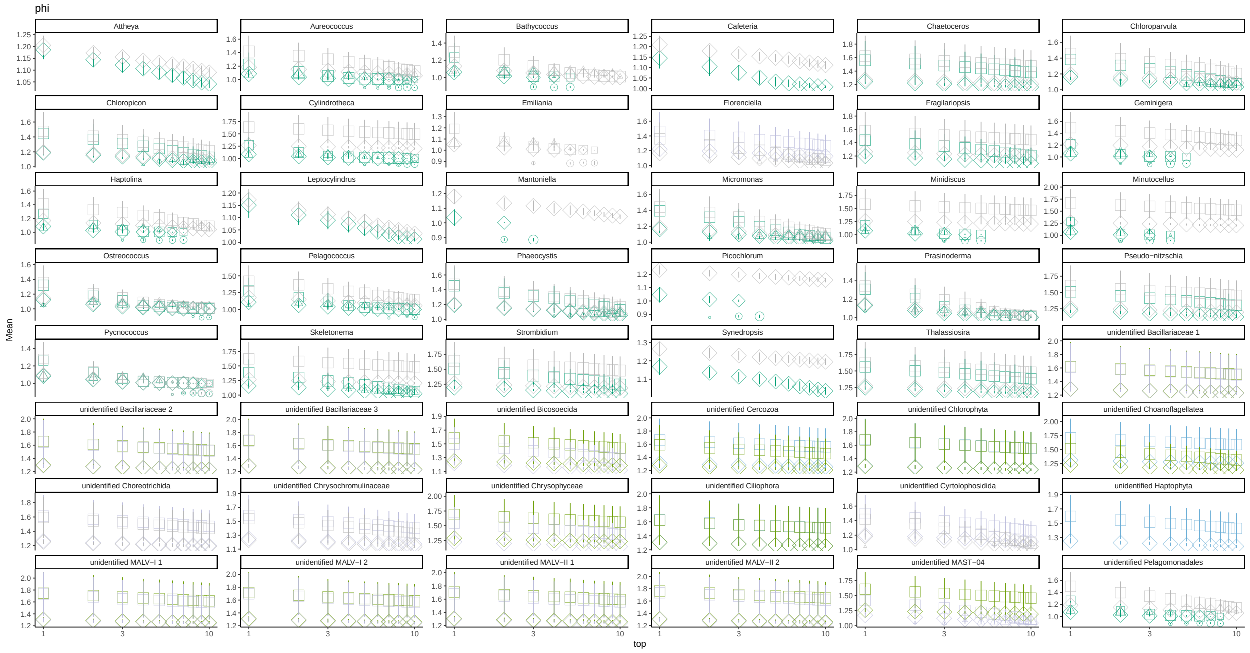


Figure S6: c

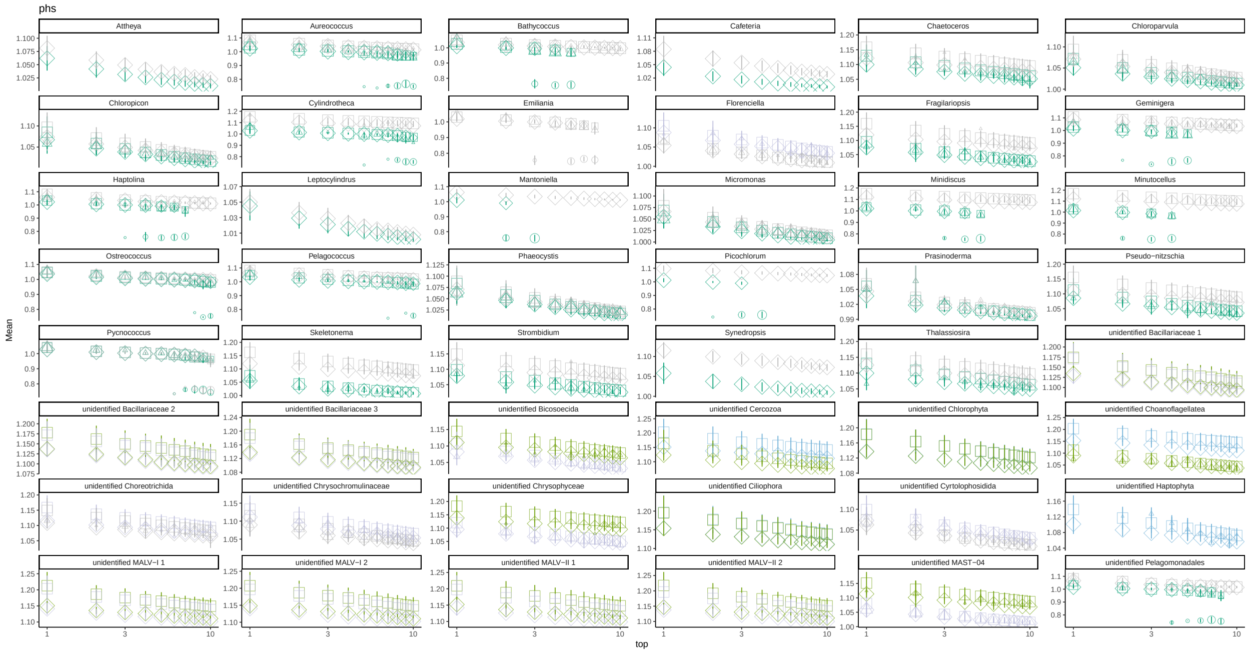


Figure S6: d

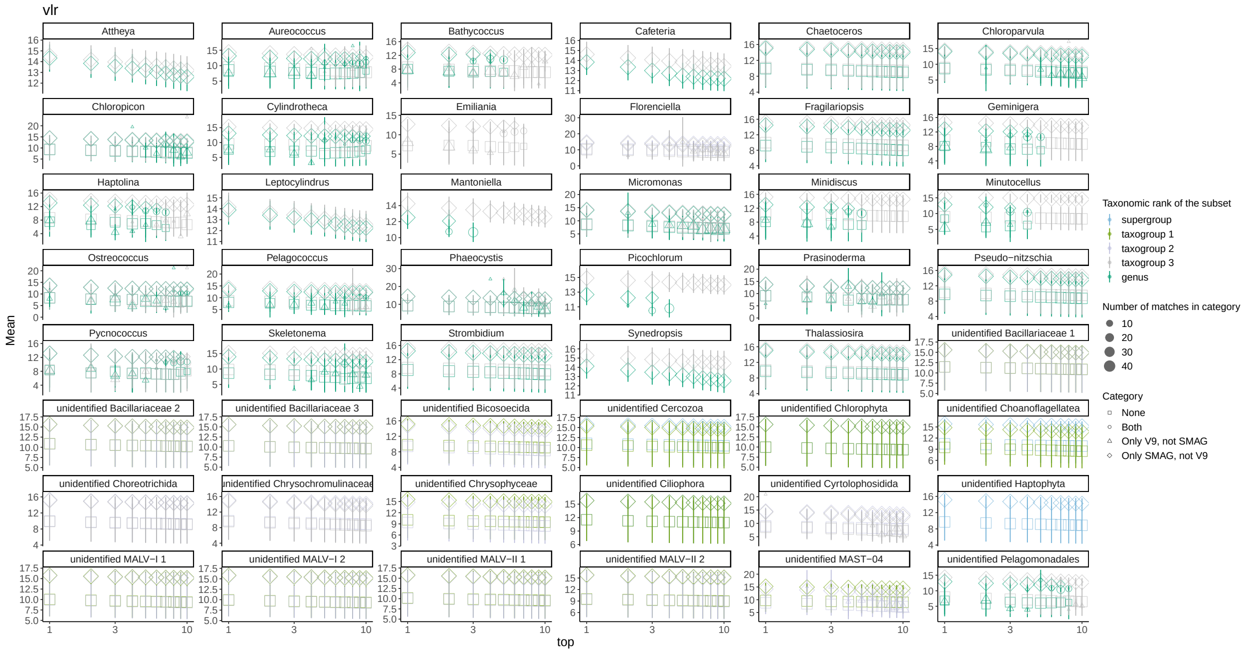


Figure S6: e

Figure S6: **Frequency of different match categories obtained from proportionality calculations for simulated datasets.** Proportionality values of V9 OTUs paired with SMAGs are shown on the vertical axis, and the rank of the match on the horizontal axis (log scale). The shape of the dots corresponds to the match category it represents (see Section S2.2). The size of the points reflects the number of replicates for which a given type of match was recovered; the error bars show the standard deviation of the mean correspondence estimates of the given type of match of the given rank. The colour represents the taxonomic subset shown. Each subfigure represents statistics for different correspondence estimation metrics, with the metrics indicated in the header of the subfigure: (a) *rho* proportionality, (b) Spearman's correlation, (c) *phi* proportionality, (d) *phs* proportionality, (e) *vlr* proportionality.

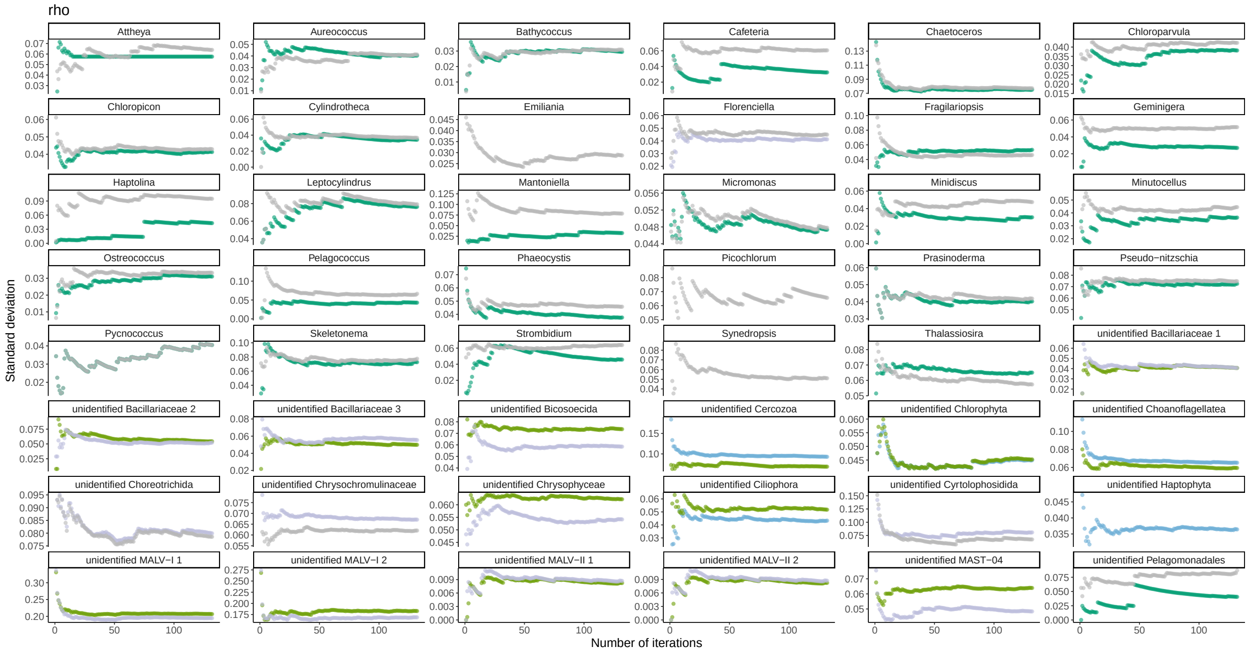


Figure S7: a

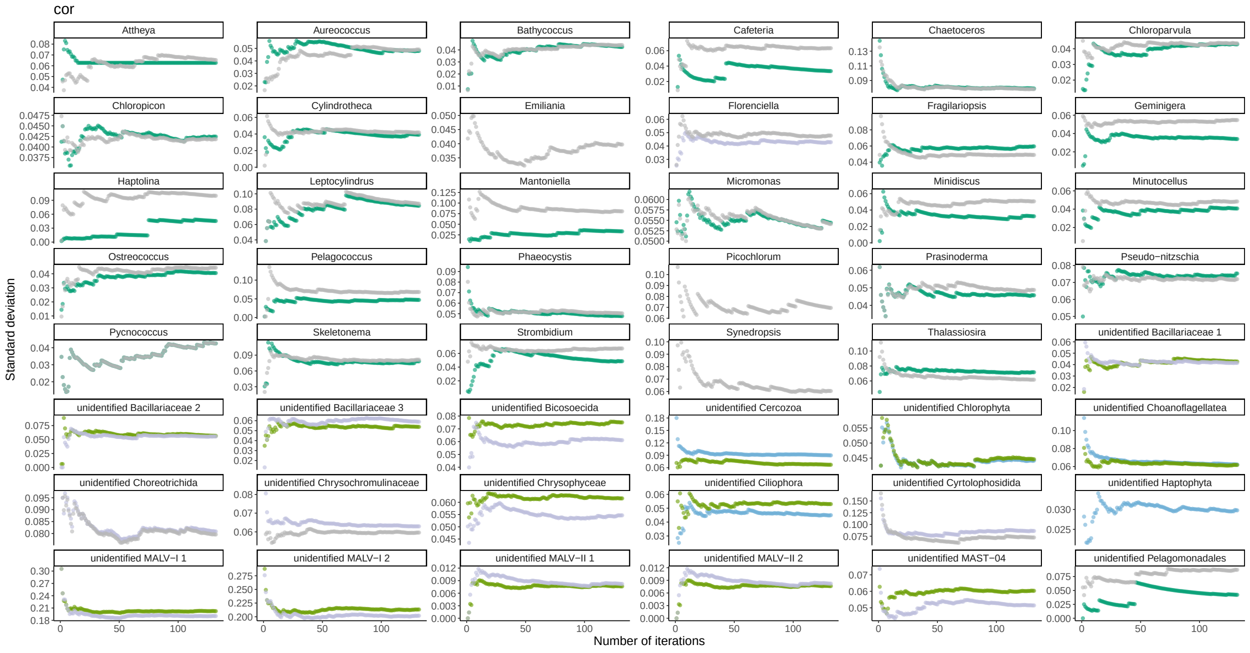


Figure S7: b

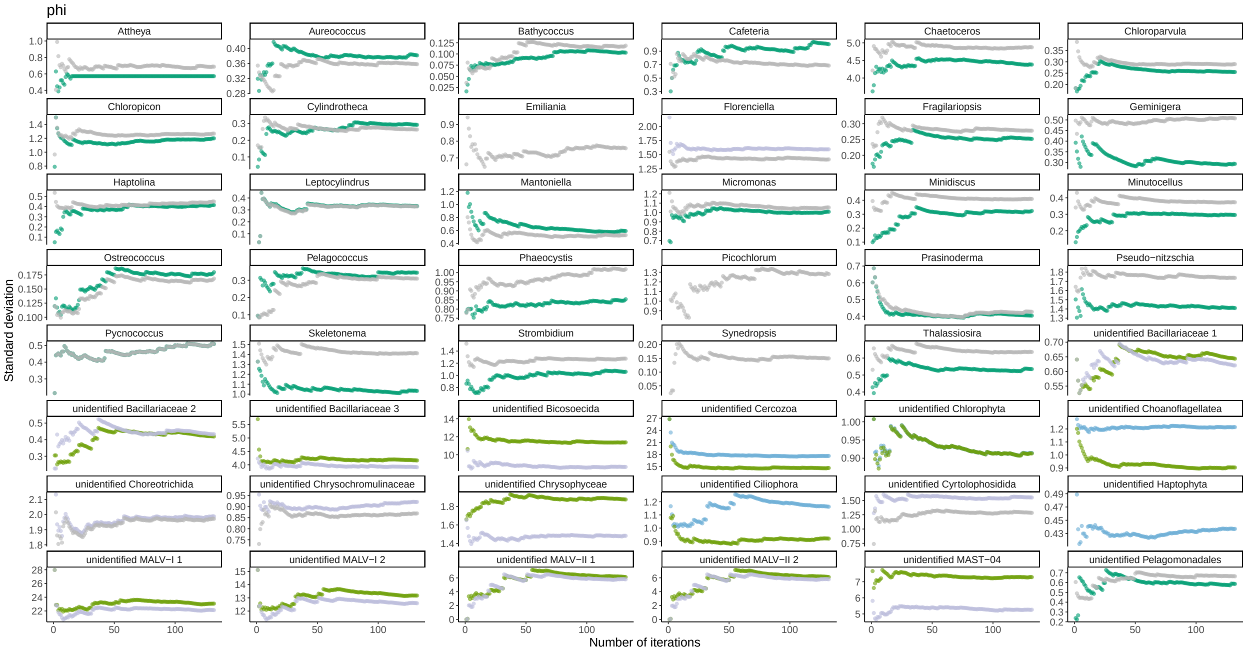


Figure S7: c

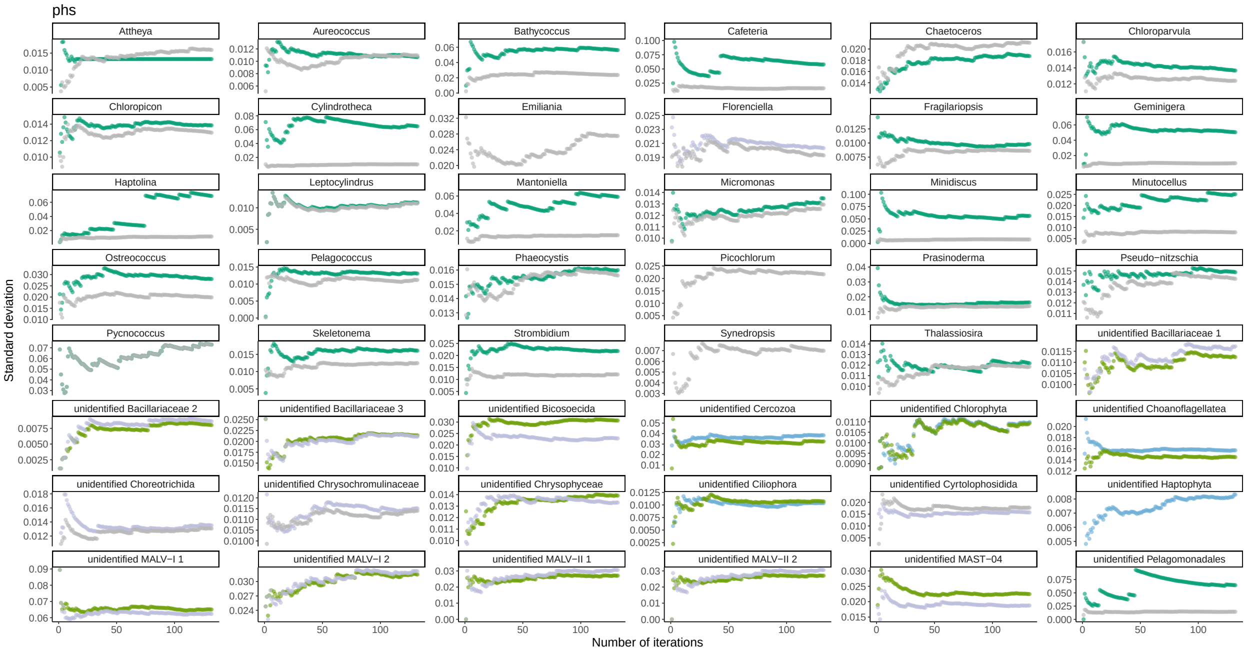


Figure S7: d

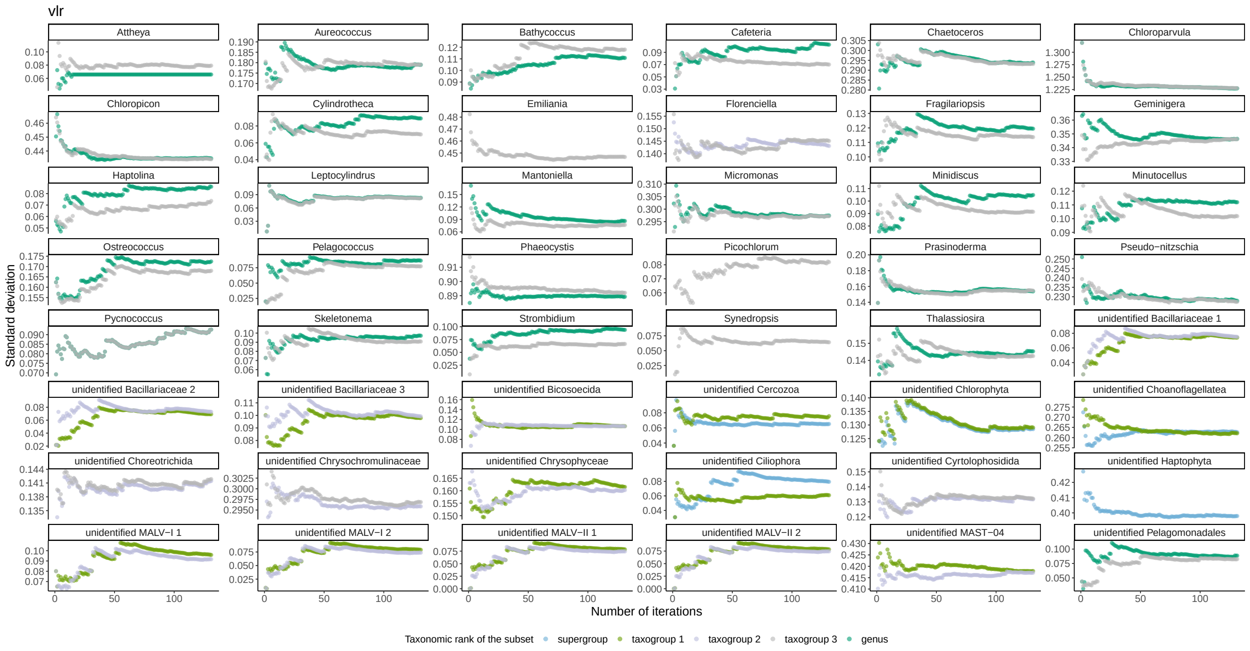

Figure S7: e

Figure S7: **Change of the standard deviation of the correspondence estimates (vertical axis) depending on the number of replicates of the shuffling experiment (horizontal axis).** Standard deviation is shown for the correspondence estimates of the top-1 best-ranked match. Colour corresponds to the taxonomic rank of the subset shown. Each subfigure represents statistics for different correspondence estimation metrics, with the metrics indicated in the header of the subfigure: (a) *rho* proportionality, (b) Spearman's correlation, (c) *phi* proportionality, (d) *phs* proportionality, (e) *vlr* proportionality.

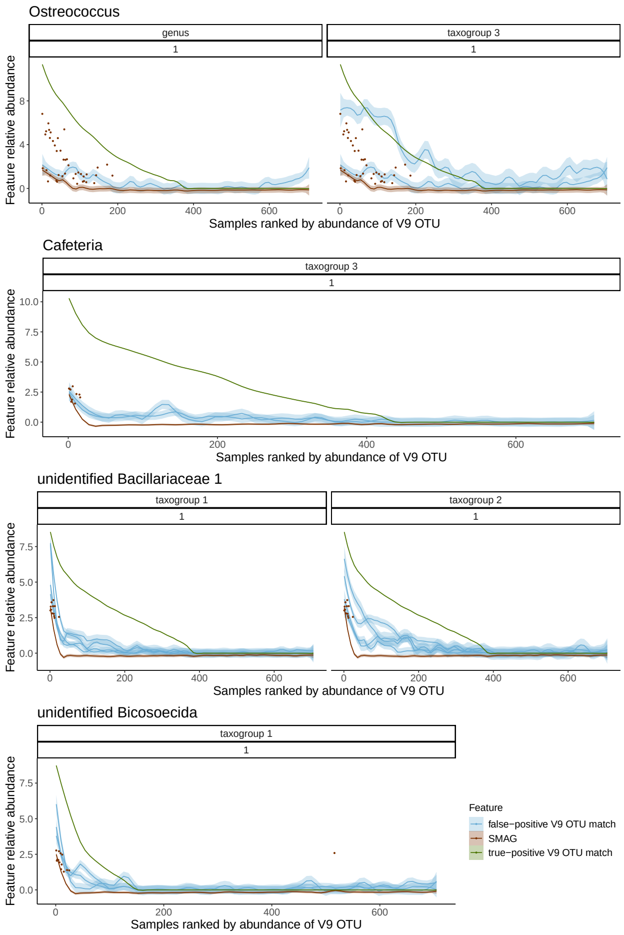


Figure S8: **Relative abundances across stations for control SMAGs, V9 OTUs present in control SMAGs, and abundances of V9 OTUs that showed the highest proportionality/correlation with control SMAGs (false positives, according to different metrics).** Each line represents a smoothened relative abundance distribution of a single feature (SMAG or V9 OTU). Samples on each plot are ranked based on the abundance of the control V9 OTU. Dots highlight SMAG abundance which is >0. Only matches from subsets of the two lowest taxonomic ranks are shown.

| Taxonomic subset name | SMAG taxonomy "genre" | Count | V9 "supergroup" | Count | V9 "taxogroup_1" | Count | V9 "taxogroup_2" | Count | V9 "taxogroup_3" | Count | V9 "genus" | Count |
|---|---|---|---|---|---|---|---|---|---|---|---|---|
| Attheya | Attheya | 1 | Stramenopiles | 10648 | Ochrophyta | 6926 | Diatomeae | 5430 | Biddulphiophyceae | 128 | Attheya | 60 |
| Aureococcus | Aureococcus | 6 | Stramenopiles | 10648 | Ochrophyta | 6926 | Pelagophyceae | 119 | Pelagomonadales | 84 | Aureococcus | 11 |
| Bathycoccus | Bathycoccus | 8 | Chloroplastida | 7436 | Chlorophyta | 7089 | Mamiellophyceae | 633 | Bathycoccaceae | 19 | Bathycoccus | 5 |
| Cafeteria | Cafeteria | 1 | Stramenopiles | 10648 | Opalozoa | 1228 | Bicosoecida | 230 | core-bicosoecids | 205 | Cafeteria | 26 |
| Chaetoceros | Chaetoceros | 11 | Stramenopiles | 10648 | Ochrophyta | 6926 | Diatomeae | 5430 | Mediophyceae | 2325 | Chaetoceros | 632 |
| Chloroparvula | chloroparvula | 7 | Chloroplastida | 7436 | Chlorophyta | 7089 | Chloropicophyceae | 138 | Chloropicophyceae | 138 | Chloroparvula | 58 |
| Chloropicon | Chloropicon | 11 | Chloroplastida | 7436 | Chlorophyta | 7089 | Chloropicophyceae | 138 | Chloropicophyceae | 138 | Chloropicon | 80 |
| Cylindrotheca | Cylindrotheca | 2 | Stramenopiles | 10648 | Ochrophyta | 6926 | Diatomeae | 5430 | Bacillariophyceae | 1929 | Cylindrotheca | 11 |
| Emiliania | Emiliania | 5 | Haptophyta | 953 | Prymnesiophyceae | 746 | Prymnesiophyceae | 746 | Isochrysidales | 7 | Emiliania | 0 |
| Florenciella | Florenciella | 3 | Stramenopiles | 10648 | Ochrophyta | 6926 | Dictyochophyceae | 269 | Florenciellales | 57 | NA | 0 |
| Fragilariopsis | Fragilariopsis | 5 | Stramenopiles | 10648 | Ochrophyta | 6926 | Diatomeae | 5430 | Bacillariophyceae | 1929 | Fragilariopsis | 135 |
| Geminigera | Geminigera | 7 | Cryptophyceae | 343 | Cryptomonadales | 439 | Cryptomonadales | 439 | lineage-B | 239 | Geminigera | 5 |
| Haptolina | Haptolina | 2 | Haptophyta | 953 | Prymnesiophyceae | 746 | Prymnesiophyceae | 746 | Prymnesiaceae | 59 | Haptolina | 7 |
| Leptocylindrus | Leptocylindrus | 1 | Stramenopiles | 10648 | Ochrophyta | 6926 | Diatomeae | 5430 | Leptocylindrophyceae | 53 | Leptocylindrus | 35 |
| Mantoniella | Mantoniella | 1 | Chloroplastida | 7436 | Chlorophyta | 7089 | Mamiellophyceae | 633 | Mamiellaceae | 68 | Mantoniella | 3 |
| Micromonas | Micromonas | 20 | Chloroplastida | 7436 | Chlorophyta | 7089 | Mamiellophyceae | 633 | Mamiellaceae | 68 | Micromonas | 44 |
| Minidiscus | Minidiscus | 2 | Stramenopiles | 10648 | Ochrophyta | 6926 | Diatomeae | 5430 | Mediophyceae | 2325 | Minidiscus | 5 |
| Minutocellus | Minutocellus | 2 | Stramenopiles | 10648 | Ochrophyta | 6926 | Diatomeae | 5430 | Mediophyceae | 2325 | Minutocellus | 4 |
| Ostreococcus | Ostreococcus | 4 | Chloroplastida | 7436 | Chlorophyta | 7089 | Mamiellophyceae | 633 | Bathycoccaceae | 19 | Ostreococcus | 14 |
| Pelagococcus | Pelagococcus | 2 | Stramenopiles | 10648 | Ochrophyta | 6926 | Pelagophyceae | 119 | Pelagomonadales | 84 | Pelagococcus | 14 |
| Phaeocystis | Phaeocystis | 27 | Haptophyta | 953 | Prymnesiophyceae | 746 | Prymnesiophyceae | 746 | Phaeocystales | 104 | Phaeocystis | 80 |
| Picochlorum | Picochlorum | 1 | Chloroplastida | 7436 | Chlorophyta | 7089 | Trebouxiophyceae | 879 | Chlorellales | 524 | Picochlorum | 4 |
| Prasinoderma | Prasinoderma | 2 | Chloroplastida | 7436 | Palmophyllophyceae | 37 | Prasinoderma-clade | 31 | Prasinoderma-clade | 31 | Prasinoderma | 21 |
| Pseudo-nitzschia | Pseudo-nitzschia | 7 | Stramenopiles | 10648 | Ochrophyta | 6926 | Diatomeae | 5430 | Bacillariophyceae | 1929 | Pseudo-nitzschia | 275 |
| Pycnococcus | Pycnococcus | 3 | Chloroplastida | 7436 | Chlorophyta | 7089 | Pycnococcaceae | 12 | Pycnococcaceae | 12 | Pycnococcus | 12 |
| Skeletonema | Skeletonema | 3 | Stramenopiles | 10648 | Ochrophyta | 6926 | Diatomeae | 5430 | Mediophyceae | 2325 | Skeletonema | 45 |
| Strombidium | Strombidium | 3 | Alveolata | 78782** | Ciliophora | 18850 | Spirotrichea | 2121 | Oligotrichia | 1361 | Sapolatum-clade | 155 |
| Synedropsis | Synedropsis | 1 | Stramenopiles | 10648 | Ochrophyta | 6926 | Diatomeae | 5430 | Bacillariophyceae | 1929 | Synedropsis | 60 |
| Thalassiosira | Thalassiosira | 5 | Stramenopiles | 10648 | Ochrophyta | 6926 | Diatomeae | 5430 | Mediophyceae | 2325 | Thalassiosira | 593 |
| unidentified Bacillariaceae 1 | New_Bacillariaceae_01 | 3 | Stramenopiles | 10648 | Ochrophyta | 6926 | Diatomeae | 5430 | NA | 0 | NA | 0 |
| unidentified Bacillariaceae 2 | New_Bacillariaceae_02 | 2 | Stramenopiles | 10648 | Ochrophyta | 6926 | Diatomeae | 5430 | NA | 0 | NA | 0 |
| unidentified Bacillariaceae 3 | Unidentified_Bacillariaceae | 2 | Stramenopiles | 10648 | Ochrophyta | 6926 | Diatomeae | 5430 | NA | 0 | NA | 0 |
| unidentified Bicosoecida | New_Bicosoecales_01 | 8 | Stramenopiles | 10648 | Opalozoa | 1228 | Bicosoecida | 230 | NA | 0 | NA | 0 |
| unidentified Cercozoa | Unidentified_Cercozoa | 2 | Rhizaria | 19891 | Cercozoa | 2625 | NA | 0 | NA | 0 | NA | 0 |
| unidentified Chlorophyta | New_Chlorophyta_01 | 6 | Chloroplastida | 7436 | Chlorophyta | 7089 | NA | 0 | NA | 0 | NA | 0 |
| unidentified Choanoflagellatea | New_Choanozoa_01 | 11 | Opisthokonta | 33545** | Choanoflagellata | 298 | NA | 0 | NA | 0 | NA | 0 |
| unidentified Choreotrichida | New_Choreotrichida _01 | 15 | Alveolata | 78782** | Ciliophora | 18850 | Spirotrichea | 2121 | Oligotrichia | 1361 | NA | 0 |
| unidentified Chrysochromulinaceae | New_Chrysochromulinaceae_01 | 39 | Haptophyta | 953 | Prymnesiophyceae | 746 | Prymnesiophyceae | 746 | Chrysochromulinaceae | 333 | NA | 0 |
| unidentified Chrysophyceae | New_Chrysophyceae | 23 | Stramenopiles | 10648 | Ochrophyta | 6926 | Chrysophyceae | 397 | NA | 0 | NA | 0 |
| unidentified Ciliophora | New_Ciliophora_01 | 2 | Alveolata | 78782** | Ciliophora | 18850 | NA | 0 | NA | 0 | NA | 0 |
| unidentified Cyrtolophosidida | New_Cyrtolophosidida_01 | 4 | Alveolata | 78782** | Ciliophora | 18850 | Colpodea | 182 | Cyrtolophosidida | 69 | NA | 0 |
| unidentified Haptophyta | New_Haptophyta_01 | 10 | Haptophyta | 953 | NA | 0 | NA | 0 | NA | 0 | NA | 0 |
| unidentified MALV-I 1 | New_MALV-I_01 | 2 | Alveolata | 78782** | Dinoflagellata | 55005** | MALV-I | 11911 | NA | 0 | NA | 0 |
| unidentified MALV-I 2 | MALV-I | 2 | Alveolata | 78782** | Dinoflagellata | 55005** | MALV-I | 11911 | NA | 0 | NA | 0 |
| unidentified MALV-II 1 | New_MALV-II_01 | 2 | Alveolata | 78782** | Dinoflagellata | 55005** | MALV-II | 9698 | NA | 0 | NA | 0 |
| unidentified MALV-II 2 | MALV-II | 3 | Alveolata | 78782** | Dinoflagellata | 55005** | MALV-II | 9698 | NA | 0 | NA | 0 |
| unidentified MAST-04 | New_MAST-4 | 25 | Stramenopiles | 10648 | Sagenista | 1655 | MAST-04 | 68 | NA | 0 | NA | 0 |
| unidentified Pelagomonadales | Sister_Pelagomonas | 2 | Stramenopiles | 10648 | Ochrophyta | 6926 | Pelagophyceae | 119 | Pelagomonadales | 84 | Pelagomonas | 8 |

Table S1: The "Taxonomic subset" column contains names of taxonomic subsets as shown on the figures, in correspondence with SMAG and V9 OTU classifications they represent. SMAG taxonomy "genre" column represents SMAG taxonomic assignment, identical to the one used in SMAG abundance dataset (Table S04 of (Delmont et al., 2022)), with the only exceptions being "Unidentified_Bacillariaceae" and "Unidentified_Cercozoa" groups. Those were artificially assigned to SAGs that were used as a control (see SMAGs Control) and had no initial taxonomic assignment in the original abundance table, but were assigned to a higher-rank taxonomic group according to Table S03 of (Delmont et al., 2022). Columns V9 "supergroup", V9 "taxogroup 1", V9 "taxogroup 2", V9 "taxogroup 3", and V9 "genus" show the classification of V9 OTUs at 5 different ranks matching SMAGs taxonomic assignment. Counts columns represent the number of SMAGs or V9 OTUs within a given taxonomic group: either SMAGs with total raw abundance >0 or V9 OTUs with total raw abundance across all samples > 100; the same sizes of taxonomic subsets were used in shuffling S3.3 and simulations S2.2, except in the case of of taxonomic subsets with >20000 V9 OTUs, which were artificially reduced to 20000 due to computational burden; those are indicated by **.

| Genome taxonomy | Genome/SMAG ID | GenBank18S ID | Scaffold with 18S | Tara V9 barcode | Tara V9 blast % identity |
|---|---|---|---|---|---|
| Cafeteria roenbergensis | VLTN01000011.1 / TARA_ION_45_MAG_00147 | MN334557 | GCA_008330645.1_CrBVI_genomic | dedd73012459fedb2d854754f2c4acf248d1712d | 100 |
| Pycnococcus provasolii | OW568862.1 / TARA_AON_82_MAG_00189 | X91264 | GCA_938743325.1_ucPycProv1.1_genomic | e637558ec51a7fe0a2cb11d338d2079bc7d74786 | 100 |
| Ostreococcus lucimarinus | NC_009366.1 / TARA_AON_82_MAG_00012 | Y15814 | GCF_000092065.1_ASM9206v1_genomic | 6b514fe39086fcf7c1325878394f32b5e6141e95 | 100 |
| Micromonas pusilla | NW_003315882.1 / TARA_AON_82_MAG_00118 | KT860843 | GCF_000151265.2_Micromonas_pusilla_CCMP1545_v2.0_genomic | 495f75dd678a6700fd8f3fc3fab2aa87800d0aa1 | 100 |
| unidentified Chrysophyceae | Metagenome_centric_SAG_TOSAG00_8 | Z38025 | Metagenome_centric_SAG_TOSAG00_8_scaffold23 | 920df2f681bb3553916becceea25586702322ecd | 99.231 |
| unidentified Bicosoecida_01 | Metagenome_centric_SAG_TOSAG00_9 | EF023971 | Metagenome_centric_SAG_TOSAG00_9_scaffold260 | aca17e422ca891974c6af687ccf37be1e685b732 | 100 |
| unidentified Chrysophyceae | Metagenome_centric_SAG_TOSAG23_30 | Z38025 | Metagenome_centric_SAG_TOSAG23_30_scaffold1 | d945a94b464aed8e5322f7fd0a04bd30f6355957 | 100 |
| unidentified Bacillariaceae 3 | Metagenome_centric_SAG_TOSAG39_4 | Y10570 | Metagenome_centric_SAG_TOSAG39_4_scaffold39 | c9ed411fc2cb7c5b69ca7d739c193c9bad6af9bd | 100 |
| unidentified Chrysophyceae | Metagenome_centric_SAG_TOSAG41_5 | Z38025 | Metagenome_centric_SAG_TOSAG41_5_scaffold18 | 1eb8e3f6b2b77f89775fe68f7693c1deb0a1a299 | 100 |
| unidentified Bacillariaceae 1 | Metagenome_centric_SAG_TOSAG46_2 | Y10570 | Metagenome_centric_SAG_TOSAG46_2_scaffold29 | a642e178c5f32337ab5f4d18c97ae6cb32d93dd9 | 100 |

Table S2: A summary on reference genomes and SAGs used for the positive controls. Columns, from left to right, depict: taxonomic assignment of SAG/reference genomes (see also S4.4), genome IDs, GenBank IDs of 18S sequences used as a query for identifying 18S location within a genome, IDs of scaffolds that contain 18S sequences, the representative barcode of V9 OTUs from Tara dataset best matching 18S from the genome, % identity and e-value for genome 18S alignment vs Tara V9 OTU match.
