## Supplementary material for "On the relationship between protist metabarcoding and protist metagenome-assembled genomes": LaTeX source for the pdf: library.bib

@article {Schlegel2016,
	author = {Schlegel, Philipp and Texada, Michael J and Miroschnikow, Anton and Peters, Marc and Schneider-Mizell, Casey M and Lacin, Haluk and Li, Feng and Fetter, Richard D and Truman, James W and Cardona, Albert and Pankratz, Michael J},
	title = {Synaptic Transmission Parallels Neuromodulation in a Central Food-Intake Circuit},
	year = {2016},
	doi = {10.1101/044990},
	publisher = {Cold Spring Harbor Labs Journals},
	abstract = {Neuromedin U (NMU) is a potent regulator of food intake and activity in mammals. While some downstream targets of NMU have been localized, connectivity of the neural circuits employing this neuropeptide is largely unknown. In Drosophila, neurons producing the homologous neuropeptide hugin regulate feeding and locomotion in a similar manner and project to structures of the central nervous system analogous to those in which NMU is found. Here, we use EM reconstruction and receptor expression analysis to map the connectome of hugin-producing neurons in the Drosophila larval central nervous system. We show that hugin-producing neurons establish distinct units that are reciprocally connected and share connectivity motifs. One of these units employs fast synaptic transmission as well as neuromodulation to target neuroendocrine cells (NSCs) of the pars intercerebralis, the Drosophila homolog of the hypothalamus. These NSCs produce CRH- and insulin-like peptides, which are homologs of downstream targets of NMU. Furthermore, most of the hugin-producing neurons, including those that target the NSCs, receive inputs from chemosensory neurons in the subesophageal zone, the brain stem analog in Drosophila. Our data positions hugin neurons as part of a novel sensory-to-endocrine network that reflects the way NMU operates in mammals. We propose that the hugin neurons interconnecting chemosensory and neuroendocrine organs are part of a physiological control system that has been conserved not only at functional and molecular levels, but at the network architecture level as well.},
	URL = {http://biorxiv.org/content/early/2016/04/07/044990},
	eprint = {http://biorxiv.org/content/early/2016/04/07/044990.full.pdf},
	journal = {bioRxiv}
}
@article{Tragin2018,
   abstract = {We compared the composition of eukaryotic communities using two genetic markers (18S rRNA V4 and V9 regions) at 27 sites sampled during Ocean Sampling Day 2014, with a focus on photosynthetic groups and, more specifically green algae (Chloro-phyta). Globally, the V4 and V9 regions of the 18S rRNA gene provided similar images of alpha diversity and ecological patterns. However, V9 provided 20 more OTUs built at 97 identity than V4. 34 of the genera were found with both markers and, of the remnant , 22 were found only with V4 and 44 only with V9. For photosynthetic groups, V4 and V9 performed equally well to describe global communities at different taxonomic levels from the division to the genus and provided similar Chlorophyta distribution patterns. However, at lower taxonomic level, the V9 dataset failed for example to describe the diversity of Dolichomastigales (Chlorophyta, Mamiellophyceae) emphasizing the lack of V9 sequences for this group and the importance of the reference database for metabarcode analysis. We conclude that in order to address questions regarding specific groups (e.g., a given genus), it is necessary to choose the marker based not only on the genetic divergence within this group but also on the existence of reference sequences in databases.},
   author = {Margot Tragin and Adriana Zingone and Daniel Vaulot},
   doi = {10.1111/1462-2920.13952},
   title = {Comparison of coastal phytoplankton composition estimated from the V4 and V9 regions of the 18S rRNA gene with a focus on photosynthetic groups and especially Chlorophyta},
   year = {2018}
   url = {https://ami-journals.onlinelibrary.wiley.com/doi/10.1111/1462-2920.13952},
}
@article{Mukherjee2020,
   author = {Indranil Mukherjee and Michaela M Salcher and Adrian-S ¸tefan Andrei and Vinicius Silva Kavagutti and Tanja Shabarova and Vesna Grujči and Markus Haber and Paul Layoun and Yoshikuni Hodoki and Shin-Ichi Nakano and Karel Šimek and Rohit Ghai},
   doi = {10.1111/1462-2920.15209},
   title = {A freshwater radiation of diplonemids},
   url = {https://ami-journals.onlinelibrary.wiley.com/doi/10.1111/1462-2920.15209},
   year = {2020},
}
@article{Decelle2014,
   abstract = {Metabarcoding is a powerful tool for exploring microbial diversity in the environment, but its accurate interpretation is impeded by diverse technical (e.g. PCR and sequencing errors) and biological biases (e.g. intra-individual polymorphism) that remain poorly understood. To help interpret environmental metabarcoding datasets, we investigated the intracellular diversity of the V4 and V9 regions of the 18S rRNA gene from Acantharia and Nassellaria (radiolarians) using 454 pyrosequencing. Individual cells of radiolarians were isolated, and PCRs were performed with generalist primers to amplify the V4 and V9 regions. Different denoising procedures were employed to filter the pyrosequenced raw amplicons (Acacia, AmpliconNoise, Linkage method). For each of the six isolated cells, an average of 541 V4 and 562 V9 amplicons assigned to radiolarians were obtained, from which one numerically dominant sequence and several minor variants were found. At the 97 identity, a diversity metrics commonly used in environmental surveys, up to 5 distinct OTUs were detected in a single cell. However, most amplicons grouped within a single OTU whereas other OTUs contained very few amplicons. Different analytical methods provided evidence that most minor variants forming different OTUs correspond to PCR and sequencing artifacts. Duplicate PCR and sequencing from the same DNA extract of a single cell had only 9 to 16 of unique amplicons in common, and alignment visualization of V4 and V9 amplicons showed that most minor variants contained substitutions in highly-conserved regions. We conclude that intracellular variability of the 18S rRNA in radiolarians is very limited despite its multi-copy nature and the existence of multiple nuclei in these protists. Our study recommends some technical guidelines to conservatively discard artificial amplicons from metabarcoding datasets, and thus properly assess the diversity and richness of protists in the environment.},
   author = {Johan Decelle and Sarah Romac and Eriko Sasaki and Fabrice Not and Frédéric Mahé},
   doi = {10.1371/JOURNAL.PONE.0104297},
   issn = {1932-6203},
   issue = {8},
   journal = {PLOS ONE},
   keywords = {Acacia,DNA extraction,DNA replication,DNA sequencing,Eukaryota,Polymerase chain reaction,Protists,Ribosomal RNA},
   month = {8},
   pages = {e104297},
   pmid = {25090095},
   publisher = {Public Library of Science},
   title = {Intracellular Diversity of the V4 and V9 Regions of the 18S rRNA in Marine Protists (Radiolarians) Assessed by High-Throughput Sequencing},
   volume = {9},
   url = {https://journals.plos.org/plosone/article?id=10.1371/journal.pone.0104297},
   year = {2014},
}
@article{Flynn2015,
   abstract = {Metabarcoding has the potential to become a rapid, sensitive, and effective approach for identifying species in complex environmental samples. Accurate molecular identification of species depends on the ability to generate operational taxonomic units (OTUs) that correspond to biological species. Due to the sometimes enormous estimates of biodiversity using this method, there is a great need to test the efficacy of data analysis methods used to derive OTUs. Here, we evaluate the performance of various methods for clustering length variable 18S amplicons from complex samples into OTUs using a mock community and a natural community of zooplankton species. We compare analytic procedures consisting of a combination of (1) stringent and relaxed data filtering, (2) singleton sequences included and removed, (3) three commonly used clustering algorithms (mothur, UCLUST, and UPARSE), and (4) three methods of treating alignment gaps when calculating sequence divergence. Depending on the combination of methods used, the number of OTUs varied by nearly two orders of magnitude for the mock community (60-5068 OTUs) and three orders of magnitude for the natural community (22-22191 OTUs). The use of relaxed filtering and the inclusion of singletons greatly inflated OTU numbers without increasing the ability to recover species. Our results also suggest that the method used to treat gaps when calculating sequence divergence can have a great impact on the number of OTUs. Our findings are particularly relevant to studies that cover taxonomically diverse species and employ markers such as rRNA genes in which length variation is extensive.},
   author = {Jullien M. Flynn and Emily A. Brown and Frédéric J.J. Chain and Hugh J. Macisaac and Melania E. Cristescu},
   doi = {10.1002/ECE3.1497},
   issn = {20457758},
   issue = {11},
   journal = {Ecology and Evolution},
   keywords = {18S rRNA,Biodiversity,High-throughput sequencing,Metabarcoding,OTU,eDNA},
   month = {6},
   pages = {2252-2266},
   pmid = {26078860},
   publisher = {John Wiley and Sons Ltd},
   title = {Toward accurate molecular identification of species in complex environmental samples: Testing the performance of sequence filtering and clustering methods},
   volume = {5},
   year = {2015},
}
@article{martin2022towards,
  title={Towards quantitative metabarcoding of eukaryotic plankton: an approach to improve 18S rRNA gene copy number bias},
  author={Martin, Jon Lapeyra and Santi, Ioulia and Pitta, Paraskevi and John, Uwe and Gypens, Nathalie},
  journal={Metabarcoding and Metagenomics},
  volume={6},
  pages={e85794},
  year={2022},
  publisher={Pensoft Publishers}
}
@article{Becker2023,
   abstract = {The diversity of soil bacteria was analyzed via metabarcoding and metagenomic approaches using DNA samples isolated from the biocrusts of 12 different Arctic and Antarctic sites. For the metabarcoding approach, the V3-4 region of the 16S rRNA was targeted. Our results showed that nearly all operational taxonomic units (OTUs = taxa) found in metabarcoding analyses were recovered in metagenomic analyses. In contrast, metagenomics identified a large number of additional OTUs absent in metabarcoding analyses. In addition, we found huge differences in the abundance of OTUs between the two methods. The reasons for these differences seem to be (1) the higher sequencing depth in metagenomics studies, which allows the detection of low-abundance community members in metagenomics, and (2) bias of primer pairs used to amplify the targeted sequence in metabarcoding, which can change the community composition dramatically even at the lower taxonomic levels. We strongly recommend using only metagenomic approaches when establishing the taxonomic profiles of whole biological communities.},
   author = {Burkhard Becker and Ekaterina Pushkareva},
   doi = {10.3390/GENES14040812/S1},
   issn = {20734425},
   issue = {4},
   journal = {Genes},
   keywords = {metabarcoding,metagenomics,microbiomes,polar regions,terrestrial communities},
   month = {4},
   pages = {812},
   pmid = {37107570},
   publisher = {MDPI},
   title = {Metagenomics Provides a Deeper Assessment of the Diversity of Bacterial Communities in Polar Soils Than Metabarcoding},
   volume = {14},
   url = {https://www.mdpi.com/2073-4425/14/4/812/htm https://www.mdpi.com/2073-4425/14/4/812},
   year = {2023},
}
@article{Frioux2020,
   abstract = {Metagenomic sequencing of complete microbial communities has greatly enhanced our understanding of the taxonomic composition of microbiotas. This has led to breakthrough developments in bioinformatic disciplines such as assembly, gene clustering, metagenomic binning of species genomes and the discovery of an incredible, so far undiscovered, taxonomic diversity. However, functional annotations and estimating metabolic processes from single species – or communities – is still challenging. Earlier approaches relied mostly on inferring the presence of key enzymes for metabolic pathways in the whole metagenome, ignoring the genomic context of such enzymes, resulting in the ‘bag-of-genes’ approach to estimate functional capacities of microbiotas. Here, we review recent developments in metagenomic bioinformatics, with a special focus on emerging technologies to simulate and estimate metabolic information, that can be derived from metagenomic assembled genomes. Genome-scale metabolic models can be used to model the emergent properties of microbial consortia and whole communities, and the progress in this area is reviewed. While this subfield of metagenomics is still in its infancy, it is becoming evident that there is a dire need for further bioinformatic tools to address the complex combinatorial problems in modelling the metabolism of large communities as a ‘bag-of-genomes’.},
   author = {Clémence Frioux and Dipali Singh and Tamas Korcsmaros and Falk Hildebrand},
   doi = {10.1016/J.CSBJ.2020.06.028},
   issn = {2001-0370},
   journal = {Computational and Structural Biotechnology Journal},
   keywords = {Bioinformatics,Gene Functions,Metabolic modelling,Metagenomic-assembled genomes,Metagenomics,Microbiota,Systems biology},
   month = {1},
   pages = {1722-1734},
   publisher = {Elsevier},
   title = {From bag-of-genes to bag-of-genomes: metabolic modelling of communities in the era of metagenome-assembled genomes},
   volume = {18},
   year = {2020},
}

@article{armbrecht2021paleo,
  title={Paleo-diatom composition from Santa Barbara Basin deep-sea sediments: a comparison of 18S-V9 and diat-rbcL metabarcoding vs shotgun metagenomics},
  author={Armbrecht, Linda and Eisenhofer, Raphael and Utge, Jos{\'e} and Sibert, Elizabeth C and Rocha, Fabio and Ward, Ryan and Pierella Karlusich, Juan Jos{\'e} and Tirichine, Leila and Norris, Richard and Summers, Mindi and others},
  journal={ISME Communications},
  volume={1},
  number={1},
  pages={66},
  year={2021},
  publisher={Nature Publishing Group UK London}
}
@article{tedersoo2015shotgun,
  title={Shotgun metagenomes and multiple primer pair-barcode combinations of amplicons reveal biases in metabarcoding analyses of fungi},
  author={Tedersoo, Leho and Anslan, Sten and Bahram, Mohammad and P{\~o}lme, Sergei and Riit, Taavi and Liiv, Ingrid and K{\~o}ljalg, Urmas and Kisand, Veljo and Nilsson, Henrik and Hildebrand, Falk and others},
  journal={MycoKeys},
  volume={10},
  pages={1--43},
  year={2015},
  publisher={Pensoft Publishers}
}
@article{Sternes2017,
   abstract = {Wine is a complex beverage, comprising hundreds of metabolites produced through the action of yeasts and bacteria in fermenting grape must. Commercially, there is now a growing trend away from using wine yeast (Saccharomyces) starter cultures, toward the historic practice of uninoculated or "wild" fermentation, where the yeasts and bacteria associated with the grapes and/or winery perform the fermentation. It is the varied metabolic contributions of these numerous non-Saccharomyces species that are thought to impart complexity and desirable taste and aroma attributes to wild ferments in comparison to their inoculated counterparts. To map the microflora of spontaneous fermentation, metagenomic techniques were employed to characterize and monitor the progression of fungal species in 5 different wild fermentations. Both amplicon-based ribosomal DNA internal transcribed spacer (ITS) phylotyping and shotgun metagenomics were used to assess community structure across different stages of fermentation. While providing a sensitive and highly accurate means of characterizing the wine microbiome, the shotgun metagenomic data also uncovered a significant overabundance bias in the ITS phylotyping abundance estimations for the common non-Saccharomyces wine yeast genus Metschnikowia. By identifying biases such as that observed for Metschnikowia, abundance measurements from future ITS phylotyping datasets can be corrected to provide more accurate species representation. Ultimately, as more shotgun metagenomic and single-strain de novo assemblies for key wine species become available, the accuracy of both ITS-amplicon and shotgun studies will greatly increase, providing a powerful methodology for deciphering the influence of the microbial community on the wine flavor and aroma.},
   author = {Peter R. Sternes and Danna Lee and Dariusz R. Kutyna and Anthony R. Borneman},
   doi = {10.1093/GIGASCIENCE/GIX040},
   issn = {2047217X},
   issue = {7},
   journal = {GigaScience},
   keywords = {Metagenomics,Phylotyping,Wine,Yeast diversity},
   month = {7},
   pages = {1-10},
   pmid = {28595314},
   publisher = {Oxford Academic},
   title = {A combined meta-barcoding and shotgun metagenomic analysis of spontaneous wine fermentation},
   volume = {6},
   url = {https://academic.oup.com/gigascience/article/6/7/gix040/3862720},
   year = {2017},
}
@article{bact,
   abstract = {In food safety the detection of food contaminations with pathogenic microorganisms is a race against time and often outpaced by error-prone epidemiological approaches. For evidence-based outbreak investigations fast and reliable techniques and procedures are required to identify the source of infection. Metagenomics has the potential to become a powerful tool in the field of modern food safety, since it allows the detection, identification and characterization of a broad range of pathogens in a single experiment without pre-cultivation within a couple of days. Nevertheless, sample handling, sequencing and data analysis are challenging and can introduce errors and biases into the analysis. In order to evaluate the potential of metagenomics in food safety, we generated a mock community containing DNA of foodborne bacteria. Herewith, we compare the aptitude of the two prevalent approaches – 16S rDNA amplicon sequencing and whole genome shotgun sequencing – for the detection of foodborne bacteria using different parameters during sample preparation, sequencing and data analysis. 16S rDNA sequencing did not only result in high deviations from the expected sample composition on genus and species level, but more importantly lacked the detection of several pathogenic species. While shotgun sequencing is more suitable for species detection, abundance estimation, genome assembly and species characterization, the performance can vary depending on the library preparation kit, which was confirmed for a naturally Francisella tularensis contaminated game meat sample. The application of the Nextera XT DNA Library Preparation Kit for shotgun sequencing did not only result in lower reference genome recovery and coverage, but also in distortions of the mock community composition. For data analysis, we propose a publicly available workflow for pathogen detection and characterization and demonstrate its benefits on the usability of metagenomic sequencing in food safety by analyzing an authentic metagenomic sample.},
   author = {Josephine Grützke and Burkhard Malorny and Jens Andre Hammerl and Anne Busch and Simon H. Tausch and Herbert Tomaso and Carlus Deneke},
   doi = {10.3389/FMICB.2019.01805/BIBTEX},
   issn = {1664302X},
   issue = {AUG},
   journal = {Frontiers in Microbiology},
   keywords = {16S,Bioinformatics,Food safety,Francisella tularensis,Harmonisation,Metagenomics,Mock community,Shotgun},
   month = {8},
   pages = {1805},
   publisher = {Frontiers Media S.A.},
   title = {Fishing in the soup – pathogen detection in food safety using metabarcoding and metagenomic sequencing},
   volume = {10},
   year = {2019},
}
@article{Vannier2016,
   abstract = {Bathycoccus is a cosmopolitan green micro-alga belonging to the Mamiellophyceae, a class of picophytoplankton that contains important contributors to oceanic primary production. A single species of Bathycoccus has been described while the existence of two ecotypes has been proposed based on metagenomic data. A genome is available for one strain corresponding to the described phenotype. We report a second genome assembly obtained by a single cell genomics approach corresponding to the second ecotype. The two Bathycoccus genomes are divergent enough to be unambiguously distinguishable in whole DNA metagenomic data although they possess identical sequence of the 18S rRNA gene including in the V9 region. Analysis of 122 global ocean whole DNA metagenome samples from the Tara-Oceans expedition reveals that populations of Bathycoccus that were previously identified by 18S rRNA V9 metabarcodes are only composed of these two genomes. Bathycoccus is relatively abundant and widely distributed in nutrient rich waters. The two genomes rarely co-occur and occupy distinct oceanic niches in particular with respect to depth. Metatranscriptomic data provide evidence for gain or loss of highly expressed genes in some samples, suggesting that the gene repertoire is modulated by environmental conditions. Phytoplankton, comprising prokaryotes and eukaryotes, contribute to nearly half of the annual global primary production 1. Picocyanobacteria of the genera Prochlorococcus and Synechococcus dominate the prokaryotic component 2. However, small eukaryotes (picoeukaryotes; < 2 μ m) can be major contributors to primary production 3,4. In contrast to cyanobacteria, the phylogenetic diversity of eukaryotic phytoplankton is wide, with species belonging to virtually all photosynthetic protist groups 5. Among them, three genera of green algae belonging to the order Mamiellales (class Mamiellophyceae 6), Micromonas, Ostreococcus and Bathycoccus are particularly important ecologically because they are found in a wide variety of oceanic ecosystems, from the poles to the tropics 7-12. The cosmopolitan distribution of these genera raises the questions of their diversity and their adaptation to local environmental conditions. These genera exhibit genetic diversity: for example, there are at least three genetically different clades of Micromonas with different habitat preferences 12,13. One ecotype of Micromonas seems to be restricted to polar waters 8,14. Ostreococcus which is the smallest free-living eukaryotic cell known to date with a cell size of 0.8 μ m 15 can be differentiated into at least four clades. Two Ostreococcus species have been formerly described: O. tauri and O. mediterreanus 15,16. Among these Ostreococcus clades, different strains seem to be adapted to different light ranges 17. However, the ecological preferences of Ostreococcus strains are probably more complex, implying other environmental parameters such as nutrients and temperature 9. The genus Bathycoccus was initially isolated at 100 m from the deep chlorophyll maximum (DCM) in the Mediterranean Sea 18 and cells with the same morphology (body scales) had been reported previously from the Atlantic Ocean 19 .},
   author = {Thomas Vannier and Jade Leconte and Yoann Seeleuthner and Samuel Mondy and Eric Pelletier and Jean-Marc Aury and Colomban de Vargas and Michael Sieracki and Daniele Iudicone and Daniel Vaulot and Patrick Wincker and Olivier Jaillon},
   doi = {10.1038/srep37900},
   journal = {Nature Publishing Group},
   title = {Survey of the green picoalga Bathycoccus genomes in the global ocean OPEN},
   url = {www.nature.com/scientificreports},
   year = {2016},
}
@article{Pitsch2019,
   abstract = {Ciliates represent central nodes in freshwater planktonic food webs, and many species show pronounced seasonality, with short-lived maxima of a few dominant taxa while many being rare or ephemeral. These observations are primarily based on morphospecies counting methods, which, however, have limitations concerning the amount and volume of samples that can be processed. For high sampling frequencies at large scales, high throughput sequencing (HTS) of freshwater ciliates seems to be a promising tool. However, several studies reported large discrepancy between species abundance determinations by molecular compared to morphological means. Therefore, we compared ciliate DNA metabarcodes (V9 regions of the 18S rRNA gene) with morphospecies counts for a 3-year study (Lake Zurich, Switzerland; biweekly sampling, n = 74). In addition, we isolated, cultivated and sequenced the 18S rRNA gene of twelve selected ciliate species that served as seeds for HTS analyses. This workflow allowed for a detailed comparison of V9 data with microscopic analyses by quantitative protargol staining (QPS). The dynamics of V9 read abundances over the seasonal cycle corroborated well with morphospecies population patterns. Annual successions of rare and ephemeral species were more adequately characterized by V9 reads than by QPS. However, numbers of species specific sequence reads only partly reflected rank orders seen by counts. In contrast, biomass-based assemblage compositions showed higher similarity to V9 read numbers, probably indicating a relation between cell sizes and numbers / sizes of macronuclei (or 18S rRNA operons). Full-length 18S rRNA sequences of ciliates assigned to certain morphospecies are urgently needed for barcoding approaches as planktonic taxa are still poorly represented in public databases and the interpretation of HTS data depends on profound reference sequences. Through linking operational taxonomic units (OTUs) with known morphospecies, we can use the deep knowledge about the autecology of these species.},
   author = {Gianna Pitsch and Estelle Patricia Bruni and Dominik Forster and Zhishuai Qu and Bettina Sonntag and Thorsten Stoeck and Thomas Posch},
   doi = {10.3389/FMICB.2019.00248/BIBTEX},
   issn = {1664302X},
   issue = {FEB},
   journal = {Frontiers in Microbiology},
   keywords = {Freshwater ciliates,Lake Zurich,Morphospecies counts,Planktonic ciliates,Seasonality,V9 region of the SSU rRNA gene},
   month = {2},
   pages = {248},
   publisher = {Frontiers Media S.A.},
   title = {Seasonality of planktonic freshwater ciliates: Are analyses based on V9 regions of the 18S rRNA gene correlated with morphospecies counts?},
   volume = {10},
   year = {2019},
}
@article{guo2016quantitative,
  title={Quantitative analysis of dinoflagellates and diatoms community via Miseq sequencing of actin gene and v9 region of 18S rDNA},
  author={Guo, Liliang and Sui, Zhenghong and Liu, Yuan},
  journal={Scientific reports},
  volume={6},
  number={1},
  pages={34709},
  year={2016},
  publisher={Nature Publishing Group UK London}
}
@article{Zhao2018,
   abstract = {Recent advances in high-throughput sequencing and metabarcoding technologies are revolutionizing our understanding of the diversity and ecology of microbial eukaryotes (protists). The interpretatio...},
   author = {Yan Zhao and Zhenzhen Yi and Alan Warren and Weibo B. Song},
   doi = {10.1098/RSPB.2017.2159},
   issn = {14712954},
   issue = {1871},
   journal = {Proceedings of the Royal Society B: Biological Sciences},
   keywords = {Euplotes,ecology,eukaryotic microbes,integrative taxonomy,species delimitation,taxonomy and systematics Keywords: ecology},
   month = {1},
   pmid = {29367393},
   publisher = {
The Royal Society
},
   title = {Species delimitation for the molecular taxonomy and ecology of the widely distributed microbial eukaryote genus Euplotes (Alveolata, Ciliophora)},
   volume = {285},
   url = {https://royalsocietypublishing.org/doi/10.1098/rspb.2017.2159},
   year = {2018},
}
@misc{karlusich2022biogeographical,
  title={Biogeographical patterns and genomes of aquatic photoautotrophs},
  author={Karlusich, Juan Jos{\'e} Pierella and Nef, Charlotte and Bowler, Chris and Dorrell, Richard G},
  year={2022},
  publisher={Wiley}
}
@article{pierella2023robust,
  title={A robust approach to estimate relative phytoplankton cell abundances from metagenomes},
  author={Pierella Karlusich, Juan Jos{\'e} and Pelletier, Eric and Zinger, Lucie and Lombard, Fabien and Zingone, Adriana and Colin, S{\'e}bastien and Gasol, Josep M and Dorrell, Richard G and Henry, Nicolas and Scalco, Eleonora and others},
  journal={Molecular Ecology Resources},
  volume={23},
  number={1},
  pages={16--40},
  year={2023},
  publisher={Wiley Online Library}
}
@article{edvardsen2016diversity,
  title={Diversity and distribution of haptophytes revealed by environmental sequencing and metabarcoding--a review},
  author={Edvardsen, Bente and Egge, Elianne Sirn{\ae}s and Vaulot, Daniel},
  journal={Perspect. Phycol},
  volume={3},
  pages={77--91},
  year={2016}
}
@article{chen2020accurate,
  title={Accurate and complete genomes from metagenomes},
  author={Chen, Lin-Xing and Anantharaman, Karthik and Shaiber, Alon and Eren, A Murat and Banfield, Jillian F},
  journal={Genome Research},
  volume={30},
  number={3},
  pages={315--333},
  year={2020},
  publisher={Cold Spring Harbor Lab}
}
@article{saary2020estimating,
  title={Estimating the quality of eukaryotic genomes recovered from metagenomic analysis with EukCC},
  author={Saary, Paul and Mitchell, Alex L and Finn, Robert D},
  journal={Genome biology},
  volume={21},
  number={1},
  pages={1--21},
  year={2020},
  publisher={BioMed Central}
}
@article{Frioux2020,
   abstract = {Metagenomic sequencing of complete microbial communities has greatly enhanced our understanding of the taxonomic composition of microbiotas. This has led to breakthrough developments in bioinformatic disciplines such as assembly, gene clustering, metagenomic binning of species genomes and the discovery of an incredible, so far undiscovered, taxonomic diversity. However, functional annotations and estimating metabolic processes from single species – or communities – is still challenging. Earlier approaches relied mostly on inferring the presence of key enzymes for metabolic pathways in the whole metagenome, ignoring the genomic context of such enzymes, resulting in the ‘bag-of-genes’ approach to estimate functional capacities of microbiotas. Here, we review recent developments in metagenomic bioinformatics, with a special focus on emerging technologies to simulate and estimate metabolic information, that can be derived from metagenomic assembled genomes. Genome-scale metabolic models can be used to model the emergent properties of microbial consortia and whole communities, and the progress in this area is reviewed. While this subfield of metagenomics is still in its infancy, it is becoming evident that there is a dire need for further bioinformatic tools to address the complex combinatorial problems in modelling the metabolism of large communities as a ‘bag-of-genomes’.},
   author = {Clémence Frioux and Dipali Singh and Tamas Korcsmaros and Falk Hildebrand},
   doi = {10.1016/J.CSBJ.2020.06.028},
   issn = {2001-0370},
   journal = {Computational and Structural Biotechnology Journal},
   keywords = {Bioinformatics,Gene Functions,Metabolic modelling,Metagenomic-assembled genomes,Metagenomics,Microbiota,Systems biology},
   month = {1},
   pages = {1722-1734},
   publisher = {Elsevier},
   title = {From bag-of-genes to bag-of-genomes: metabolic modelling of communities in the era of metagenome-assembled genomes},
   volume = {18},
   year = {2020},
}
@article{giner2016environmental,
  title={Environmental sequencing provides reasonable estimates of the relative abundance of specific picoeukaryotes},
  author={Giner, Caterina R and Forn, Irene and Romac, Sarah and Logares, Ramiro and De Vargas, Colomban and Massana, Ramon},
  journal={Applied and Environmental Microbiology},
  volume={82},
  number={15},
  pages={4757--4766},
  year={2016},
  publisher={Am Soc Microbiol}
}
@article{de2015eukaryotic,
  title={Eukaryotic plankton diversity in the sunlit ocean},
  author={De Vargas, Colomban and Audic, St{\'e}phane and Henry, Nicolas and Decelle, Johan and Mah{\'e}, Fr{\'e}d{\'e}ric and Logares, Ramiro and Lara, Enrique and Berney, C{\'e}dric and Le Bescot, Noan and Probert, Ian and others},
  journal={Science},
  volume={348},
  number={6237},
  pages={1261605},
  year={2015},
  publisher={American Association for the Advancement of Science}
}
@article{quince2017shotgun,
  title={Shotgun metagenomics, from sampling to analysis},
  author={Quince, Christopher and Walker, Alan W and Simpson, Jared T and Loman, Nicholas J and Segata, Nicola},
  journal={Nature biotechnology},
  volume={35},
  number={9},
  pages={833--844},
  year={2017},
  publisher={Nature Publishing Group US New York}
}
@article{eren2015anvi,
  title={Anvi’o: an advanced analysis and visualization platform for ‘omics data},
  author={Eren, A Murat and Esen, {\"O}zcan C and Quince, Christopher and Vineis, Joseph H and Morrison, Hilary G and Sogin, Mitchell L and Delmont, Tom O},
  journal={PeerJ},
  volume={3},
  pages={e1319},
  year={2015},
  publisher={PeerJ Inc.}
}
@article{sekse2017high,
  title={High throughput sequencing for detection of foodborne pathogens},
  author={Sekse, Camilla and Holst-Jensen, Arne and Dobrindt, Ulrich and Johannessen, Gro S and Li, Weihua and Spilsberg, Bj{\o}rn and Shi, Jianxin},
  journal={Frontiers in Microbiology},
  volume={8},
  pages={2029},
  year={2017},
  publisher={Frontiers Media SA}
}
@article{geisen2019user,
  title={A user guide to environmental protistology: primers, metabarcoding, sequencing, and analyses},
  author={Geisen, Stefan and Vaulot, Daniel and Mah{\'e}, Fr{\'e}d{\'e}ric and Lara, Enrique and de Vargas, Colomban and Bass, David},
  journal={BioRxiv},
  pages={850610},
  year={2019},
  publisher={Cold Spring Harbor Laboratory}
}
@article{gong2019estimation,
  title={Estimation of 18S gene copy number in marine eukaryotic plankton using a next-generation sequencing approach},
  author={Gong, Weida and Marchetti, Adrian},
  journal={Frontiers in marine Science},
  volume={6},
  pages={219},
  year={2019},
  publisher={Frontiers Media SA}
}
@article{latz2022short,
  title={Short-and long-read metabarcoding of the eukaryotic rRNA operon: evaluation of primers and comparison to shotgun metagenomics sequencing},
  author={Latz, Meike AC and Grujcic, Vesna and Brugel, Sonia and Lycken, Jenny and John, Uwe and Karlson, Bengt and Andersson, Agneta and Andersson, Anders F},
  journal={Molecular Ecology Resources},
  volume={22},
  number={6},
  pages={2304--2318},
  year={2022},
  publisher={Wiley Online Library}
}
@article{pawlowski2012cbol,
  title={CBOL protist working group: barcoding eukaryotic richness beyond the animal, plant, and fungal kingdoms},
  author={Pawlowski, Jan and Audic, St{\'e}phane and Adl, Sina and Bass, David and Belbahri, Lassa{\^a}d and Berney, C{\'e}dric and Bowser, Samuel S and Cepicka, Ivan and Decelle, Johan and Dunthorn, Micah and others},
  journal={PLoS biology},
  volume={10},
  number={11},
  pages={e1001419},
  year={2012},
  publisher={Public Library of Science San Francisco, USA}
}
@article{V9data,
   author = {Frédéric Mahé and Nicolas Henry and Colomban de Vargas and Coordinators Tara Oceans Consortium and Participants Tara Oceans Expedition},
   doi = {10.5281/ZENODO.7236051},
   keywords = {18S,Tara Oceans,eukaryotes,metabarcoding,plankton},
   month = {7},
   title = {rDNA 18S V9 metabarcoding tables (Swarm) for Tara Oceans Expedition (2009-2013), including Tara Polar Circle Expedition (2013)},
   url = {https://zenodo.org/record/7236051},
   year = {2022},
}
@article{quinn2017propr,
  title={propr: an R-package for identifying proportionally abundant features using compositional data analysis},
  author={Quinn, Thomas P and Richardson, Mark F and Lovell, David and Crowley, Tamsyn M},
  journal={Scientific reports},
  volume={7},
  number={1},
  pages={1--9},
  year={2017},
  publisher={Nature Publishing Group}
}
@article{erb2016should,
  title={How should we measure proportionality on relative gene expression data?},
  author={Erb, Ionas and Notredame, Cedric},
  journal={Theory in Biosciences},
  volume={135},
  pages={21--36},
  year={2016},
  publisher={Springer}
}
@article{quinn2019field,
  title={A field guide for the compositional analysis of any-omics data},
  author={Quinn, Thomas P and Erb, Ionas and Gloor, Greg and Notredame, Cedric and Richardson, Mark F and Crowley, Tamsyn M},
  journal={GigaScience},
  volume={8},
  number={9},
  pages={giz107},
  year={2019},
  publisher={Oxford University Press}
}
@article{lovell2015proportionality,
  title={Proportionality: a valid alternative to correlation for relative data},
  author={Lovell, David and Pawlowsky-Glahn, Vera and Egozcue, Juan Jos{\'e} and Marguerat, Samuel and B{\"a}hler, J{\"u}rg},
  journal={PLoS computational biology},
  volume={11},
  number={3},
  pages={e1004075},
  year={2015},
  publisher={Public Library of Science San Francisco, CA USA}
}
@article{delmont2022functional,
  title={Functional repertoire convergence of distantly related eukaryotic plankton lineages abundant in the sunlit ocean},
  author={Delmont, Tom O and Gaia, Morgan and Hinsinger, Damien D and Fr{\'e}mont, Paul and Vanni, Chiara and Fernandez-Guerra, Antonio and Eren, A Murat and Kourlaiev, Artem and d'Agata, Leo and Clayssen, Quentin and others},
  journal={Cell Genomics},
  volume={2},
  number={5},
  pages={100123},
  year={2022},
  publisher={Elsevier}
}
@article{V4data,
   author = {Frédéric Mahé and Nicolas Henry and Colomban de Vargas and Coordinators Tara Oceans Consortium and Participants Tara Oceans Expedition},
   doi = {10.5281/ZENODO.7235995},
   month = {10},
   title = {rDNA 18S V4 metabarcoding tables (Swarm) for Tara Oceans Expedition (2009-2013), including Tara Polar Circle Expedition (2013)},
   url = {https://zenodo.org/record/7235995},
   year = {2022},
}
@article{obiol2020metagenomic,
  title={A metagenomic assessment of microbial eukaryotic diversity in the global ocean},
  author={Obiol, Aleix and Giner, Caterina R and S{\'a}nchez, Pablo and Duarte, Carlos M and Acinas, Silvia G and Massana, Ramon},
  journal={Molecular Ecology Resources},
  volume={20},
  number={3},
  pages={718--731},
  year={2020},
  publisher={Wiley Online Library}
}
@article{grossart2020linking,
  title={Linking metagenomics to aquatic microbial ecology and biogeochemical cycles},
  author={Grossart, Hans-Peter and Massana, Ramon and McMahon, Katherine D and Walsh, David A},
  journal={Limnology and Oceanography},
  volume={65},
  pages={S2--S20},
  year={2020},
  publisher={Wiley Online Library}
}
@article{catlett2020evaluation,
  title={Evaluation of accuracy and precision in an amplicon sequencing workflow for marine protist communities},
  author={Catlett, Dylan and Matson, Paul G and Carlson, Craig A and Wilbanks, Elizabeth G and Siegel, David A and Iglesias-Rodriguez, M Debora},
  journal={Limnology and Oceanography: Methods},
  volume={18},
  number={1},
  pages={20--40},
  year={2020},
  publisher={Wiley Online Library}
}
@article{lavrinienko2021does,
  title={Does intraspecific variation in rDNA copy number affect analysis of microbial communities?},
  author={Lavrinienko, Anton and Jernfors, Toni and Koskim{\"a}ki, Janne J and Pirttil{\"a}, Anna Maria and Watts, Phillip C},
  journal={Trends in microbiology},
  volume={29},
  number={1},
  pages={19--27},
  year={2021},
  publisher={Elsevier}
}
@article{bell2021comparing,
  title={Comparing whole-genome shotgun sequencing and DNA metabarcoding approaches for species identification and quantification of pollen species mixtures},
  author={Bell, Karen L and Petit III, Robert A and Cutler, Anya and Dobbs, Emily K and Macpherson, J Michael and Read, Timothy D and Burgess, Kevin S and Brosi, Berry J},
  journal={Ecology and Evolution},
  volume={11},
  number={22},
  pages={16082--16098},
  year={2021},
  publisher={Wiley Online Library}
}
@article{paula2022metabarcoding,
  title={Metabarcoding versus mapping unassembled shotgun reads for identification of prey consumed by arthropod epigeal predators},
  author={Paula, D{\'e}bora Pires and Barros, Suellen Karina Albertoni and Pitta, Rafael Major and Barreto, Marliton Rocha and Togawa, Roberto Coiti and Andow, David A},
  journal={GigaScience},
  volume={11},
  year={2022},
  publisher={Oxford Academic}
}
@article{creer2016ecologist,
  title={The ecologist's field guide to sequence-based identification of biodiversity},
  author={Creer, Simon and Deiner, Kristy and Frey, Serita and Porazinska, Dorota and Taberlet, Pierre and Thomas, W Kelley and Potter, Caitlin and Bik, Holly M},
  journal={Methods in Ecology and Evolution},
  volume={7},
  number={9},
  pages={1008--1018},
  year={2016},
  publisher={Wiley Online Library}
}
@article{zepeda2015environmental,
  title={Environmental genes and genomes: understanding the differences and challenges in the approaches and software for their analyses},
  author={Zepeda Mendoza, Marie Lisandra and Sicheritz-Ponten, Thomas and Gilbert, M Thomas P},
  journal={Briefings in Bioinformatics},
  volume={16},
  number={5},
  pages={745--758},
  year={2015},
  publisher={Oxford University Press}
}
@article{porter2018scaling,
  title={Scaling up: A guide to high-throughput genomic approaches for biodiversity analysis},
  author={Porter, Teresita M and Hajibabaei, Mehrdad},
  journal={Molecular ecology},
  volume={27},
  number={2},
  pages={313--338},
  year={2018},
  publisher={Wiley Online Library}
}
@article{erb2016should,
  title={How should we measure proportionality on relative gene expression data?},
  author={Erb, Ionas and Notredame, Cedric},
  journal={Theory in Biosciences},
  volume={135},
  pages={21--36},
  year={2016},
  publisher={Springer}
}
@article{patuzzi2019metasparsim,
  title={metaSPARSim: a 16S rRNA gene sequencing count data simulator},
  author={Patuzzi, Ilaria and Baruzzo, Giacomo and Losasso, Carmen and Ricci, Antonia and Di Camillo, Barbara},
  journal={BMC bioinformatics},
  volume={20},
  number={9},
  pages={1--13},
  year={2019},
  publisher={BioMed Central}
}
  @Manual{Rbase,
    title = {R: A Language and Environment for Statistical Computing},
    author = {{R Core Team}},
    organization = {R Foundation for Statistical Computing},
    address = {Vienna, Austria},
    year = {2020},
    url = {https://www.R-project.org/},
  }
@Article{randomizematrix,
    title = {Picante: {R} tools for integrating phylogenies and ecology},
    author = {S.W. Kembel and P.D. Cowan and M.R. Helmus and W.K. Cornwell and H. Morlon and D.D. Ackerly and S.P. Blomberg and C.O. Webb},
    journal = {Bioinformatics},
    volume = {26},
    pages = {1463--1464},
    year = {2010},
  }


@article{perezcobas2020,
  title={ Metagenomic approaches in microbial ecology: an update on whole-genome and marker gene sequencing analyses},
  author={Pérez-Cobas A.E., Gomez-Valero L., Buchrieser C.},
  journal={Microb Genom},
  volume={6},
  number={8},
  pages={},
  year={2020},
  publisher={BioMed Central}
}

@article{rondon2000,
author = {Michelle R. Rondon and Paul R. August and Alan D. Bettermann and Sean F. Brady and Trudy H. Grossman and Mark R. Liles and Kara A. Loiacono and Berkley A. Lynch and Ian A. MacNeil and Charles Minor and Choi Lai Tiong and Michael Gilman and Marcia S. Osburne and Jon Clardy and Jo Handelsman and Robert M. Goodman},
title = {Cloning the Soil Metagenome: a Strategy for Accessing the Genetic and Functional Diversity of Uncultured Microorganisms},
journal = {Applied and Environmental Microbiology},
volume = {66},
number = {6},
pages = {2541-2547},
year = {2000},
doi = {10.1128/AEM.66.6.2541-2547.2000},

URL = {https://journals.asm.org/doi/abs/10.1128/AEM.66.6.2541-2547.2000},
eprint = {https://journals.asm.org/doi/pdf/10.1128/AEM.66.6.2541-2547.2000}
,
    abstract = { Recent progress in molecular microbial ecology has revealed that traditional culturing methods fail to represent the scope of microbial diversity in nature, since only a small proportion of viable microorganisms in a sample are recovered by culturing techniques. To develop methods to investigate the full extent of microbial diversity, we used a bacterial artificial chromosome (BAC) vector to construct libraries of genomic DNA isolated directly from soil (termed metagenomic libraries). To date, we have constructed two such libraries, which contain more than 1 Gbp of DNA. Phylogenetic analysis of 16S rRNA gene sequences recovered from one of the libraries indicates that the BAC libraries contain DNA from a wide diversity of microbial phyla, including sequences from diverse taxa such as the low-G+C, gram-positive Acidobacterium,Cytophagales, and Proteobacteria. Initial screening of the libraries in Escherichia coli identified several clones that express heterologous genes from the inserts, confirming that the BAC vector can be used to maintain, express, and analyze environmental DNA. The phenotypes expressed by these clones include antibacterial, lipase, amylase, nuclease, and hemolytic activities. Metagenomic libraries are a powerful tool for exploring soil microbial diversity, providing access to the genetic information of uncultured soil microorganisms. Such libraries will be the basis of new initiatives to conduct genomic studies that link phylogenetic and functional information about the microbiota of environments dominated by microorganisms that are refractory to cultivation. }
}

@article{swarm,
    author = {Mahé, Frédéric and Czech, Lucas and Stamatakis, Alexandros and Quince, Christopher and de Vargas, Colomban and Dunthorn, Micah and Rognes, Torbjørn},
    title = "{Swarm v3: towards tera-scale amplicon clustering}",
    journal = {Bioinformatics},
    volume = {38},
    number = {1},
    pages = {267-269},
    year = {2021},
    month = {07},
    abstract = "{Previously we presented swarm, an open-source amplicon clustering programme that produces fine-scale molecular operational taxonomic units (OTUs) that are free of arbitrary global clustering thresholds. Here, we present swarm v3 to address issues of contemporary datasets that are growing towards tera-byte sizes.When compared with previous swarm versions, swarm v3 has modernized C++ source code, reduced memory footprint by up to 50\\%, optimized CPU-usage and multithreading (more than 7 times faster with default parameters), and it has been extensively tested for its robustness and logic.Source code and binaries are available at https://github.com/torognes/swarm.Supplementary data are available at Bioinformatics online.}",
    issn = {1367-4803},
    doi = {10.1093/bioinformatics/btab493},
    url = {https://doi.org/10.1093/bioinformatics/btab493},
    eprint = {https://academic.oup.com/bioinformatics/article-pdf/38/1/267/49006260/btab493.pdf},
}
@article{Ayling2019,
    author = {Ayling, Martin and Clark, Matthew D and Leggett, Richard M},
    title = "{New approaches for metagenome assembly with short reads}",
    journal = {Briefings in Bioinformatics},
    volume = {21},
    number = {2},
    pages = {584-594},
    year = {2019},
    month = {02},
    abstract = "{In recent years, the use of longer range read data combined with advances in assembly algorithms has stimulated big improvements in the contiguity and quality of genome assemblies. However, these advances have not directly transferred to metagenomic data sets, as assumptions made by the single genome assembly algorithms do not apply when assembling multiple genomes at varying levels of abundance. The development of dedicated assemblers for metagenomic data was a relatively late innovation and for many years, researchers had to make do using tools designed for single genomes. This has changed in the last few years and we have seen the emergence of a new type of tool built using different principles. In this review, we describe the challenges inherent in metagenomic assemblies and compare the different approaches taken by these novel assembly tools.}",
    issn = {1477-4054},
    doi = {10.1093/bib/bbz020},
    url = {https://doi.org/10.1093/bib/bbz020},
    eprint = {https://academic.oup.com/bib/article-pdf/21/2/584/33583908/bbz020.pdf},
}


@article {richterselfpromortion,
article_type = {journal},
title = {Genomic evidence for global ocean plankton biogeography shaped by large-scale current systems},
author = {Richter, Daniel J and Watteaux, Romain and Vannier, Thomas and Leconte, Jade and Frémont, Paul and Reygondeau, Gabriel and Maillet, Nicolas and Henry, Nicolas and Benoit, Gaëtan and Da Silva, Ophélie and Delmont, Tom O and Fernàndez-Guerra, Antonio and Suweis, Samir and Narci, Romain and Berney, Cédric and Eveillard, Damien and Gavory, Frederick and Guidi, Lionel and Labadie, Karine and Mahieu, Eric and Poulain, Julie and Romac, Sarah and Roux, Simon and Dimier, Céline and Kandels, Stefanie and Picheral, Marc and Searson, Sarah and Tara Oceans Coordinators and Pesant, Stéphane and Aury, Jean-Marc and Brum, Jennifer R and Lemaitre, Claire and Pelletier, Eric and Bork, Peer and Sunagawa, Shinichi and Lombard, Fabien and Karp-Boss, Lee and Bowler, Chris and Sullivan, Matthew B and Karsenti, Eric and Mariadassou, Mahendra and Probert, Ian and Peterlongo, Pierre and Wincker, Patrick and de Vargas, Colomban and Ribera d'Alcalà, Maurizio and Iudicone, Daniele and Jaillon, Olivier},
editor = {Louca, Stilianos and Schuman, Meredith C},
volume = 11,
year = 2022,
month = {aug},
pub_date = {2022-08-03},
pages = {e78129},
citation = {eLife 2022;11:e78129},
doi = {10.7554/eLife.78129},
url = {https://doi.org/10.7554/eLife.78129},
abstract = {Biogeographical studies have traditionally focused on readily visible organisms, but recent technological advances are enabling analyses of the large-scale distribution of microscopic organisms, whose biogeographical patterns have long been debated. Here we assessed the global structure of plankton geography and its relation to the biological, chemical, and physical context of the ocean (the ‘seascape’) by analyzing metagenomes of plankton communities sampled across oceans during the \textit{Tara} Oceans expedition, in light of environmental data and ocean current transport. Using a consistent approach across organismal sizes that provides unprecedented resolution to measure changes in genomic composition between communities, we report a pan-ocean, size-dependent plankton biogeography overlying regional heterogeneity. We found robust evidence for a basin-scale impact of transport by ocean currents on plankton biogeography, and on a characteristic timescale of community dynamics going beyond simple seasonality or life history transitions of plankton.},
keywords = {plankton biogeography, metagenomics, metabarcoding, microbial oceanography},
journal = {eLife},
issn = {2050-084X},
publisher = {eLife Sciences Publications, Ltd},
}

@article{del2014others,
  title={The others: our biased perspective of eukaryotic genomes},
  author={Del Campo, Javier and Sieracki, Michael E and Molestina, Robert and Keeling, Patrick and Massana, Ramon and Ruiz-Trillo, I{\~n}aki},
  journal={Trends in ecology \& evolution},
  volume={29},
  number={5},
  pages={252--259},
  year={2014},
  publisher={Elsevier}
}
@article{ibarbalz2019global,
  title={Global trends in marine plankton diversity across kingdoms of life},
  author={Ibarbalz, Federico M and Henry, Nicolas and Brand{\~a}o, Manoela C and Martini, S{\'e}verine and Busseni, Greta and Byrne, Hannah and Coelho, Luis Pedro and Endo, Hisashi and Gasol, Josep M and Gregory, Ann C and others},
  journal={Cell},
  volume={179},
  number={5},
  pages={1084--1097},
  year={2019},
  publisher={Elsevier}
}
@article{boenigk2012concepts,
  title={Concepts in protistology: species definitions and boundaries},
  author={Boenigk, Jens and Ereshefsky, Marc and Hoef-Emden, Kerstin and Mallet, James and Bass, David},
  journal={European Journal of Protistology},
  volume={48},
  number={2},
  pages={96--102},
  year={2012},
  publisher={Elsevier}
}
@article{bai2022vertical,
  title={Vertical community patterns of Labyrinthulomycetes protists reveal their potential importance in the oceanic biological pump},
  author={Bai, Mohan and Xie, Ningdong and He, Yaodong and Li, Jiaqian and Collier, Jackie L and Hunt, Dana E and Johnson, Zackary I and Jiao, Nianzhi and Wang, Guangyi},
  journal={Environmental Microbiology},
  volume={24},
  number={4},
  pages={1703--1713},
  year={2022},
  publisher={Wiley Online Library}
}
@article{berney2017unieuk,
  title={UniEuk: time to speak a common language in protistology!},
  author={Berney, C{\'e}dric and Ciuprina, Andreea and Bender, Sara and Brodie, Juliet and Edgcomb, Virginia and Kim, Eunsoo and Rajan, Jeena and Parfrey, Laura Wegener and Adl, Sina and Audic, St{\'e}phane and others},
  journal={Journal of Eukaryotic Microbiology},
  volume={64},
  number={3},
  pages={407--411},
  year={2017},
  publisher={Wiley Online Library}
}
