## Supplementary material for "On the relationship between protist metabarcoding and protist metagenome-assembled genomes": LaTeX source for the pdf: main.tex

\documentclass{article}

\usepackage{biorxiv} % github.com/alexsbaldwin/biorxiv-inspired-latex-style


\usepackage[utf8]{inputenc} % allow utf-8 input
\usepackage[T1]{fontenc}    % use 8-bit T1 fonts
\usepackage{hyperref}       % hyperlinks
\usepackage{url}            % simple URL typesetting
\usepackage{booktabs}       % professional-quality tables
\usepackage{amsfonts}       % blackboard math symbols
\usepackage{nicefrac}       % compact symbols for 1/2, etc.
\usepackage{microtype}      % microtypography
\usepackage{lipsum}		% Can be removed after putting your text content
\usepackage{lineno}
\usepackage{graphicx}
\usepackage{nameref,hyperref}
\usepackage{geometry}
\usepackage{rotating}
\usepackage{svg}
%%%%%%%%
\usepackage{pdflscape}
\usepackage{afterpage}
\usepackage{capt-of}% or use the larger `caption` package
\usepackage{hyperref}
\usepackage{float}
\usepackage{cleveref}
\usepackage{import}


\linenumbers


\title{On the relationship between protist metabarcoding and protist metagenome-assembled genomes}

%\date{September 9, 1985}	% Here you can change the date presented in the paper title
%\date{} 					% Or removing it
\author{
  Daryna Zavadska\\
  https://orcid.org/0000-0002-8742-7044\\
  \texttt{} \\
  \scriptsize{Institut de Biologia Evolutiva (CSIC-Universitat Pompeu Fabra), } \\
  \scriptsize{Passeig Marítim de la Barceloneta 37-49, 08003 Barcelona, Spain} \\ 
  %% examples of more authors
   \And
  Nicolas Henry\\
  https://orcid.org/0000-0002-7702-1382\\
  \texttt{} \\
  \scriptsize{CNRS, FR2424, ABiMS, Station Biologique de Roscoff,}\\
  \scriptsize{Sorbonne Université, Roscoff, France}\\
  \AND
  Adrià Auladell \\
  https://orcid.org/0000-0003-2260-6946\\
  \texttt{} \\
  \scriptsize{Institut de Biologia Evolutiva (CSIC-Universitat Pompeu Fabra),} \\
  \scriptsize{Passeig Marítim de la Barceloneta 37-49, 08003 Barcelona, Spain} \\ 
  \And
  Cédric Berney\\
  https://orcid.org/0000-0001-8689-9907 \\
  \texttt{} \\
  \scriptsize{CNRS, UMR7144, AD2M, Station Biologique de Roscoff,}\\
  \scriptsize{Sorbonne Université, Roscoff, France}  \\
  \And
  Daniel J. Richter\\
  https://orcid.org/0000-0002-9238-5571 \\
  \texttt{} \\
  \scriptsize{Institut de Biologia Evolutiva (CSIC-Universitat Pompeu Fabra),} \\
  \scriptsize{Passeig Marítim de la Barceloneta 37-49, 08003 Barcelona, Spain} \\ 
}

% Uncomment to override  the `A preprint' in the header
%\renewcommand{\headeright}{}


\begin{document}
\maketitle


\begin{abstract}
%the nature of the problem 
The two most commonly used approaches to study the composition of environmental protist communities are marker gene metabarcoding and whole genome analysis through metagenomics. Raw metabarcoding data are usually processed into Operational Taxonomic Units (OTUs) or amplicon sequence variants (ASVs) through clustering or denoising approaches, respectively. Analogous approaches have been developed to assemble metagenomic sequence reads into metagenome-assembled genomes (MAGs). Understanding the correspondence between the data produced by these two approaches can help to integrate information between the datasets and to explain how metabarcoding OTUs and MAGs are related with the underlying biological entities they are hypothesised to represent. Due to the nature of their construction, MAGs do not contain the most commonly used barcoding loci, meaning that sequence homology approaches cannot be used to match OTUs and MAGs.

%what we did to try to solve it 
We made an attempt to match V9 metabarcoding OTUs from the 18S rRNA gene (V9 OTUs) and MAGs from the \textit{Tara} Oceans expedition (2009-2013) based on the correspondence of their relative abundances across the same set of samples.
We evaluated the performance of several methods for detecting correspondence between features in these two compositional datasets and developed a series of controls to filter artefacts of data structure and processing. After selecting the best-performing correspondence metrics, ranking the V9 OTU/MAG matches by their proportionality/correlation coefficients and applying a set of selection criteria, we identified candidate matches between V9 OTUs and MAGs. 

%our results, 
In a subset of cases, V9 OTUs and MAGs could be successfully matched with one another with a one-to-one correspondence, implying that they likely represent the same underlying biological entity. 
More generally, matches we observed could be classified into 4 scenarios: 
Scenario I - one V9 OTU matches more than one MAG;
Scenario II - more than one V9 OTU matches more than one MAG;
Scenario III - more than one V9 OTU matches one MAG;
Scenario IV - one V9 OTU matches one MAG.

%implications. 
These diverse scenarios for V9 OTU-MAG matches illustrate the complex nature of the OTU/MAG relationship. Notably, we found some instances in which different OTU-MAG matches from the same taxonomic group were not classified in the same scenario, with all four scenarios possible even within the same taxonomic group, illustrating that factors beyond taxonomic lineage influence the relationship between OTUs and MAGs. Overall, each scenario produces a different interpretation of V9 OTUs, MAGs and how they compare in terms of the genomic and ecological diversity that they represent.

\end{abstract}


% keywords can be removed
\keywords{Protist \and 18S V9 \and Metagenome-assembled genome \and Tara Oceans}


%\input{intro.tex}
%\input{methods.tex}
\input{results.tex}
%\input{discussion}
\input{suppl.tex}

%\bibliographystyle{apalike}  
%\bibliography{references}  % uses external references.bib file

\end{document}
