## Supplementary material for "On the relationship between protist metabarcoding and protist metagenome-assembled genomes": LaTeX source for the pdf: Figure_3.pdf

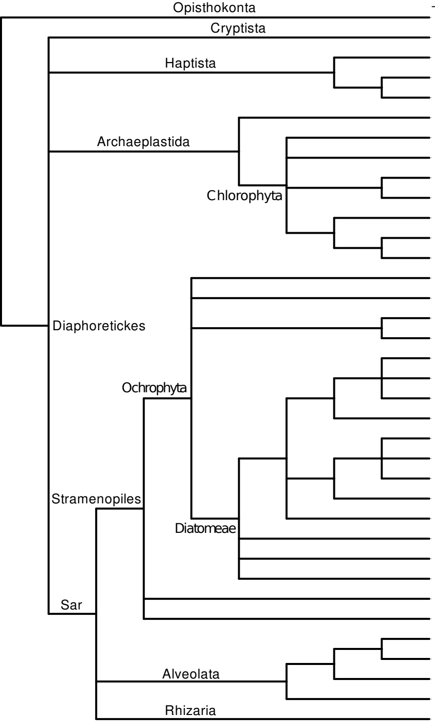


| Group | No. MAGs | No. OTUs in Taxonomic Subset | | Scenario I - Many MAGs, one V9 OTU | | | | Scenario II - Many MAGs, many V9 OTU | | | | Scenario III - One MAG, many V9 OTUs | | | | Scenario IV - One MAG, one V9 OTU | | | |
|---|---|---|---|---|---|---|---|---|---|---|---|---|---|---|---|---|---|---|---|
| | | Lower Rank | Higher Rank | High | Med. | Low | V.Low | High | Med. | Low | V.Low | High | Med. | Low | V.Low | High | Med. | Low | V.Low |
| unidentified Choanoflagellatea | 11 | 298 | 33,545 | | | | | | | | | | | | | 1 | 3 | | 1 |
| Geminigera | 7 | 5 | 239 | | | | | | | | | | 2 | | | | | | |
| Phaeocystis | 27 | 80 | 104 | | | 3 | 8 | | | 3 | 7 | | | | | | | | |
| Haptolina | 2 | 7 | 59 | | 2 | | | | | | | | | | | | | | |
| unidentified Chrysochromulinaceae | 39 | 333 | 746 | | 2 | 2 | 5 | | | 10 | 2 | | | 1 | 3 | 1 | | | 3 |
| Prasinoderma | 2 | 21 | 31 | | | | | | | | | | | | | 2 | | | |
| Picochlorum | 1 | 4 | 524 | | | | | | | | | | | | | 1 | | | |
| unidentified Chlorophyta | 6 | 7,089 | 7,436 | | | | | | | | | | | 1 | | 3 | | | 2 |
| Chloroparvula | 7 | 158 | 138 | | 2 | | | | 2 | | | | 1 | | | 1 | 1 | | |
| Chloropicon | 11 | 80 | 138 | | | | | | 1 | | 1 | | | | | 1 | | 1 | |
| Ostreococcus | 4 | 14 | 19 | | 1 | 2 | 1 | | | | | | | | | | | | |
| Micromonas | 20 | 44 | 68 | | 10 | | 3 | | | | | | | | | | 2 | 1 | |
| Mantoniella | 1 | 3 | 68 | | | | | | | | | | | | | 1 | | | |
| Florenciella | 3 | 0 | 57 | | 2 | | | | | | | | | | | | 1 | | |
| unidentified Chrysophyceae | 23 | 397 | 6,926 | | | | | | | 2 | 1 | | | 1 | 1 | | | 4 | |
| Aureococcus | 6 | 11 | 84 | | | | | | | 1 | 5 | | | | | | | | |
| Pelagococcus | 2 | 14 | 84 | | | | | | | 1 | 1 | | | | | | | | |
| Pseudo-nitzschia | 7 | 275 | 1,929 | | | | | | | | 2 | | | | | | | | 1 |
| Cylindrotheca | 2 | 11 | 1,929 | | | | | | | | | | | | | | 1 | | |
| Fragilariopsis | 5 | 135 | 1,929 | | | | 2 | | | | 3 | | | | | | | | |
| Synedropsis | 1 | 60 | 1,929 | | | | | | | | | | 1 | | | | | | |
| Thalassiosira | 5 | 593 | 2,325 | | 1 | | 2 | | | | | | | | | | | | 2 |
| Minidiscus | 2 | 5 | 2,325 | | | | 2 | | | | | | | | | | | | |
| Skeletonema | 3 | 45 | 2,325 | | | | | | | | | | | | | 1 | 2 | | |
| Chaetoceros | 11 | 632 | 2,325 | | | | | | | | | | | | 1 | 1 | 1 | | 1 |
| Attheya | 1 | 60 | 128 | | | | | | | | | 1 | | | | | | | |
| unidentified Bacillariaceae 3 | 2 | 5,430 | 6,926 | | | | | | | | | | | 1 | | | | | |
| unidentified Bacillariaceae 2 | 2 | 5,430 | 6,926 | | | | | | | | | | | | | | 1 | | 1 |
| Leptocylindrus | 1 | 35 | 53 | | | | | | | | | | | | | 1 | | | |
| unidentified Bicosoecida | 8 | 230 | 1,228 | | | | | | | | | | | | | | | | 1 |
| unidentified MAST-04 | 25 | 68 | 1,655 | | 1 | 5 | 2 | | | | | | | | | | | 1 | 1 |
| unidentified Choreotrichida | 15 | 1,361 | 2,121 | | | | 7 | | | 1 | 2 | | | | 1 | | | | 4 |
| unidentified Cyrtolophosidida | 4 | 69 | 182 | | | | | | | | | | | | | | 1 | | |
| unidentified Ciliophora | 2 | 18,850 | 78,782 | | | | | | | | | | | | 1 | | | | 1 |
| unidentified MALV-I 1 | 2 | 11,911 | 55,005 | | | | | | | | | | | | | | | | 1 |
| unidentified Cercozoa | 2 | 2,625 | 19,891 | | | | | | | | | | | | | | | | 1 |
