## Supplementary figures and images for "On the relationship between protist metabarcoding and protist metagenome-assembled genomes"

### alg_propr_calc.pdf

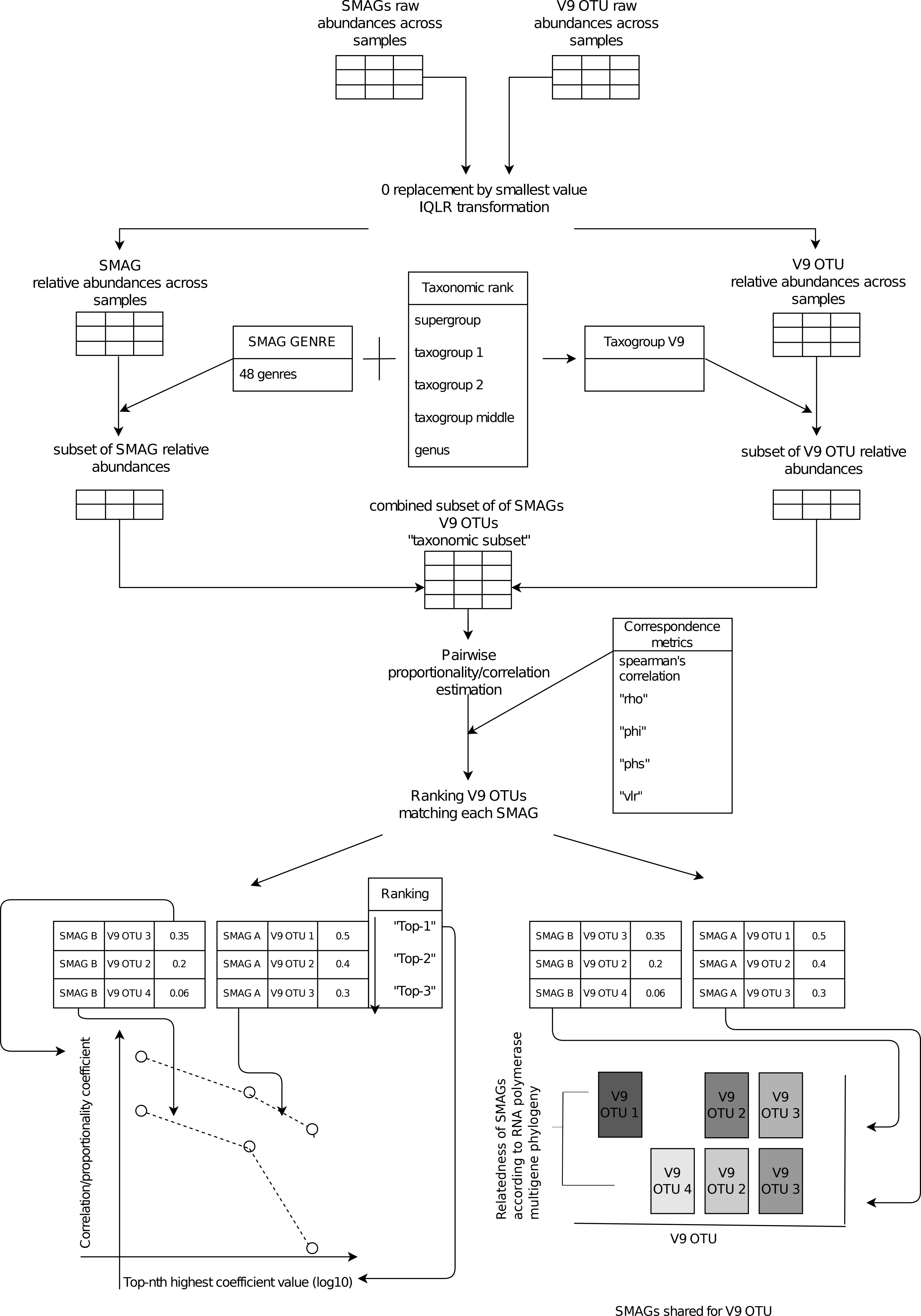

### concept.pdf

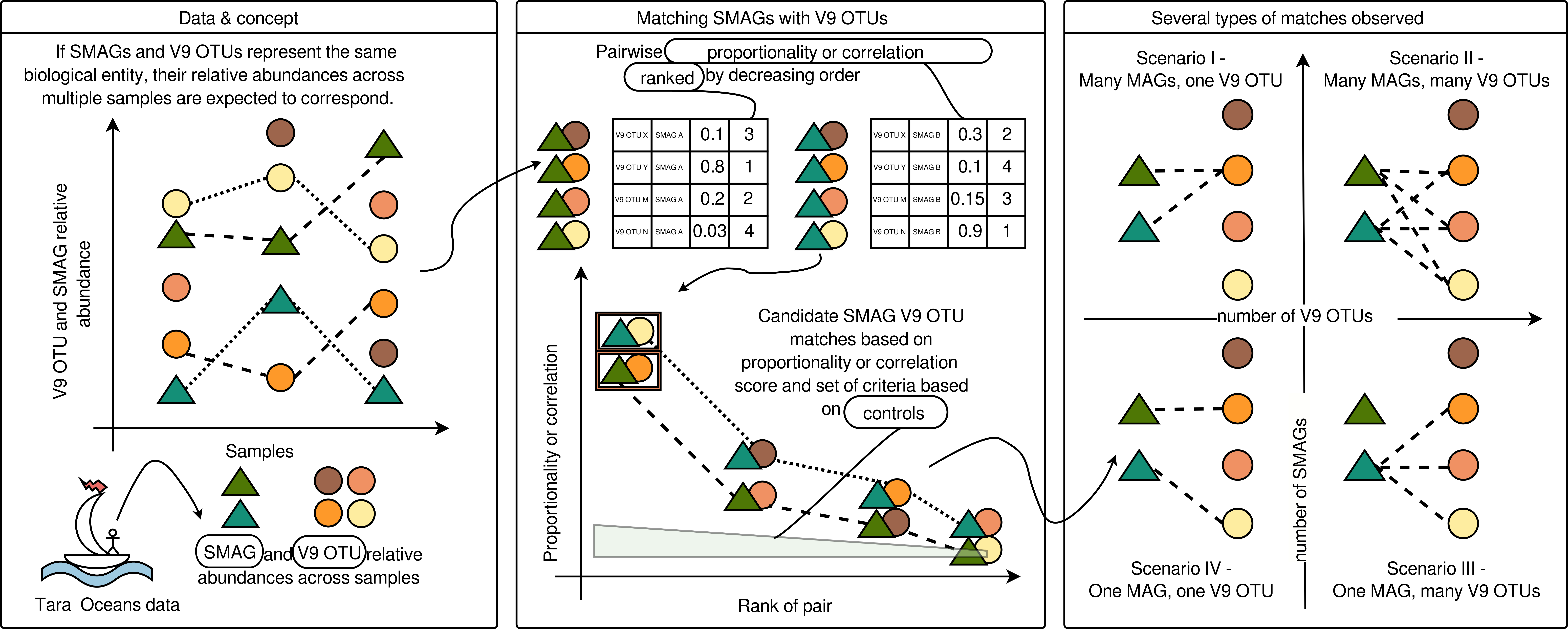

### concept_plot1.pdf

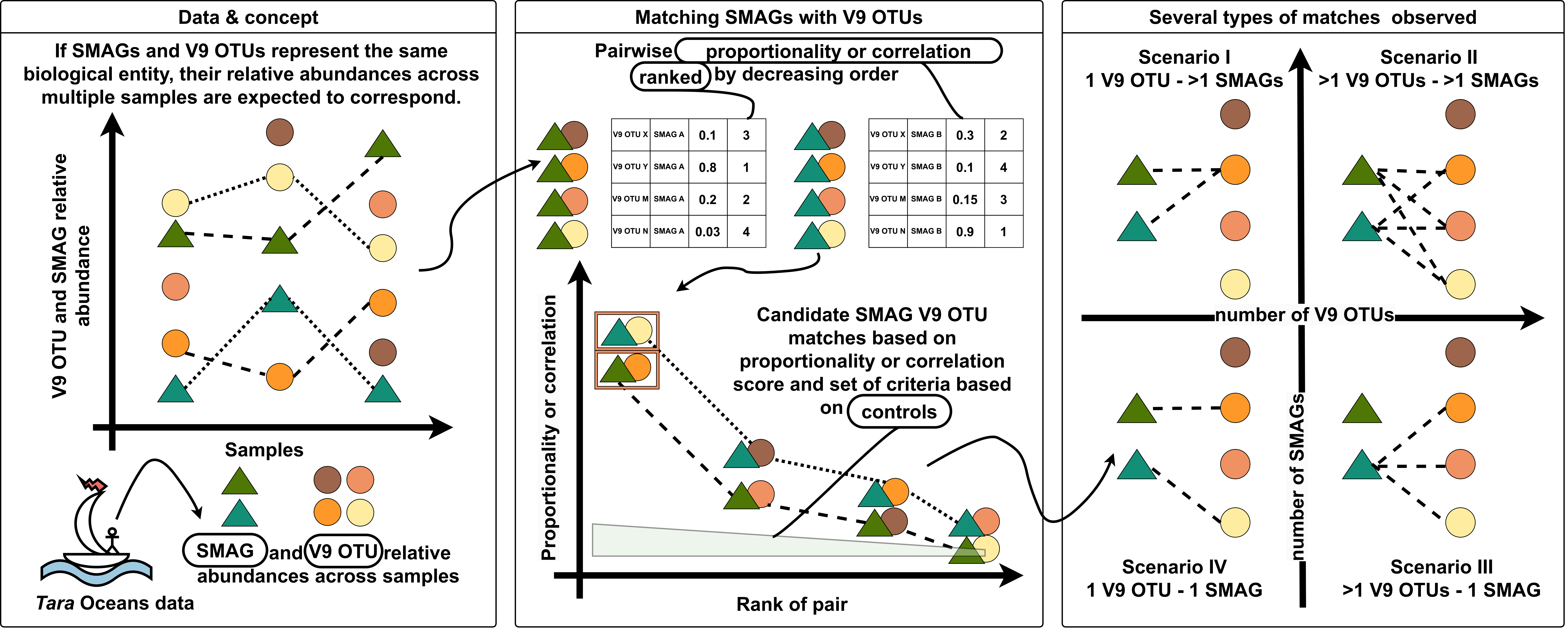

### cor_calc_plots.pdf

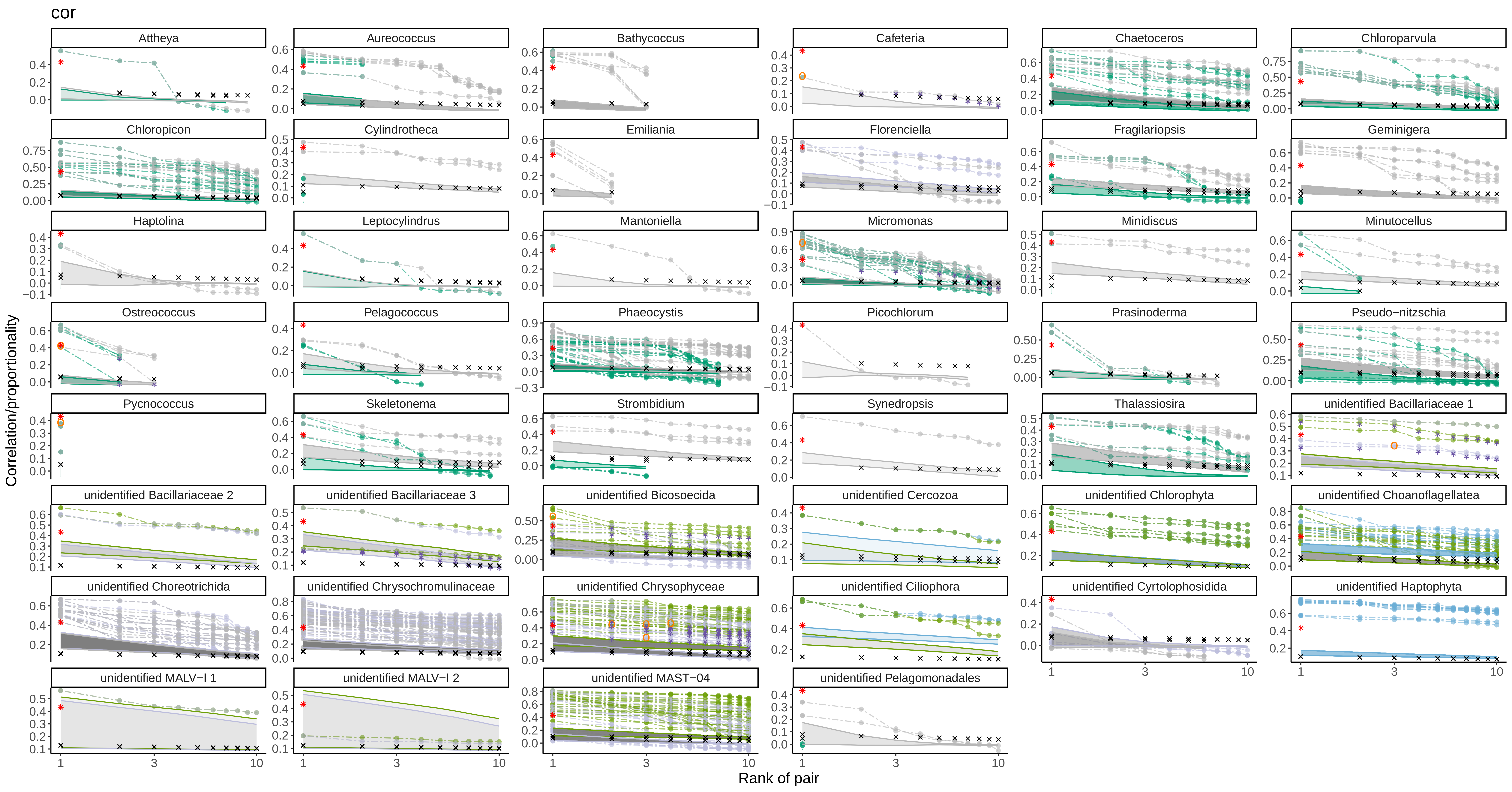

### cor_uniq_facet_plots2.pdf

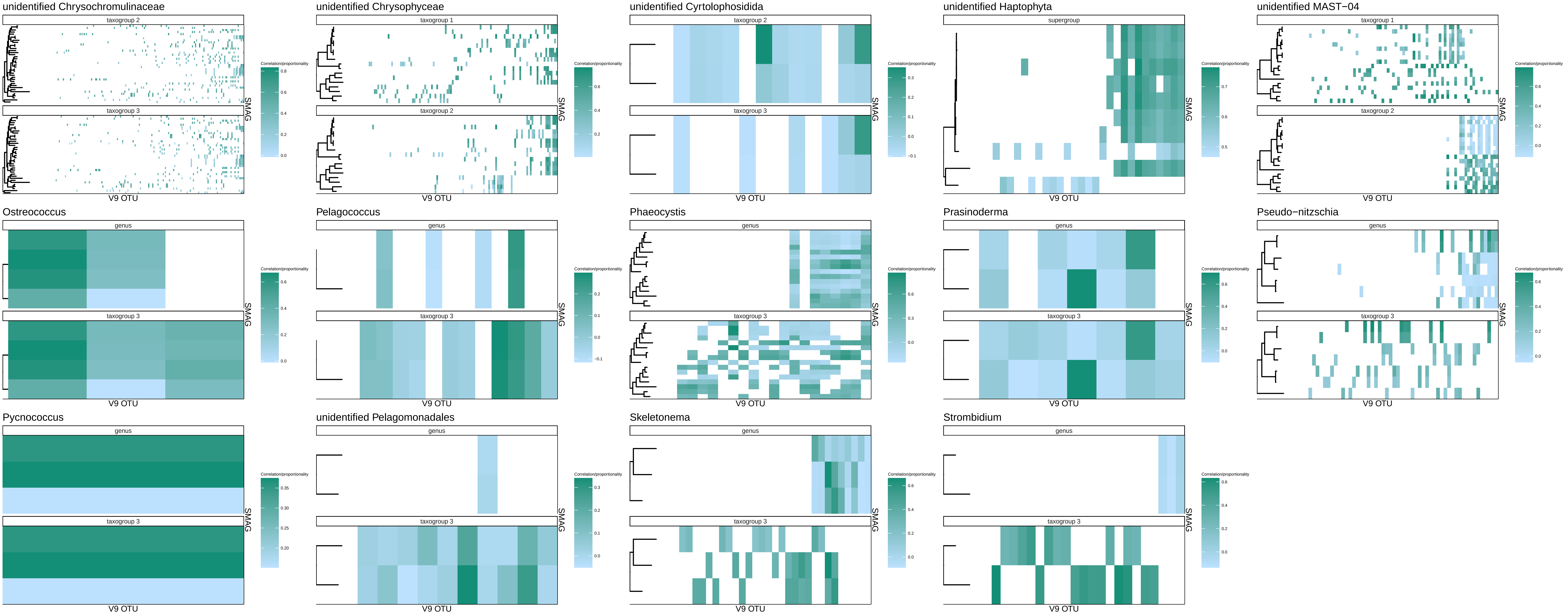

### cor_uniq_facet_plots.pdf

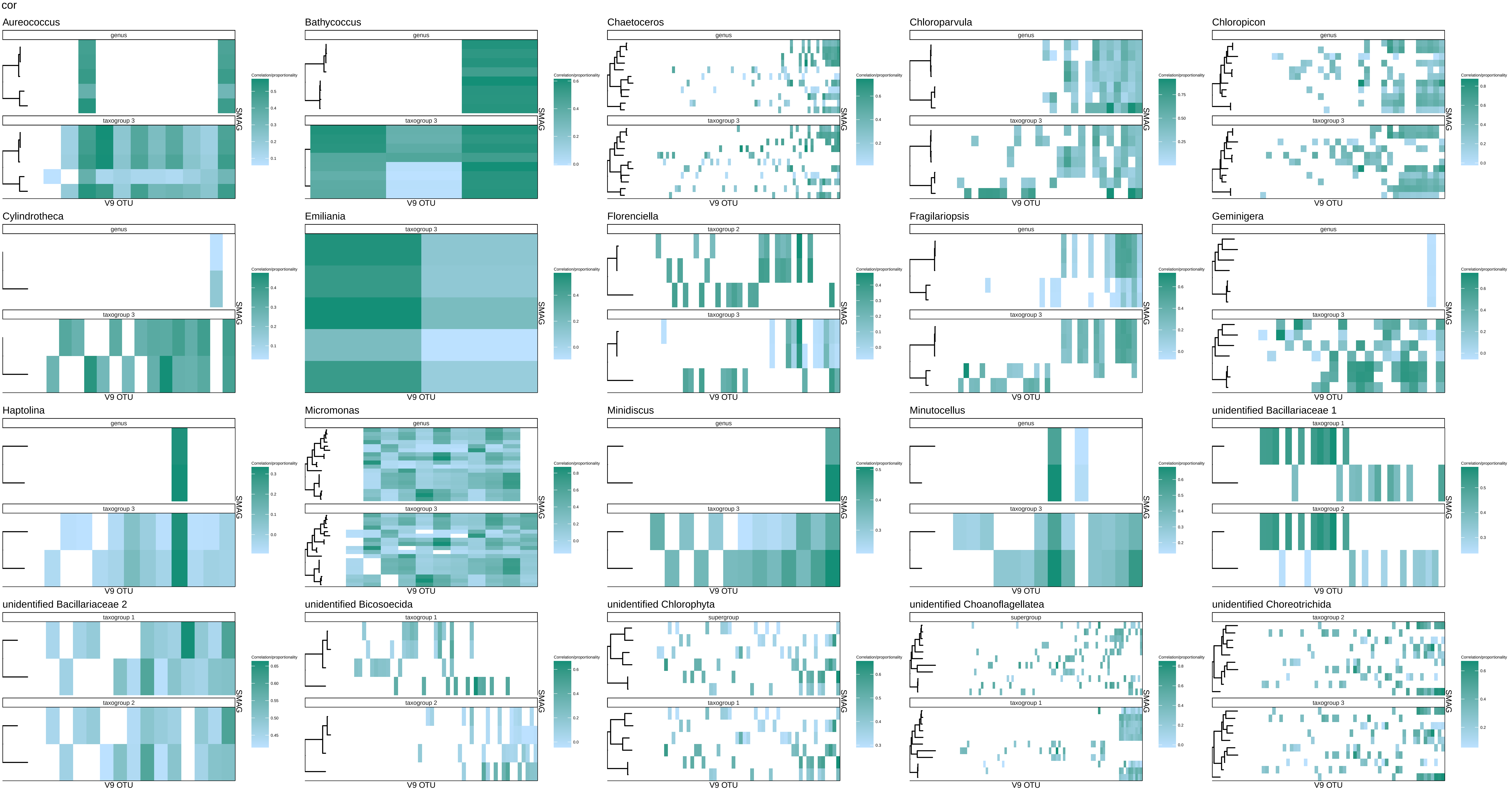

### draft_fig_1.pdf

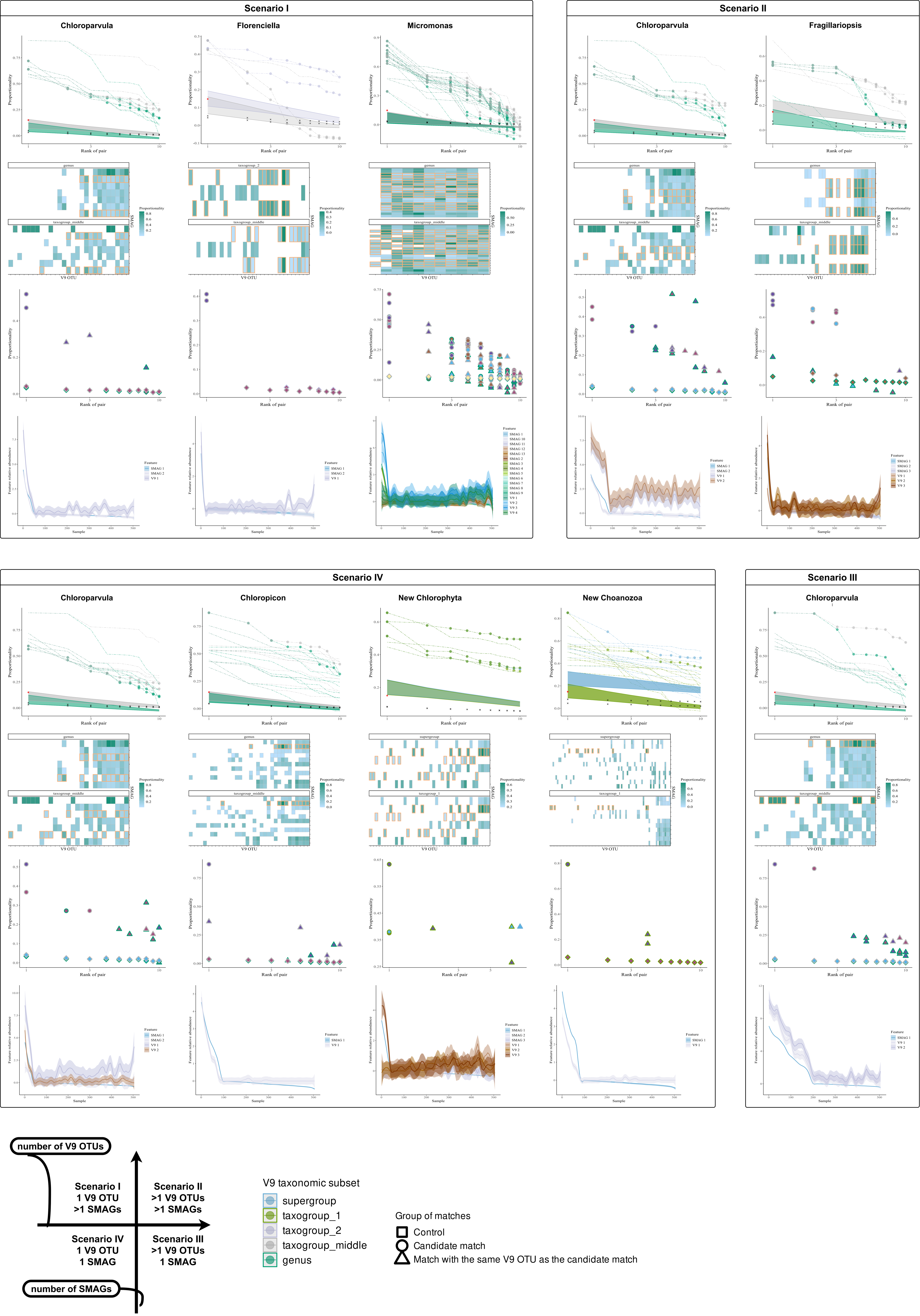

### draft_fig_1_edit1.pdf

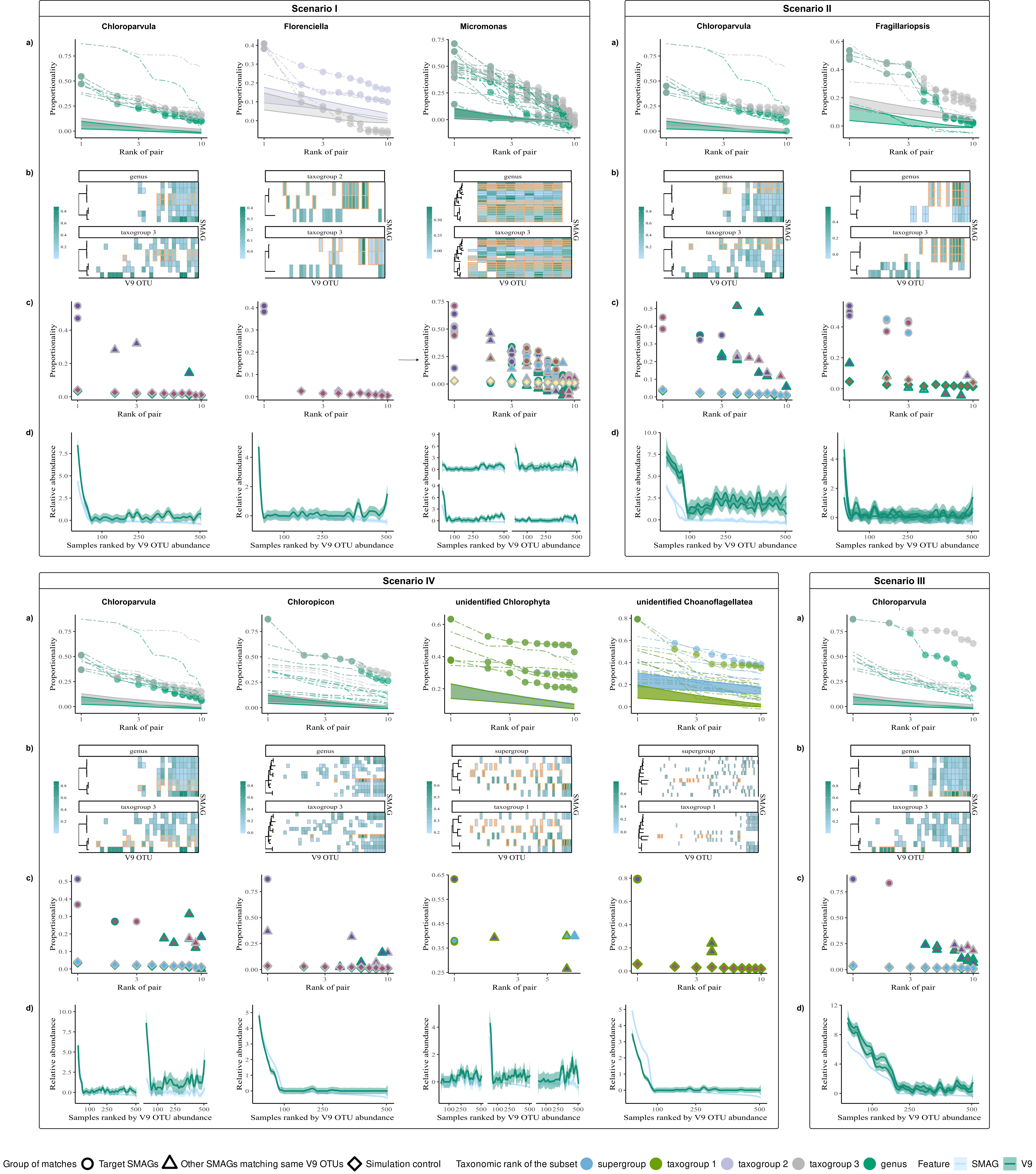

### draft_fig_1_edit3.pdf

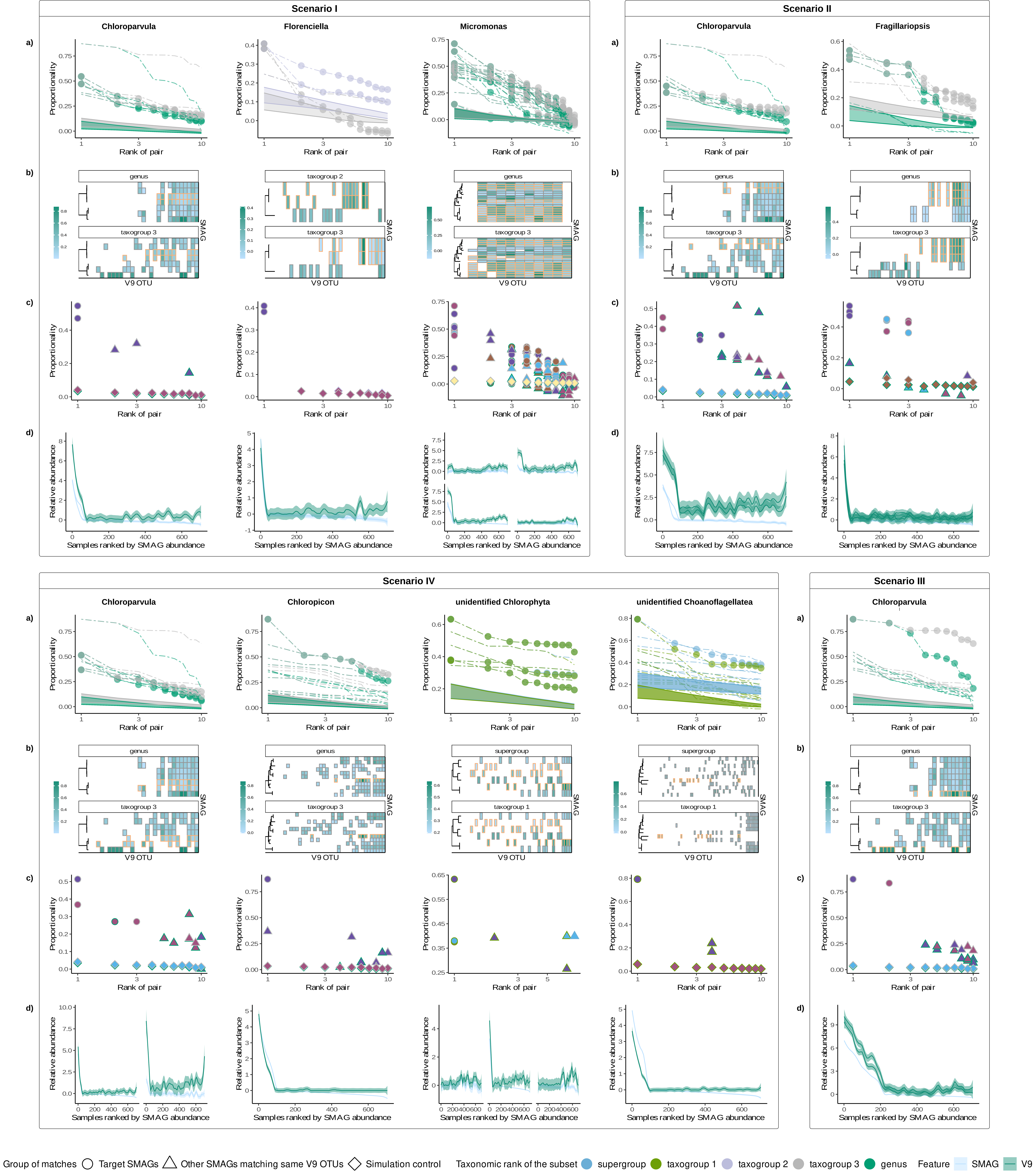

### Figure_1.pdf

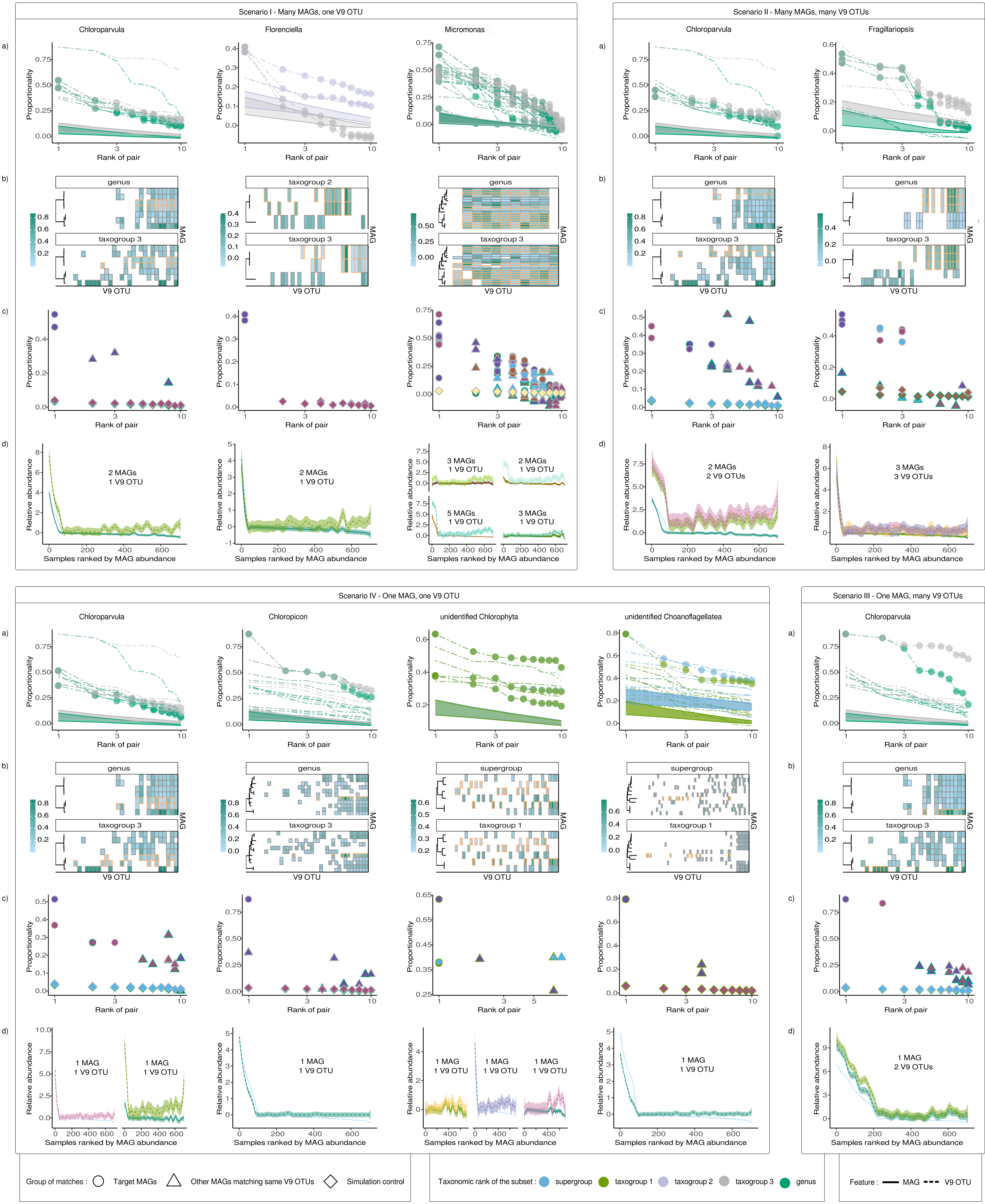

### FINSMAG_control_scatterplots.pdf

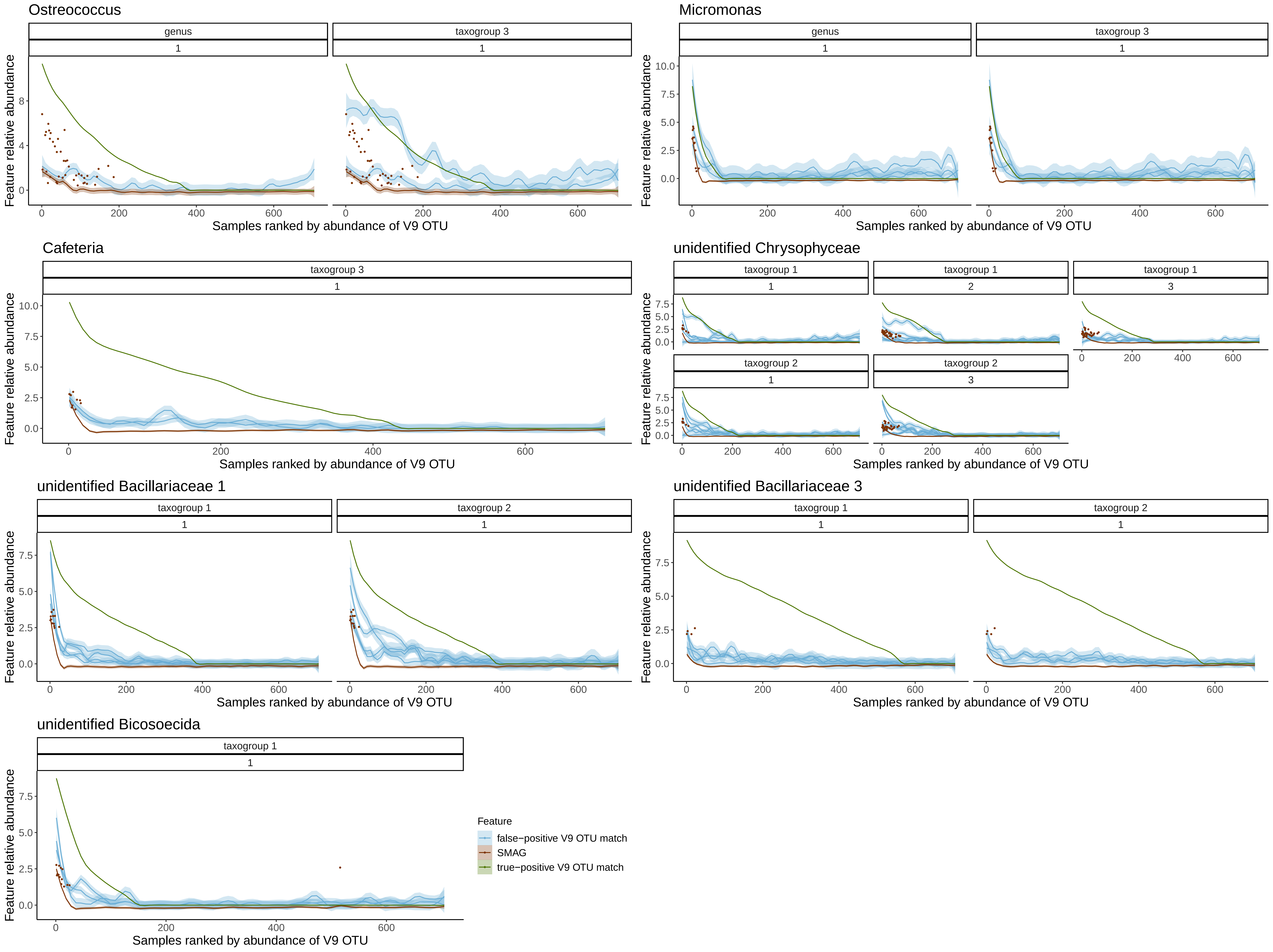

### phi_calc_plots.pdf

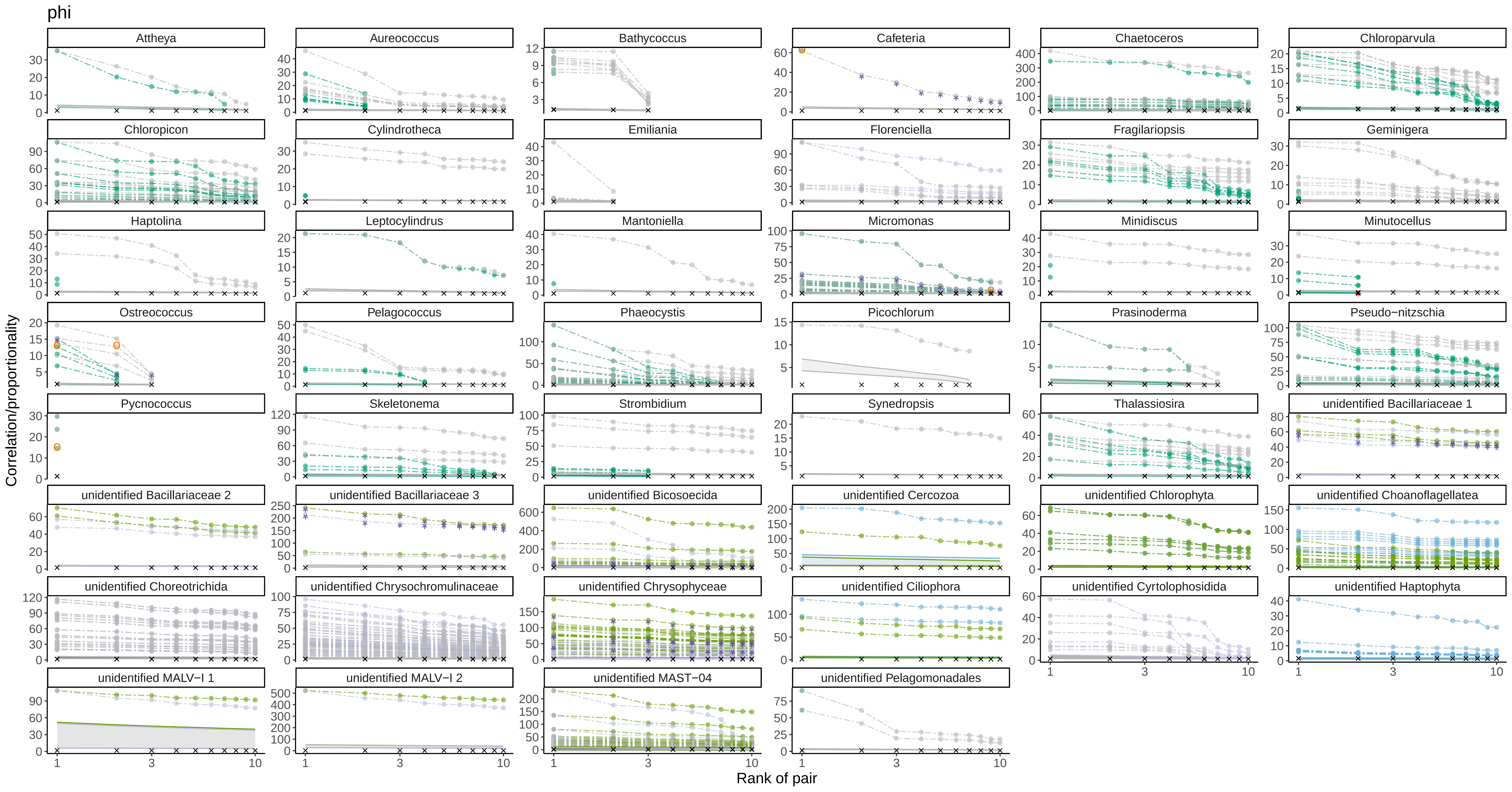

### phi_uniq_facet_plots2.pdf

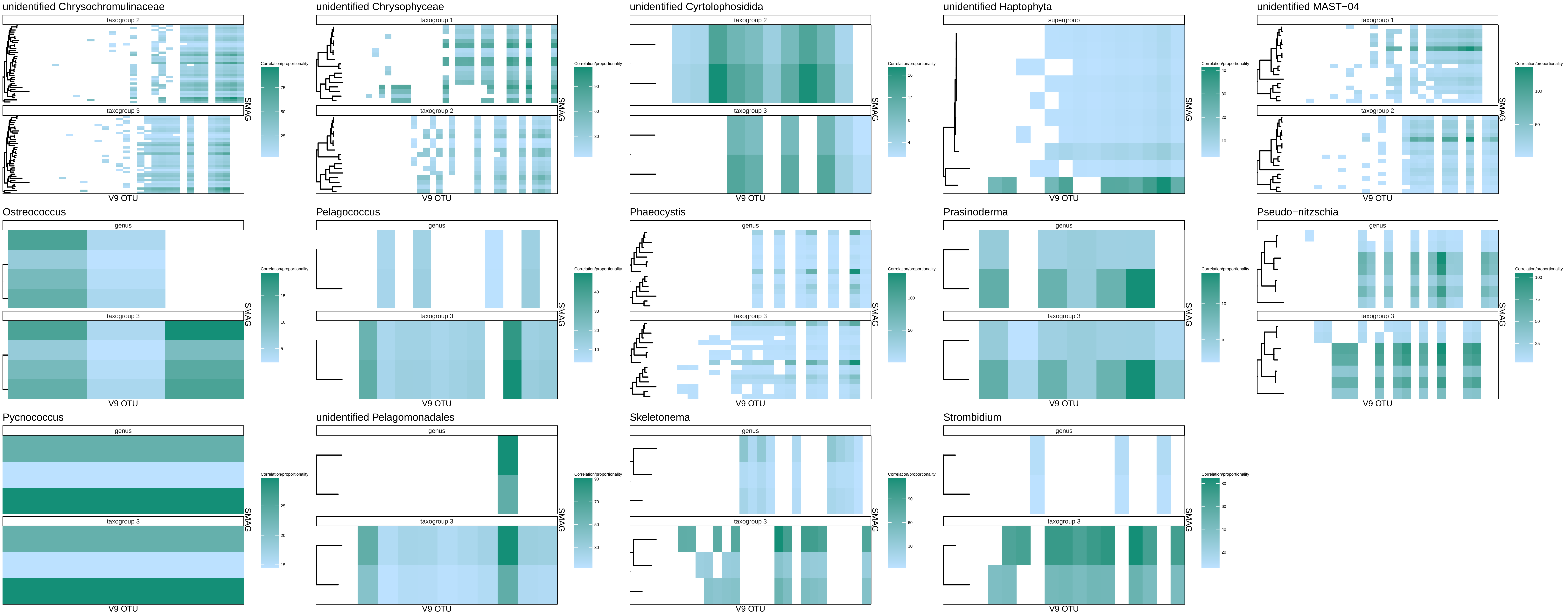

### phi_uniq_facet_plots.pdf

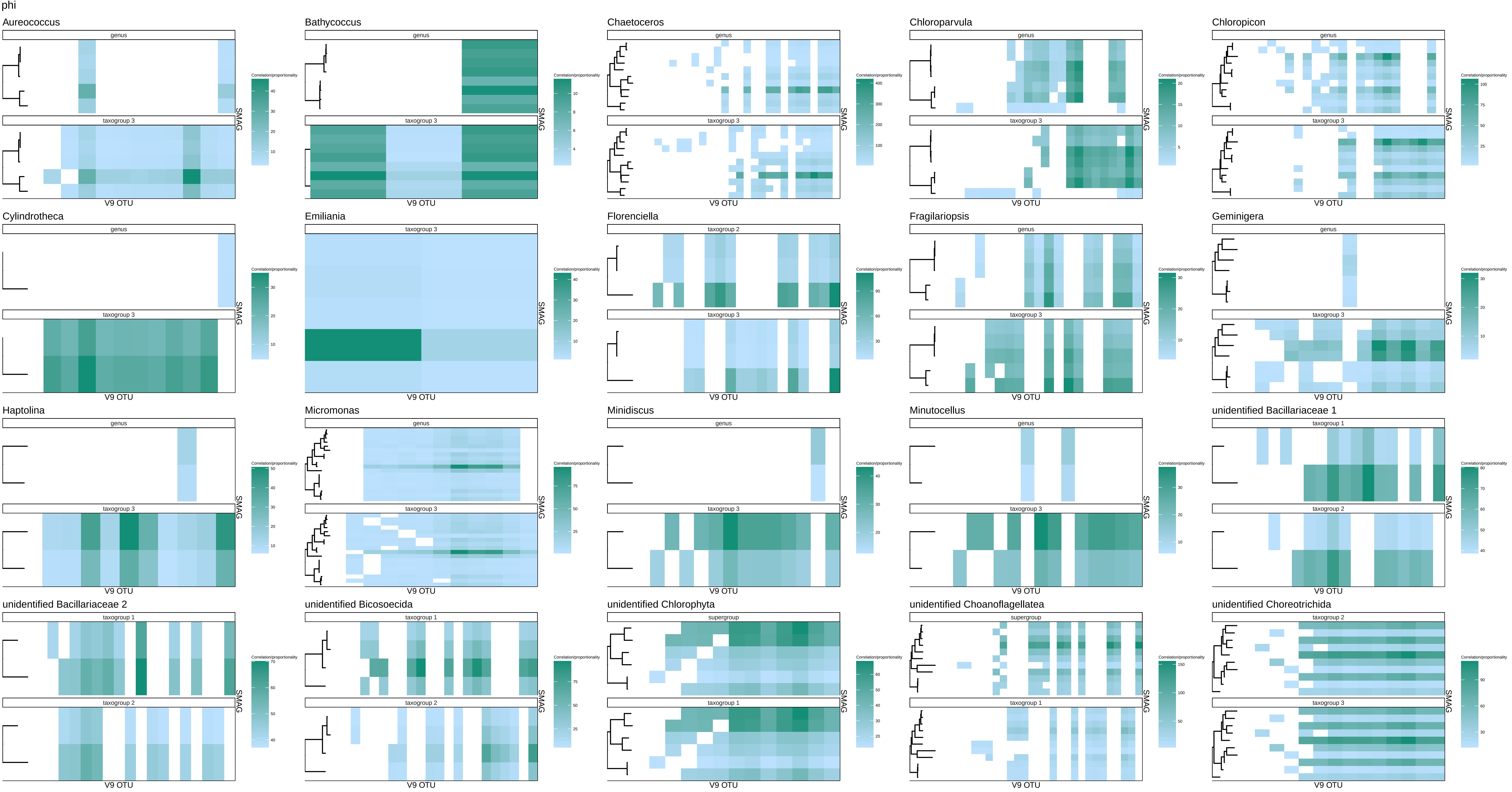

### phs_calc_plots.pdf

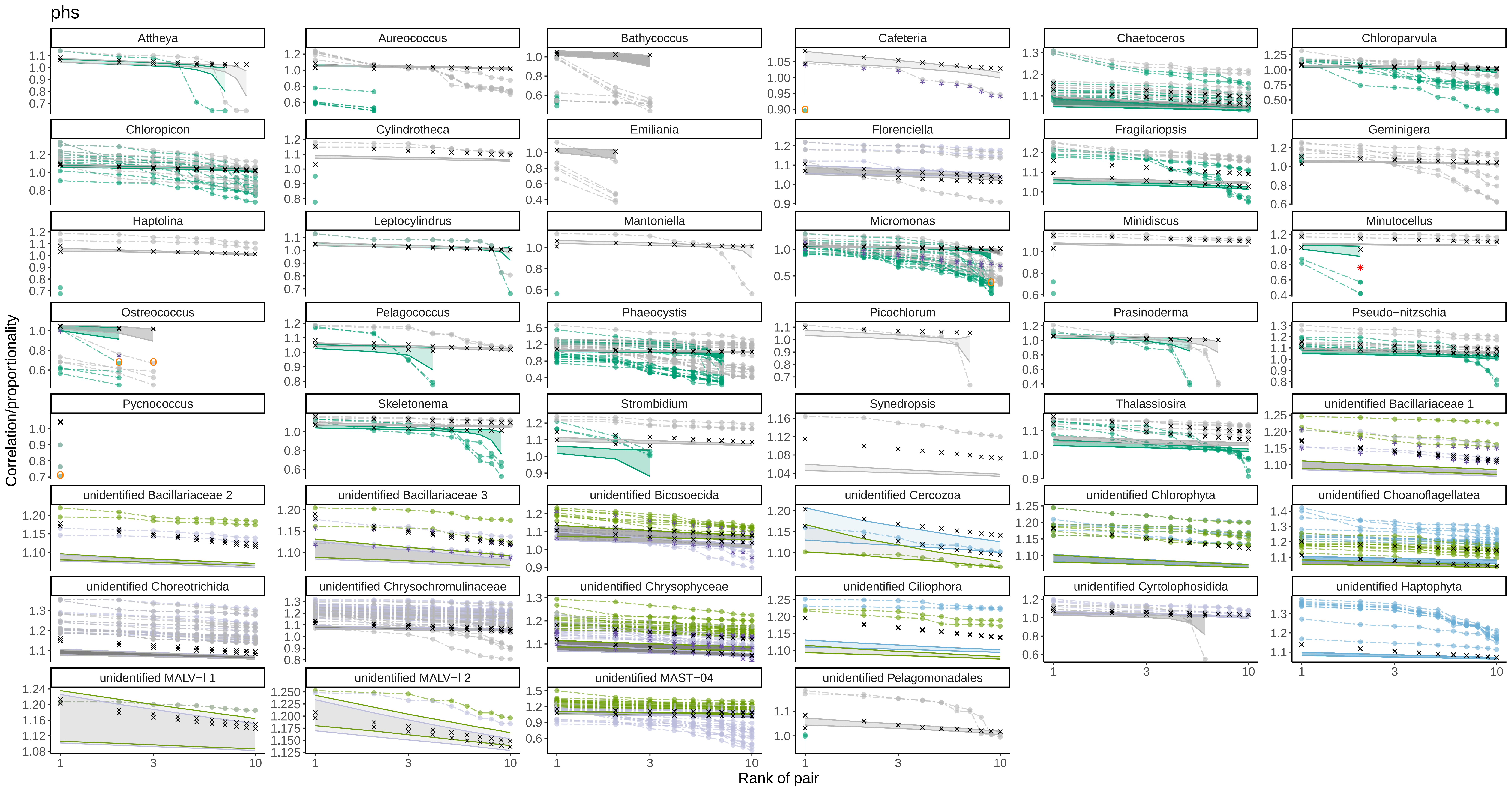

### phs_uniq_facet_plots2.pdf

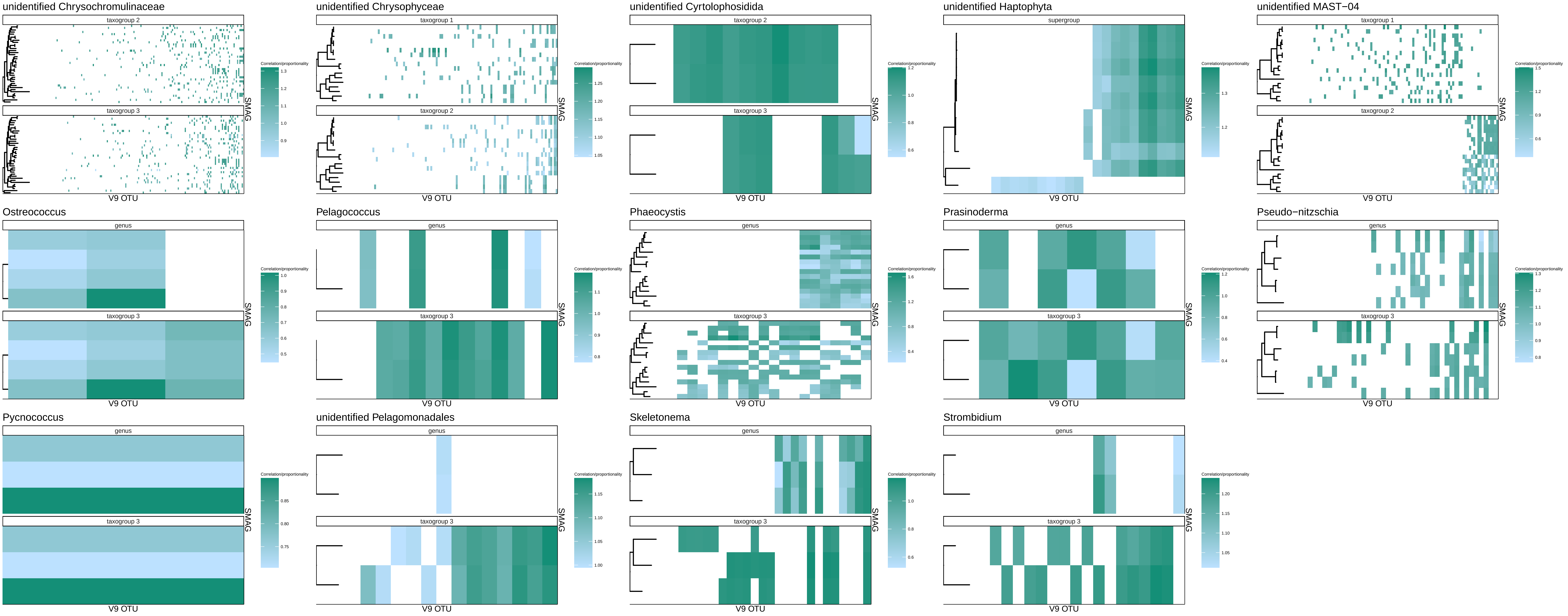

### phs_uniq_facet_plots.pdf

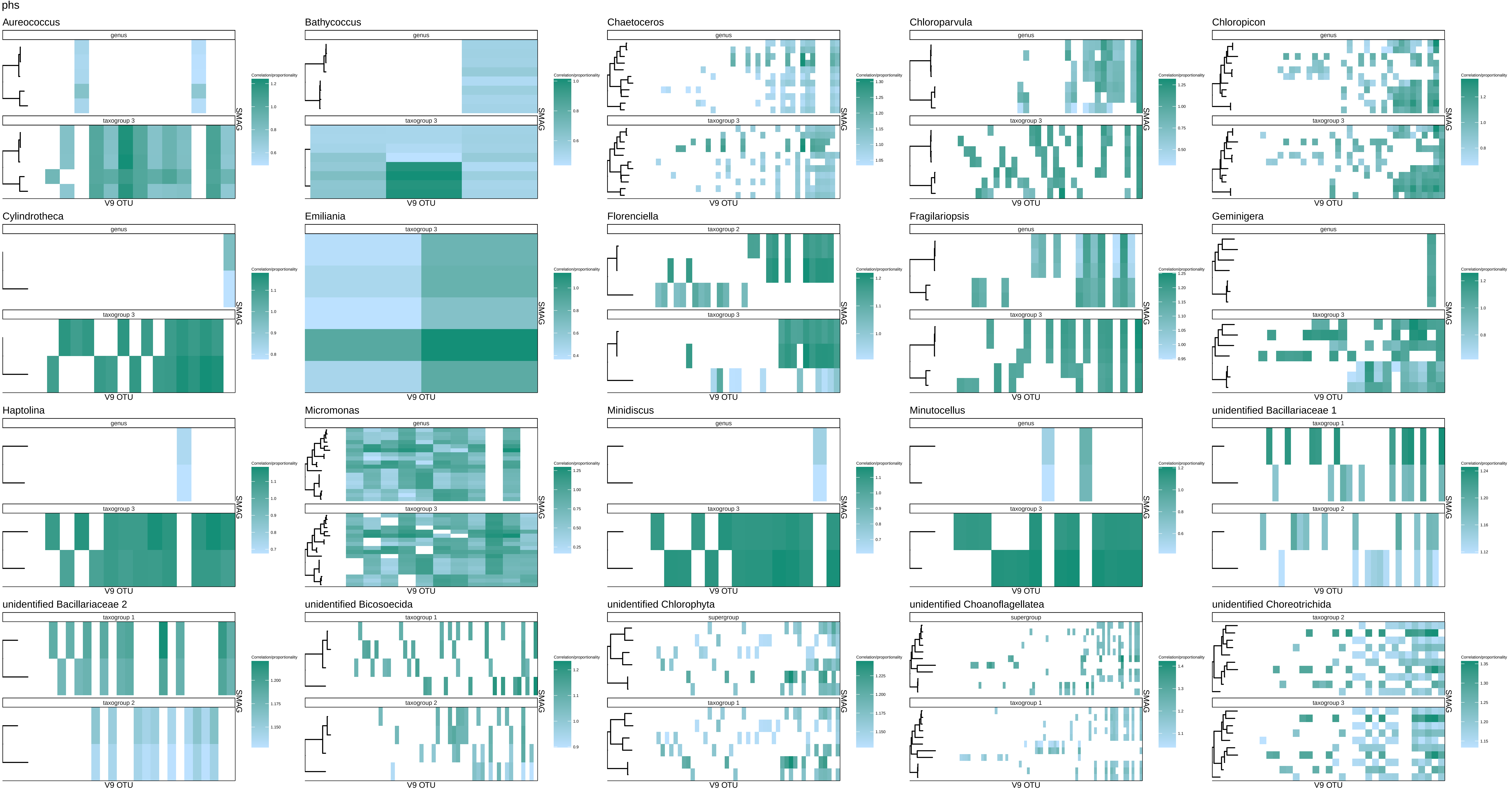

### rho_calc_plots.pdf

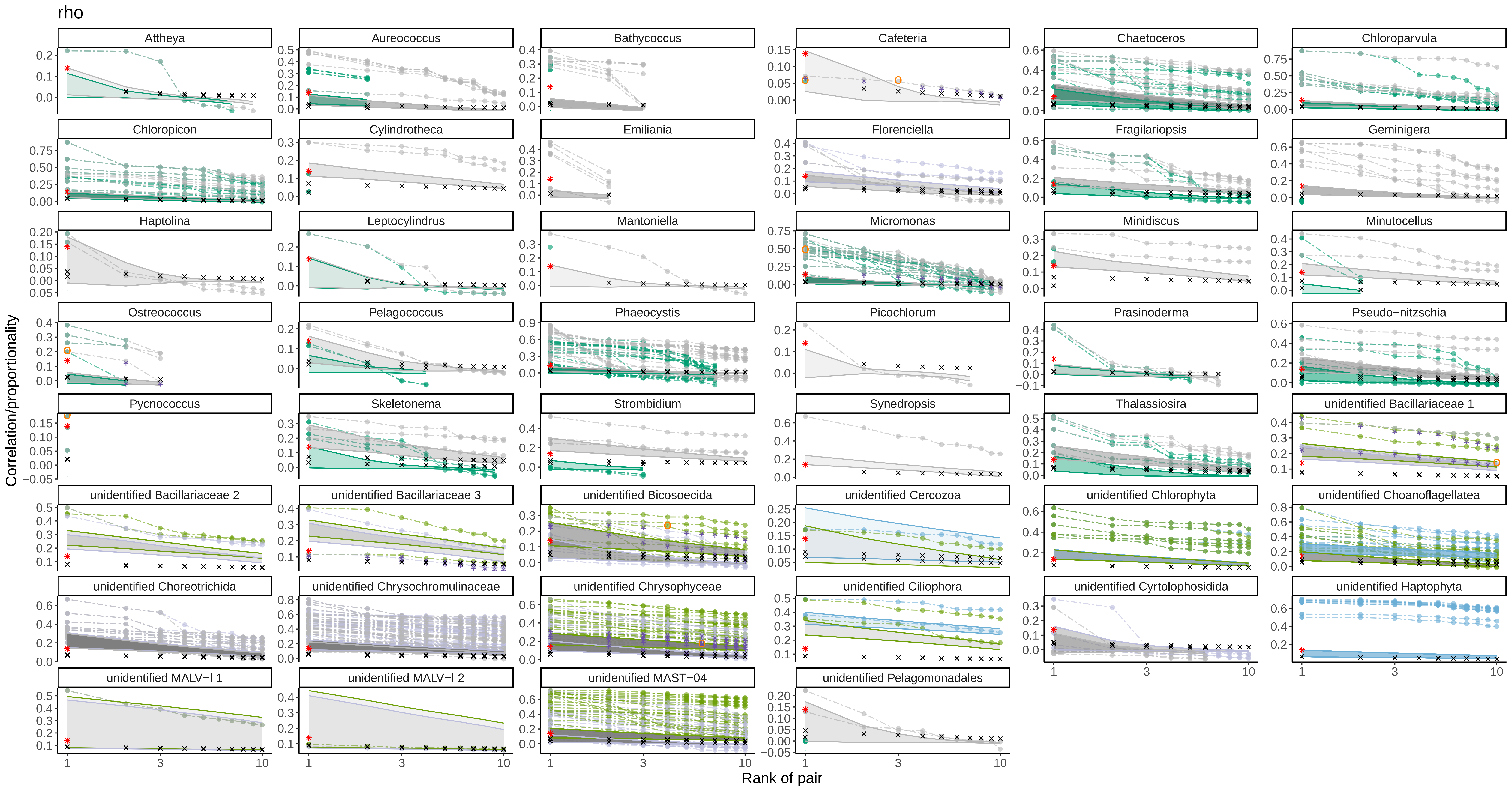

### rho_uniq_facet_plots2.pdf

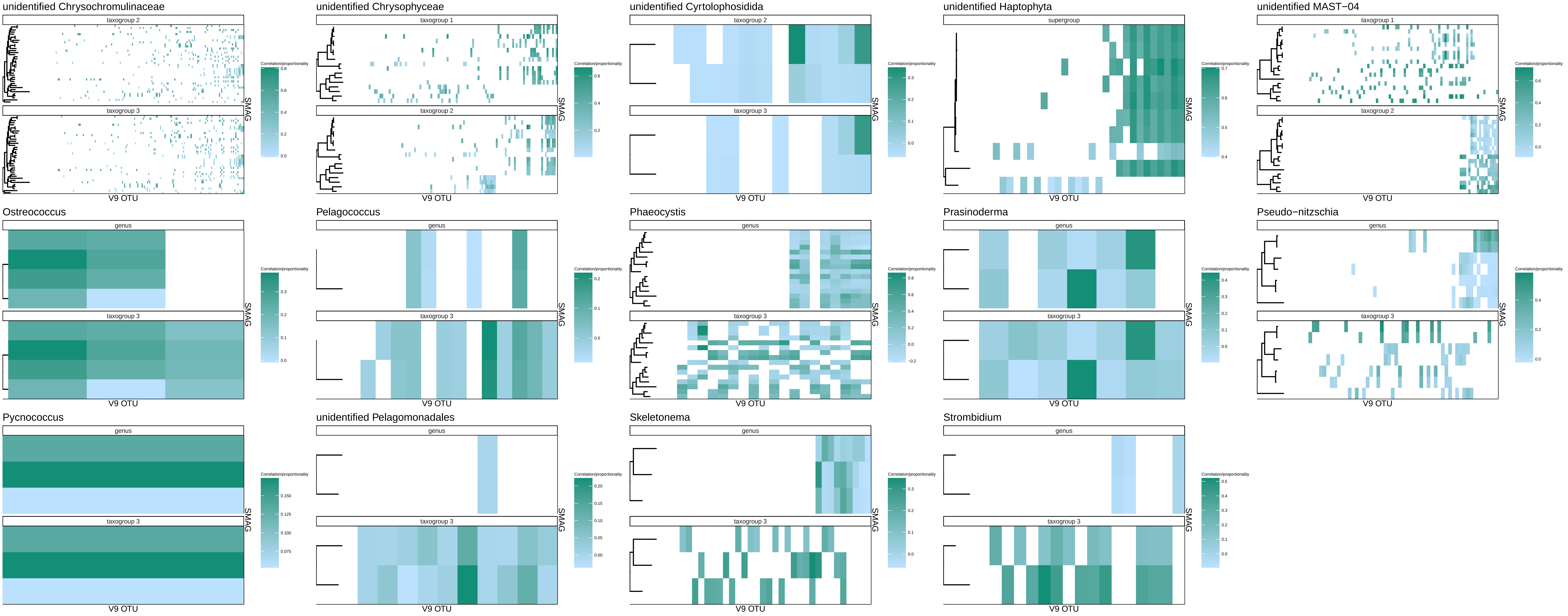

### rho_uniq_facet_plots.pdf

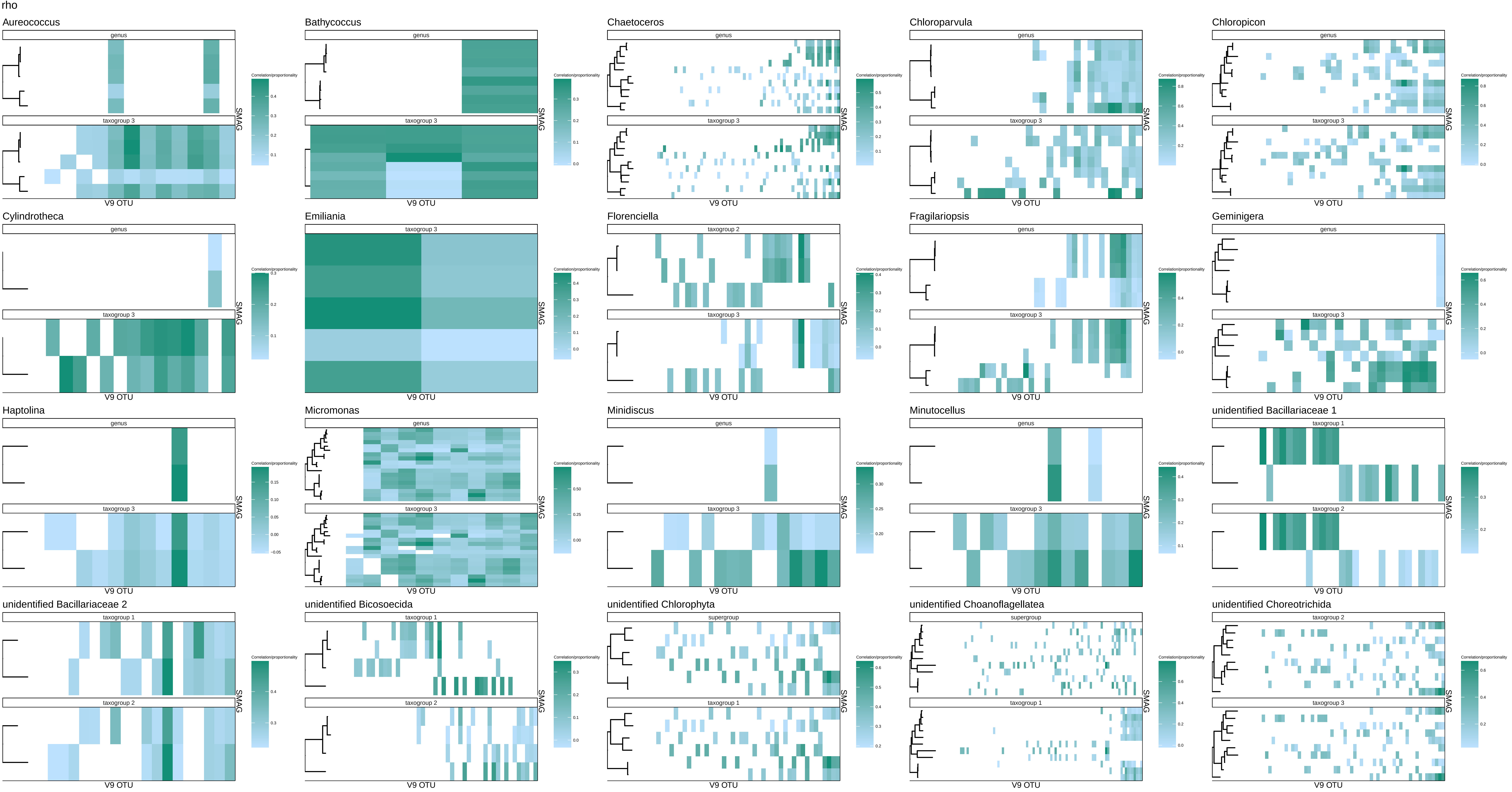

### rhoSMAG_control_scatterplots.pdf

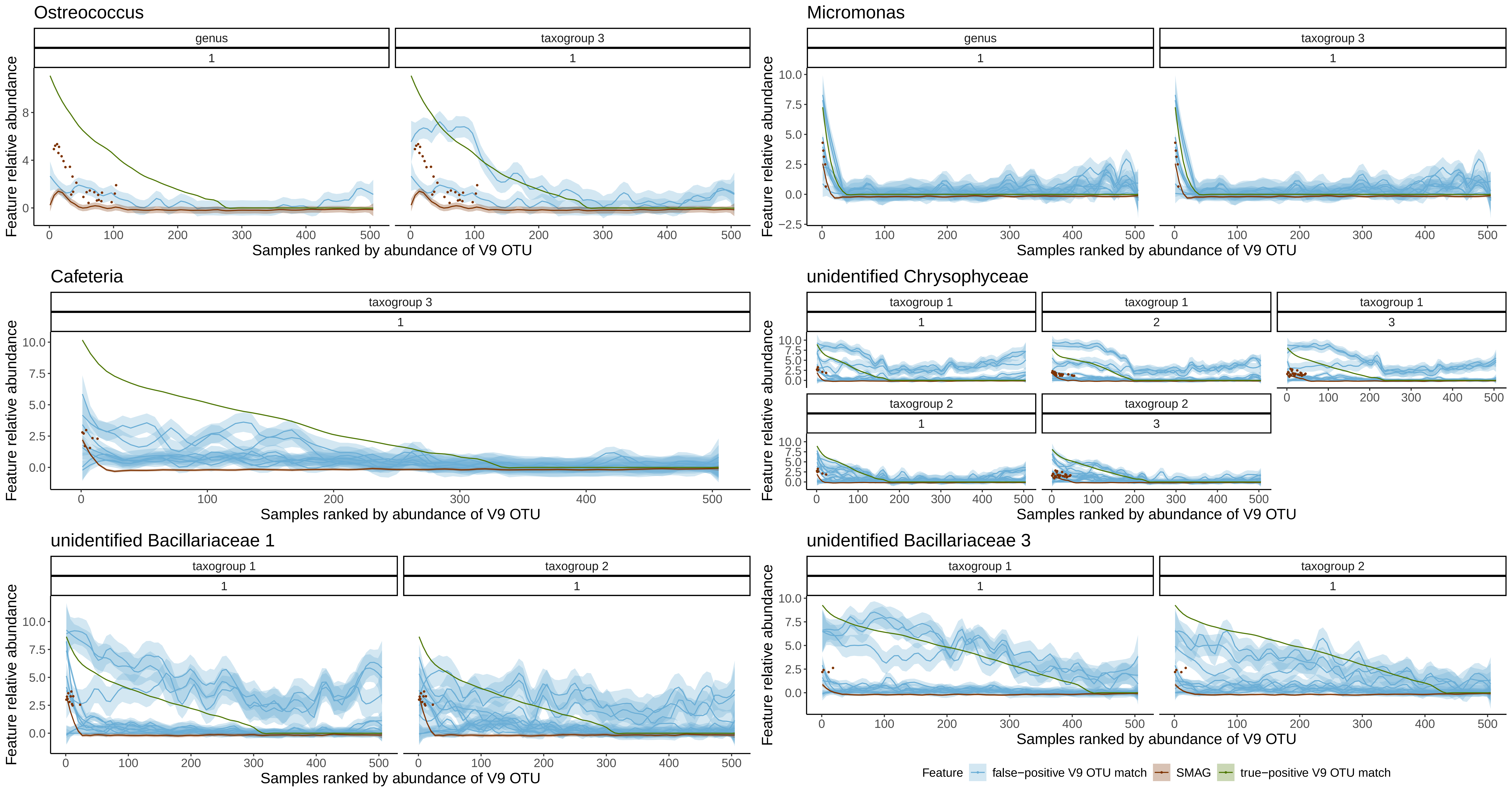

### shuffle_se_cor.pdf

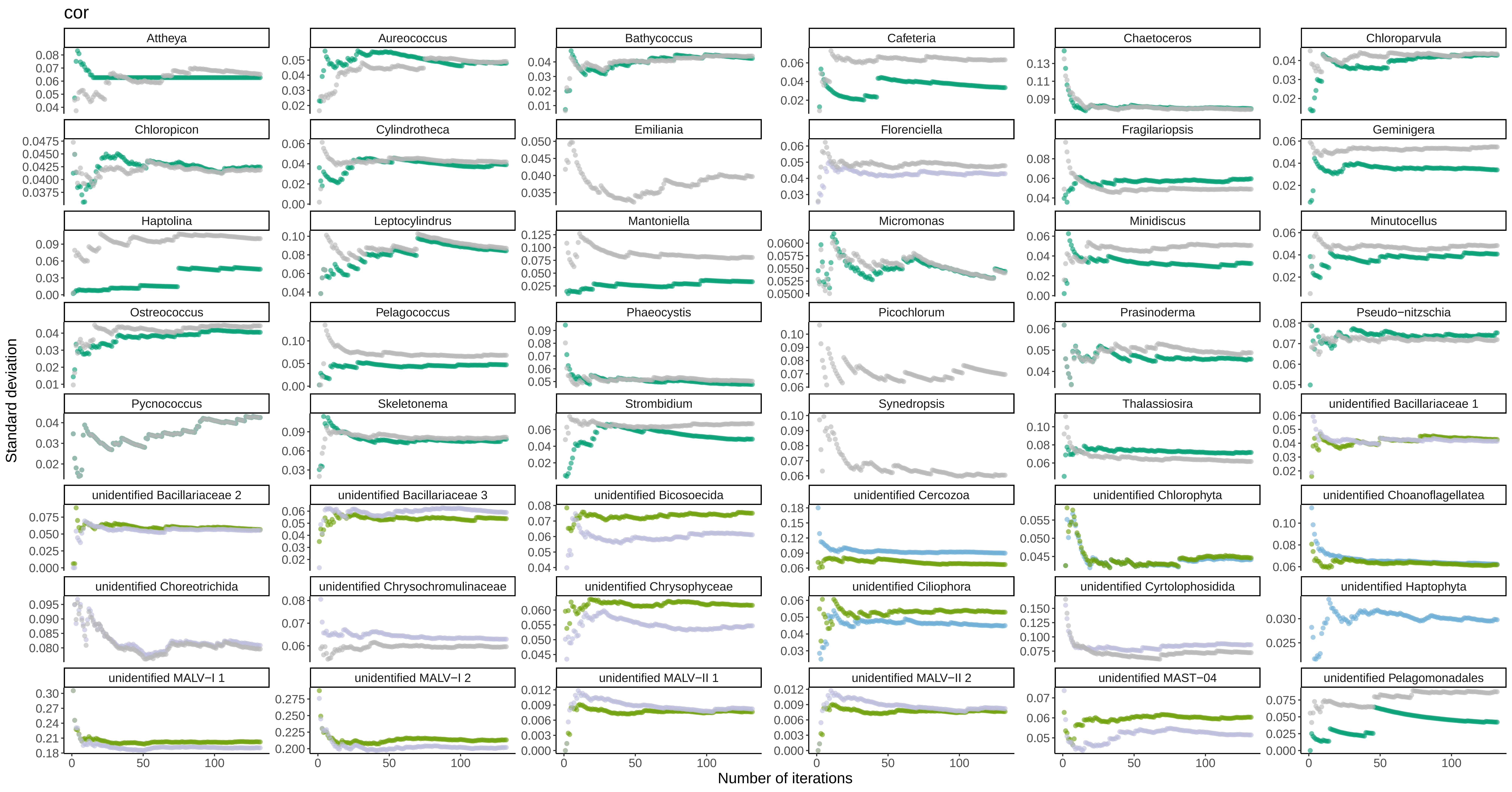

### shuffle_se_phi.pdf

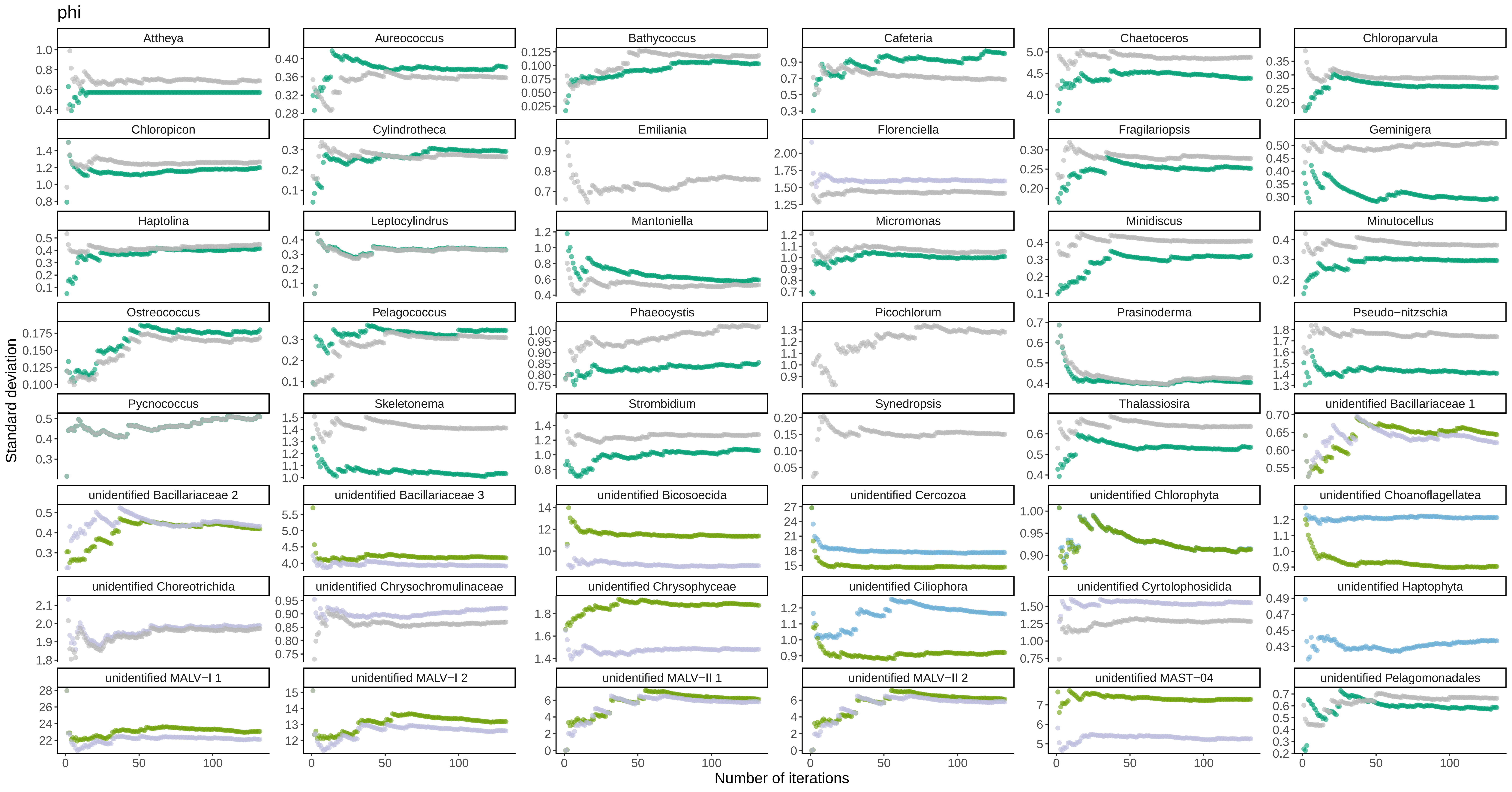

### shuffle_se_phs.pdf

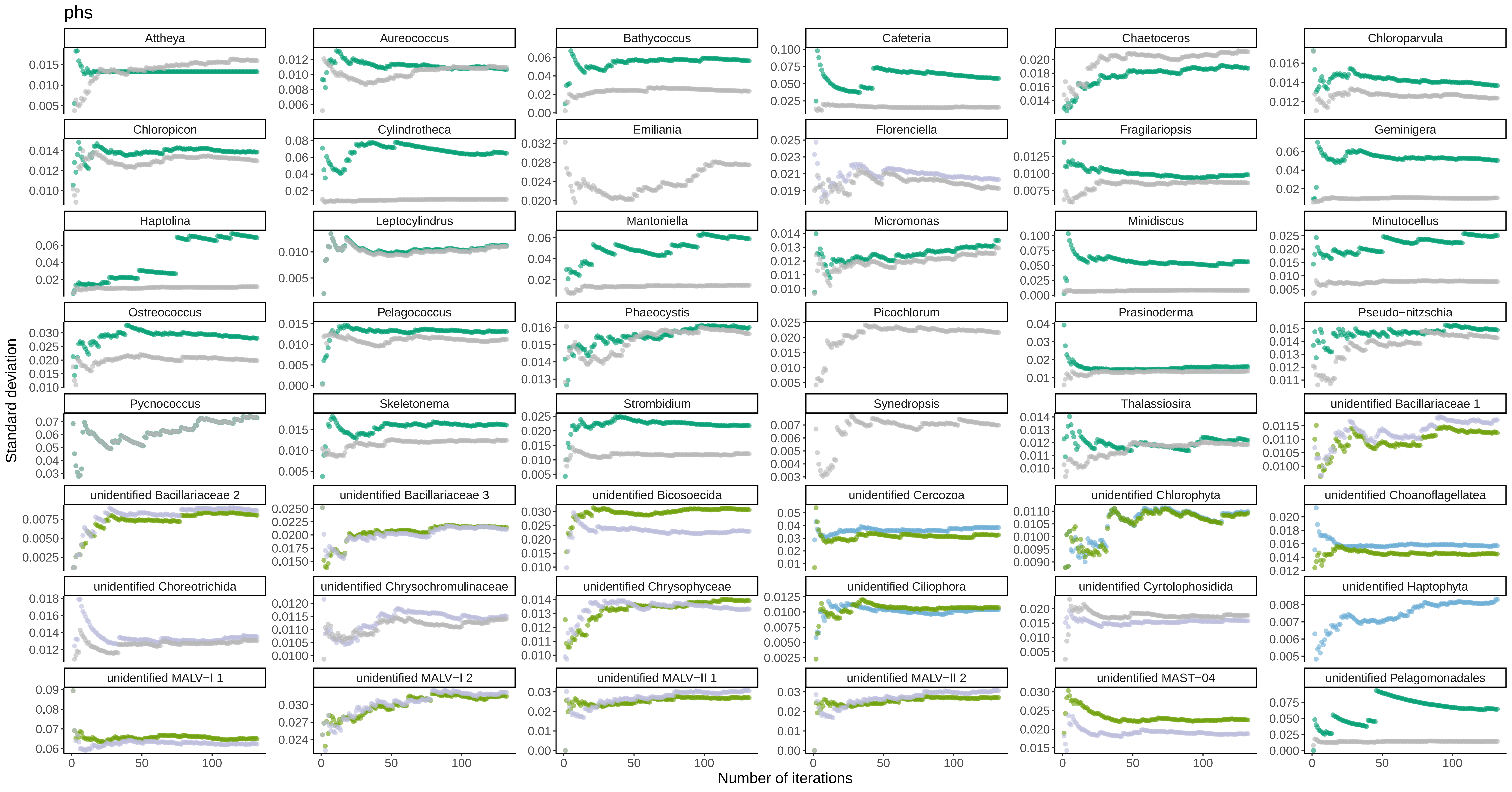

### shuffle_se_rho.pdf

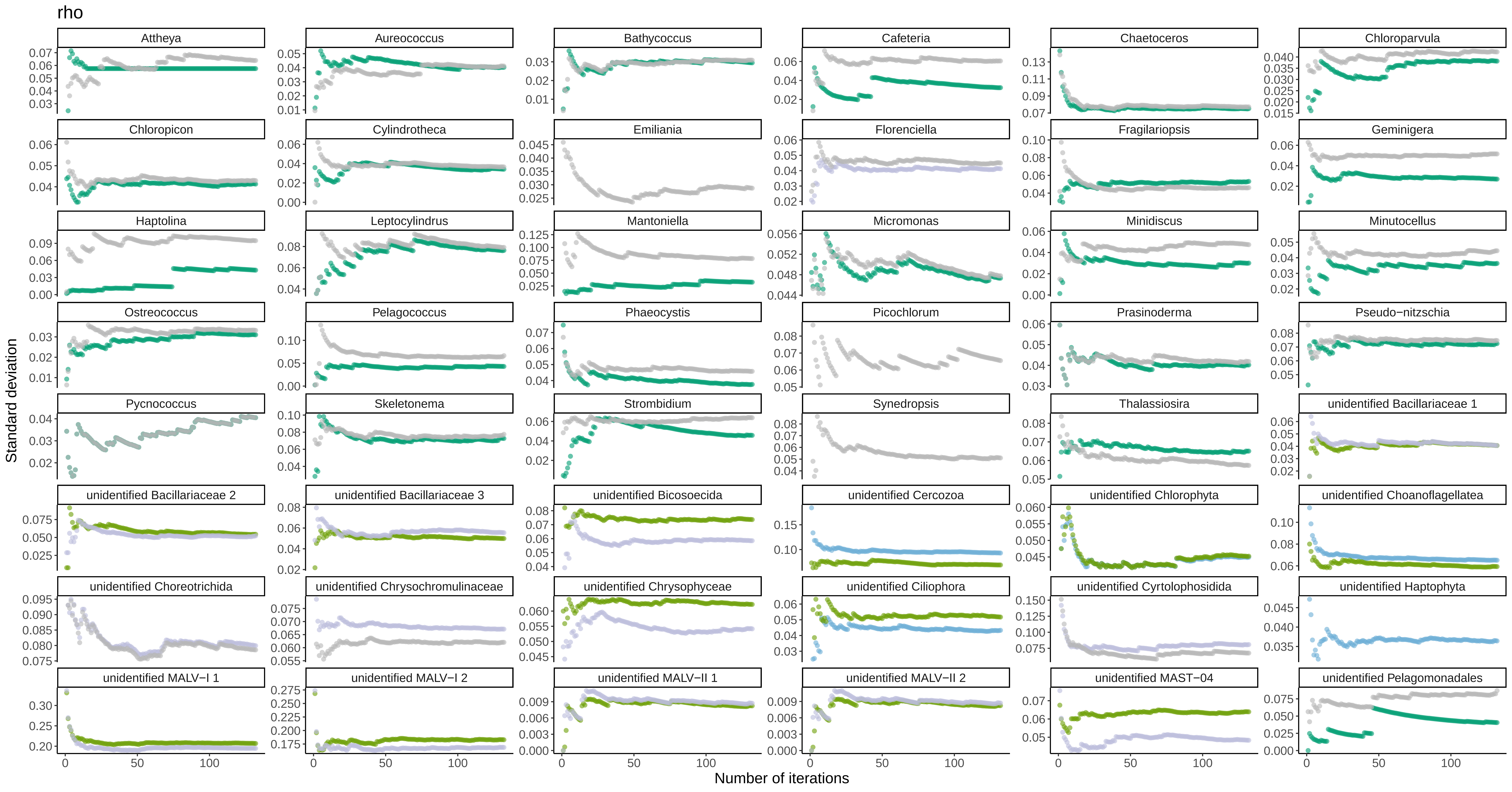

### shuffle_se_vlr.pdf

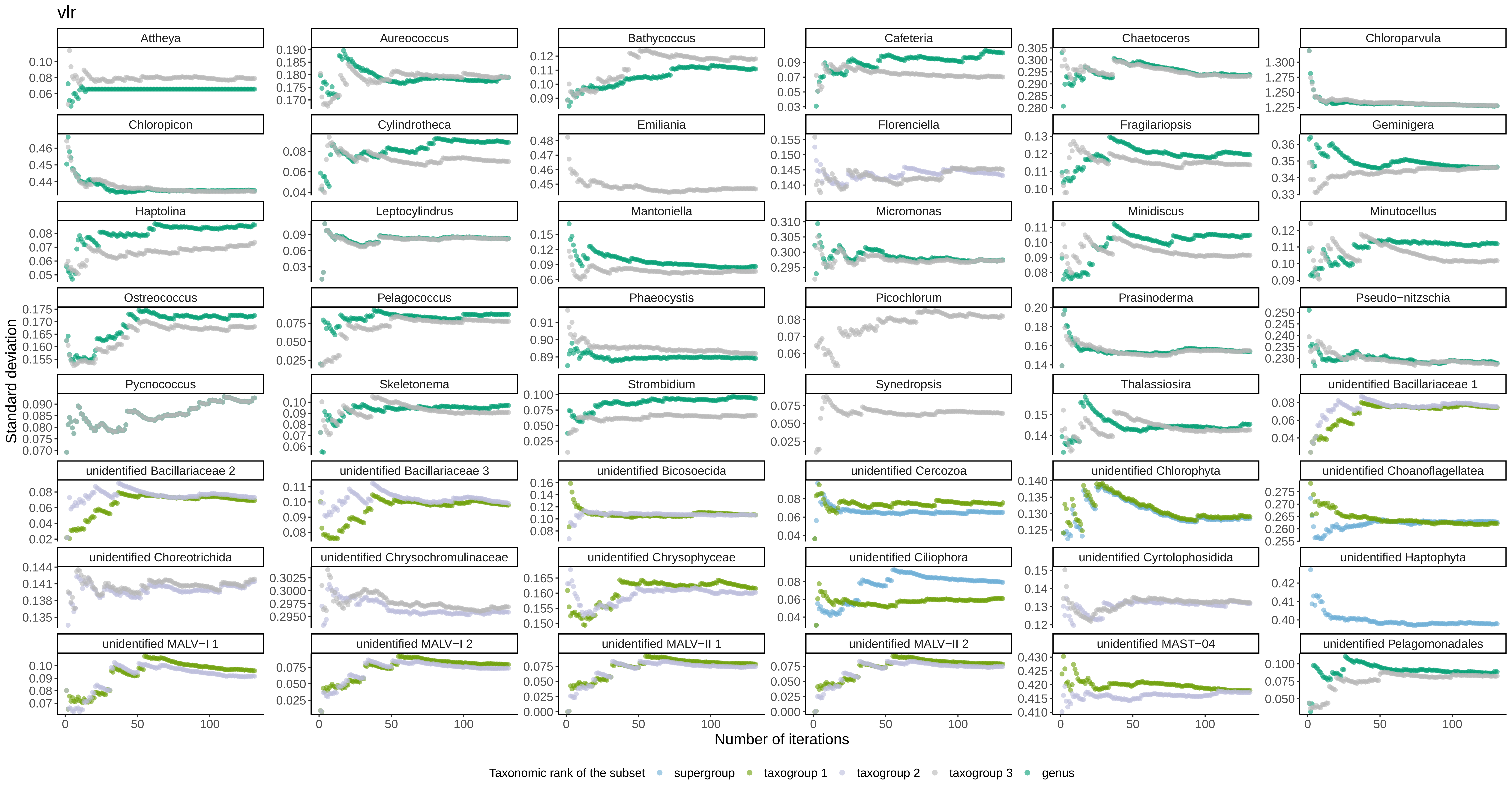

### sim_matchtype_counts_cor.pdf

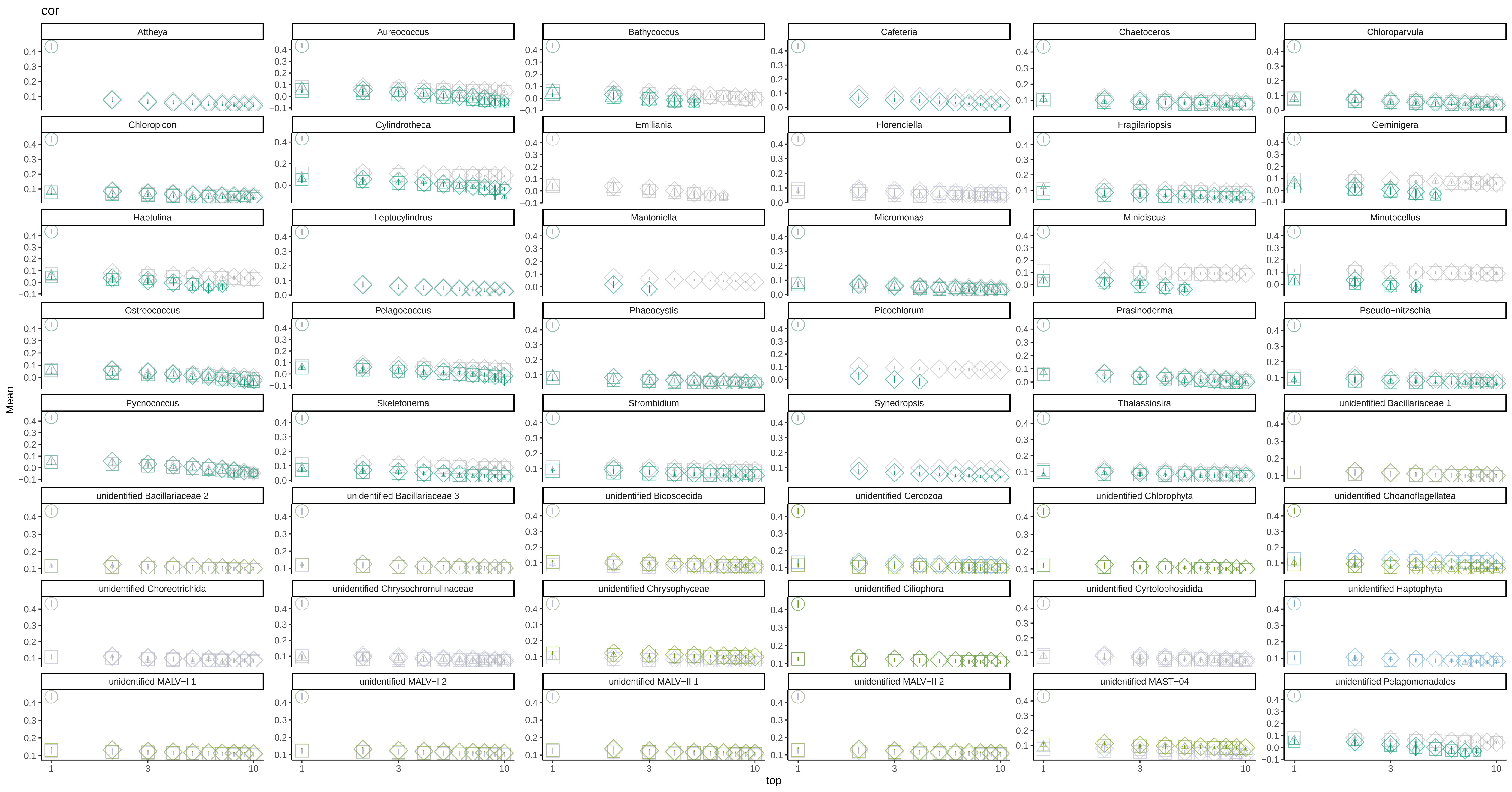

### sim_matchtype_counts_phi.pdf

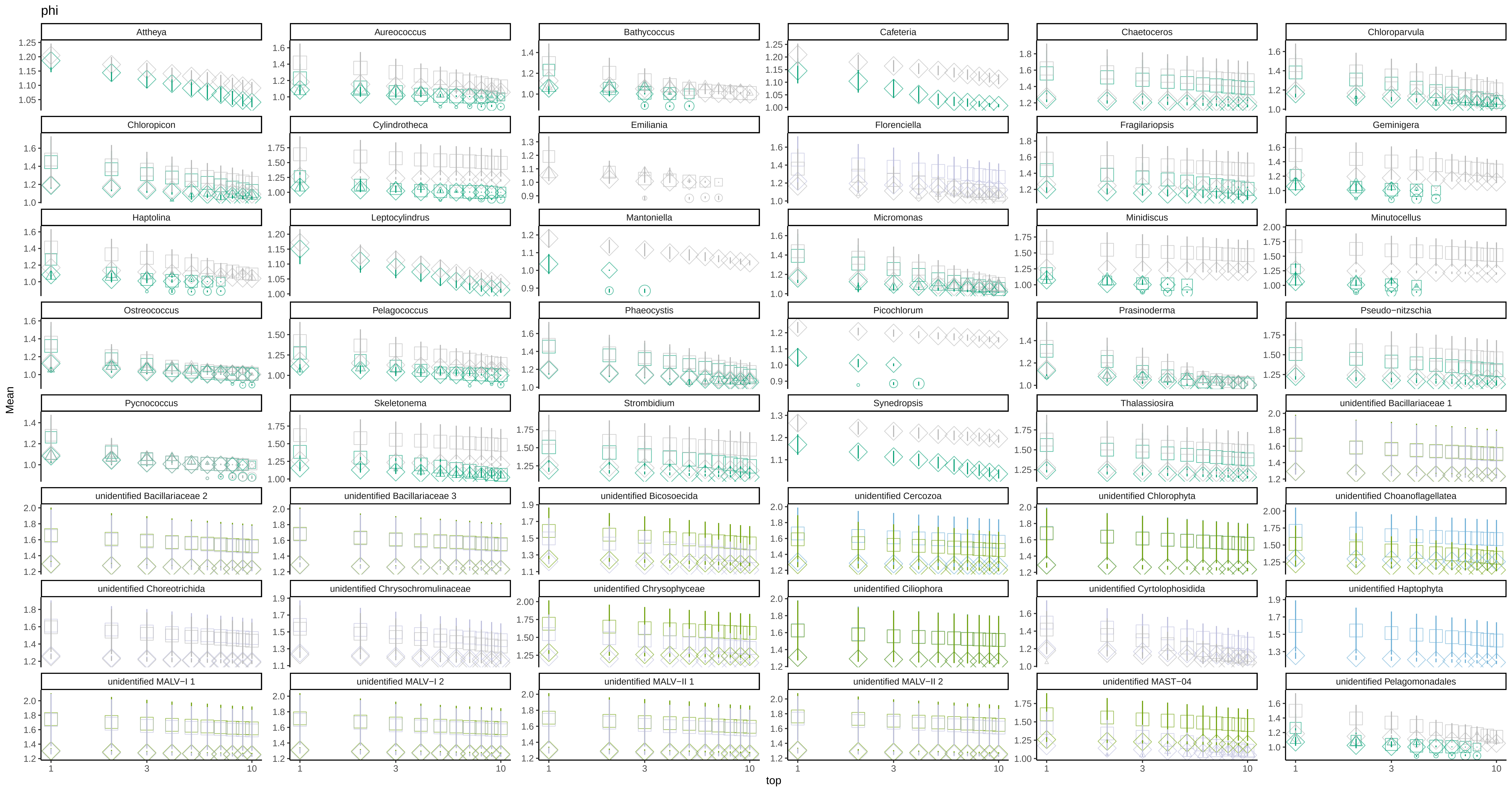

### sim_matchtype_counts_phs.pdf

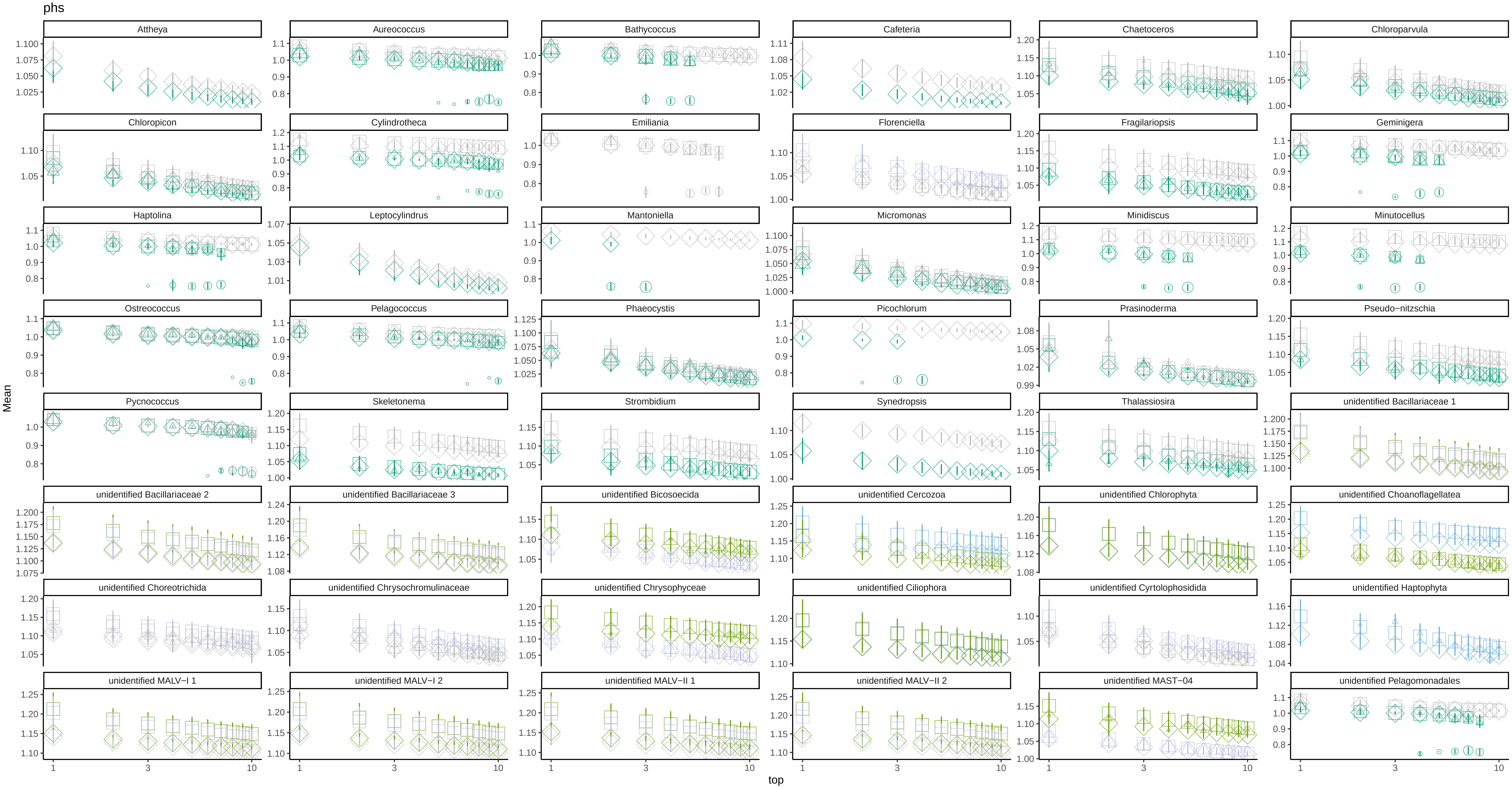

### sim_matchtype_counts_rho.pdf

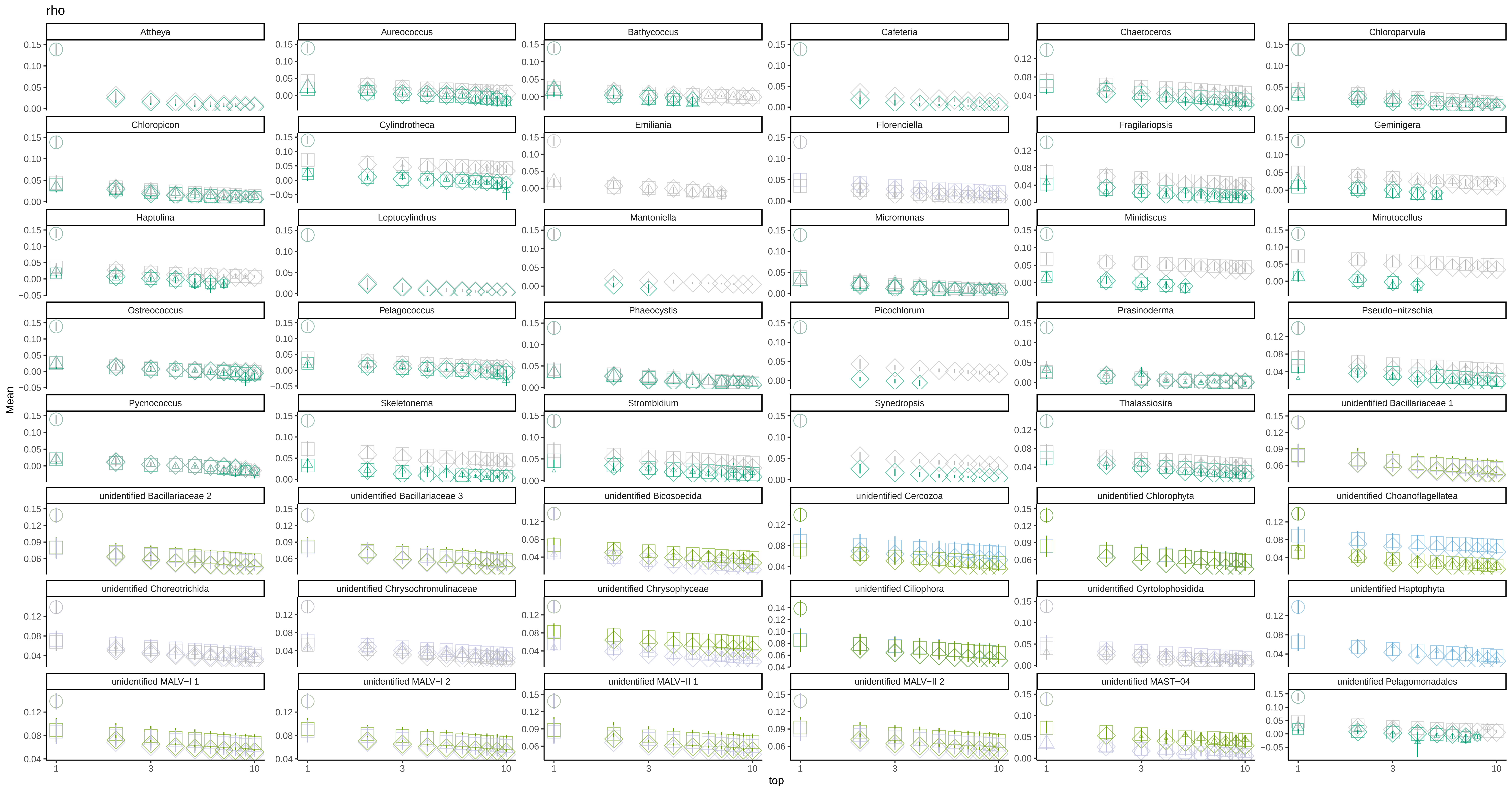

### sim_matchtype_counts_vlr.pdf

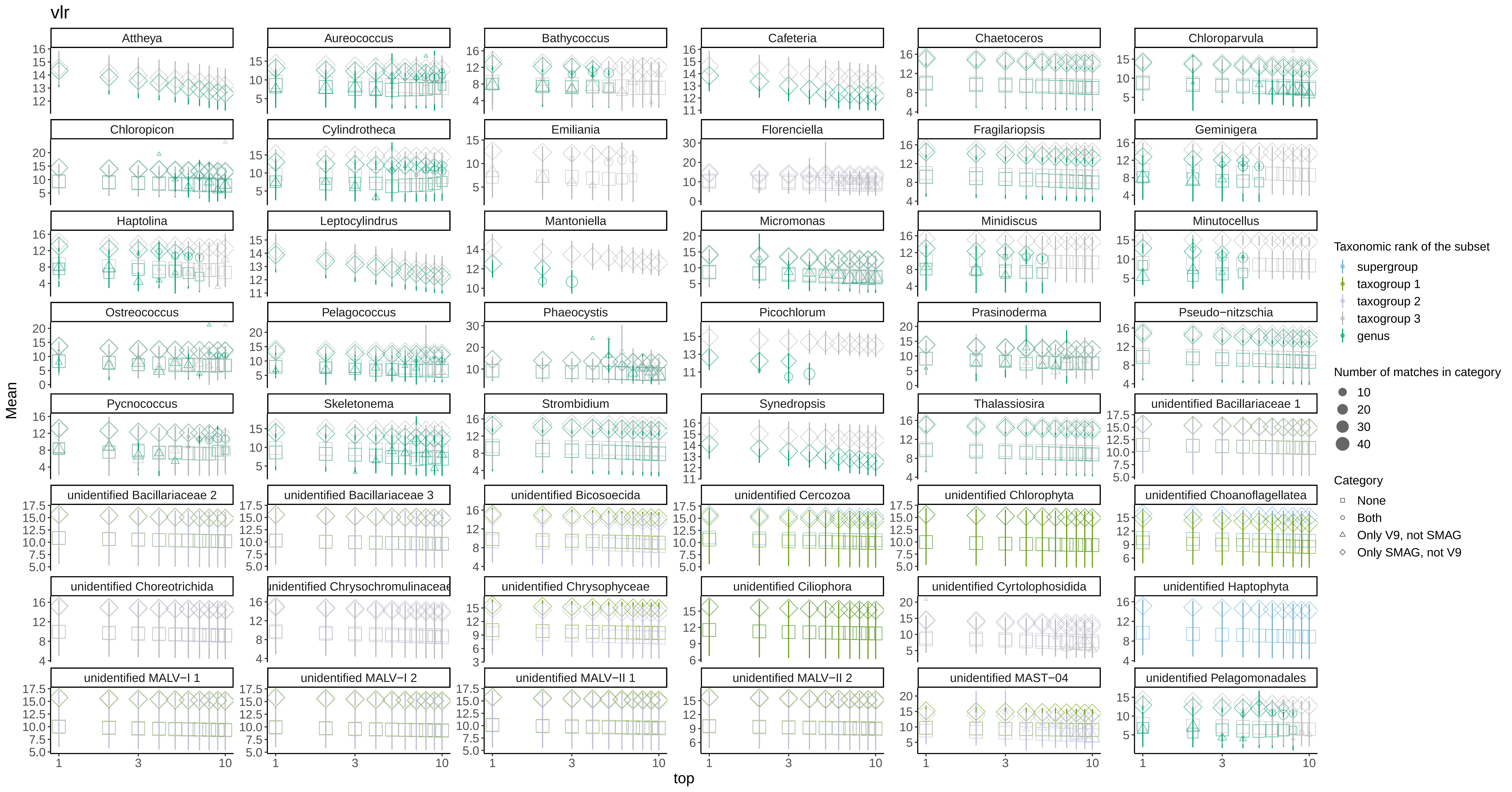

### sim_se_cor.pdf

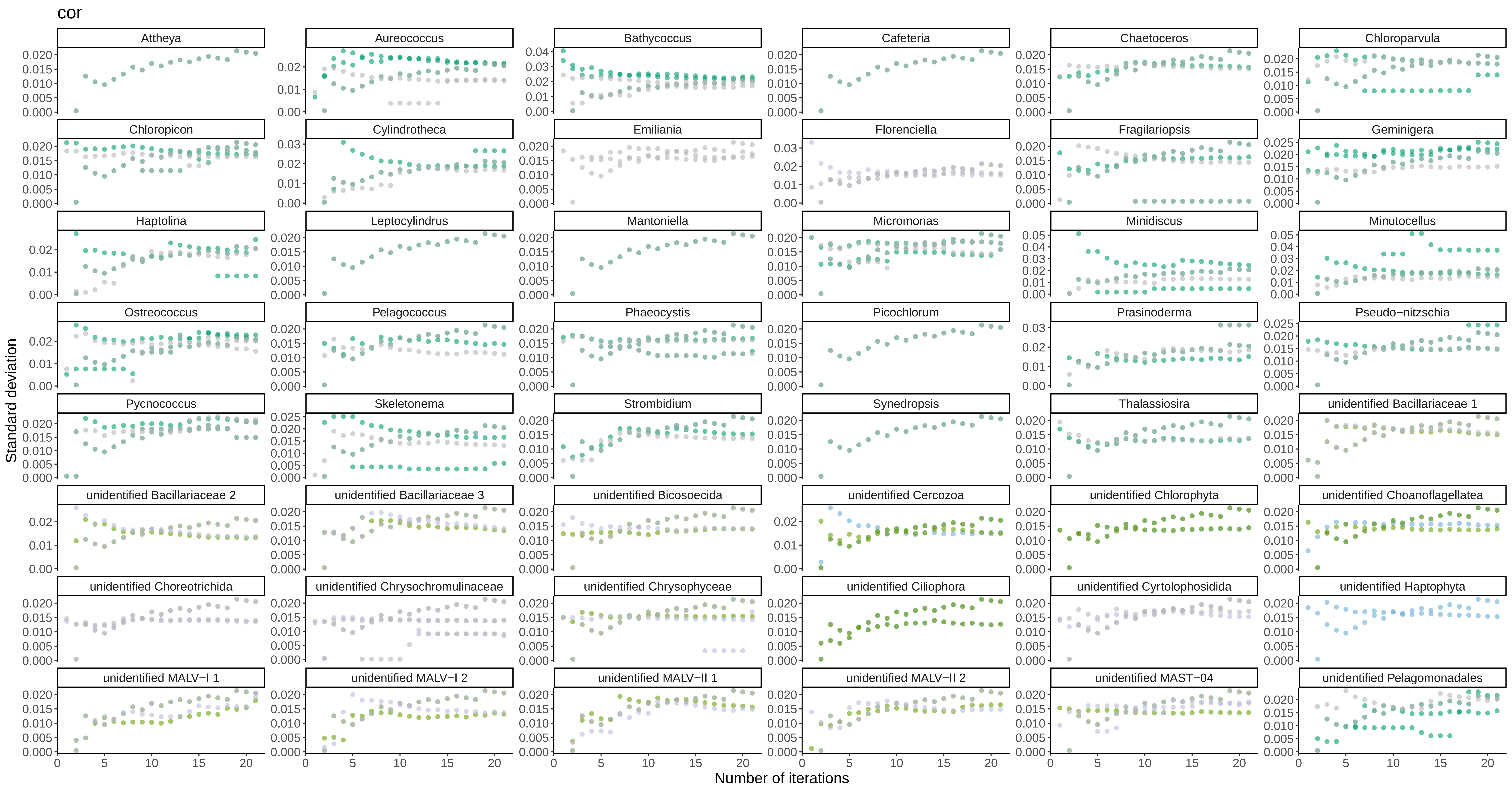

### sim_se_phi.pdf

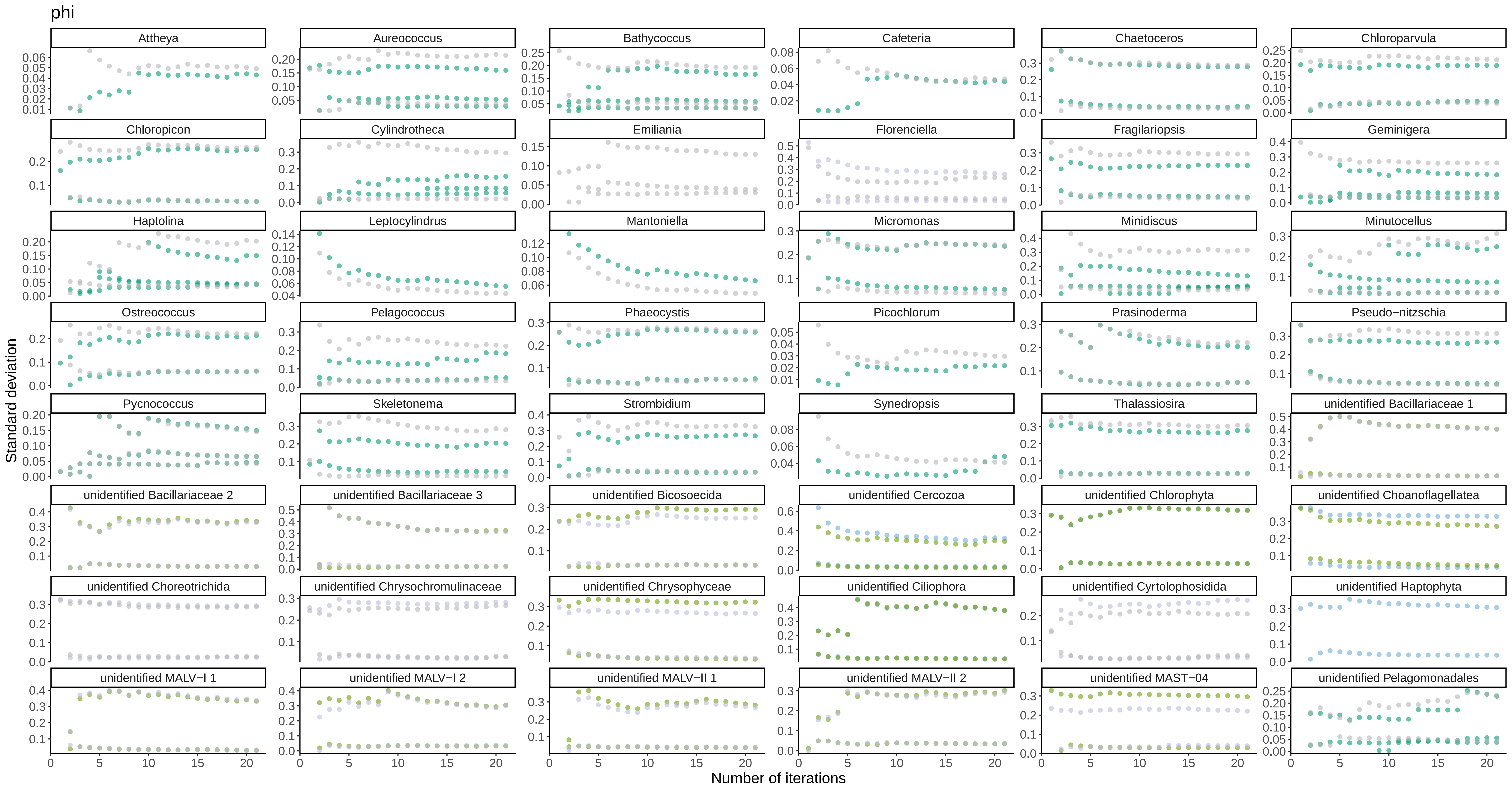

### sim_se_phs.pdf

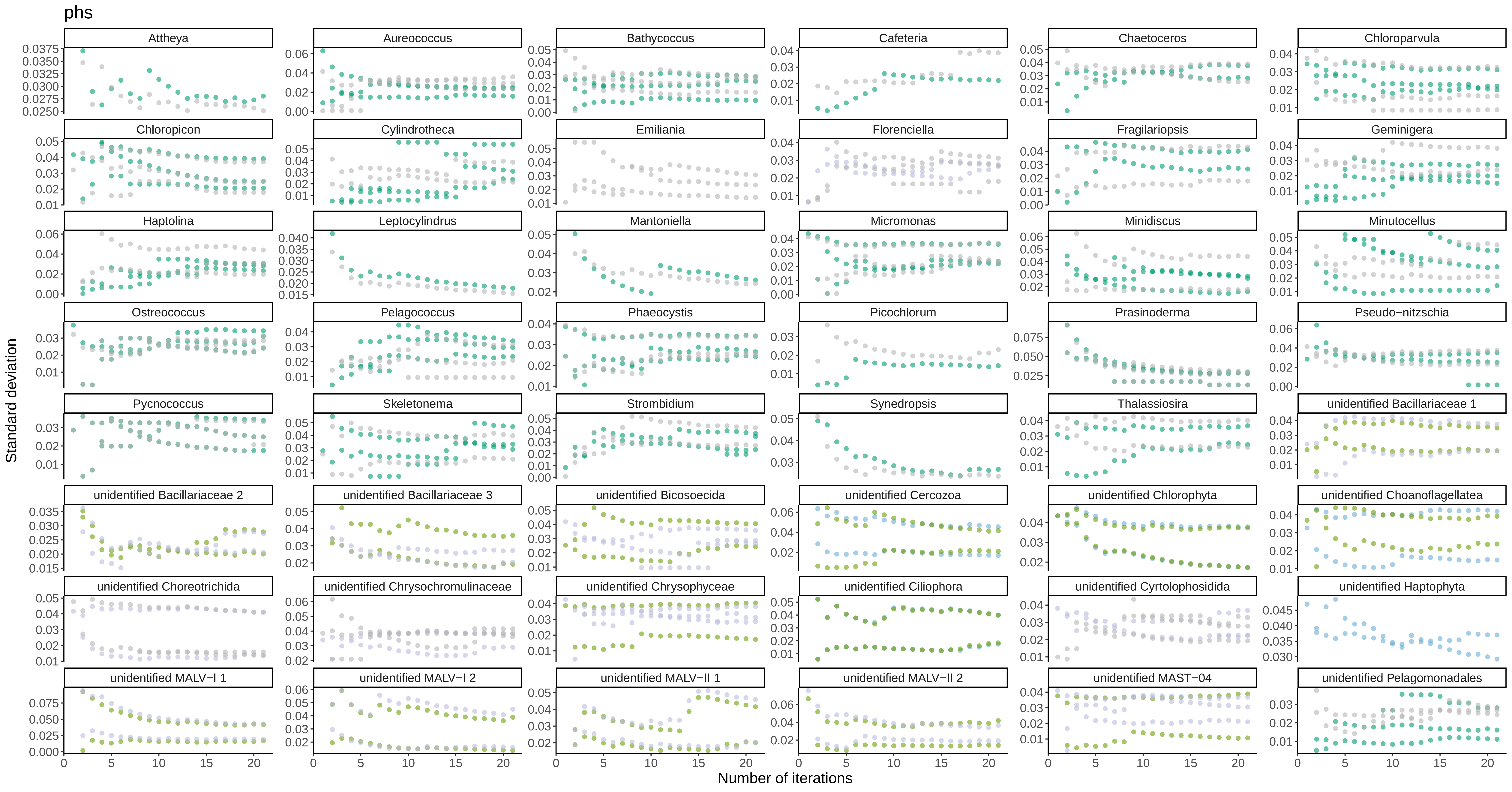

### sim_se_rho.pdf

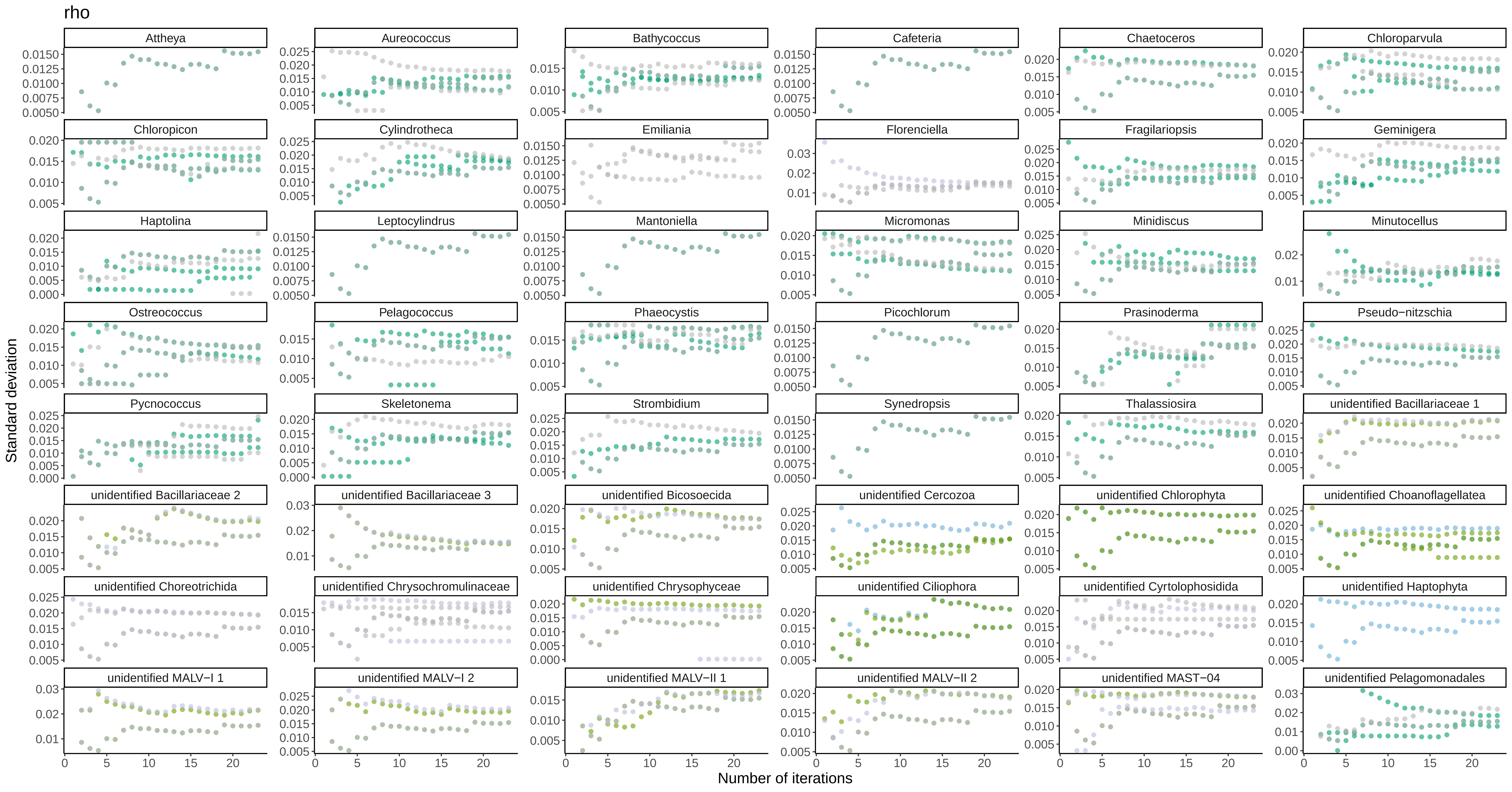

### sim_se_vlr.pdf

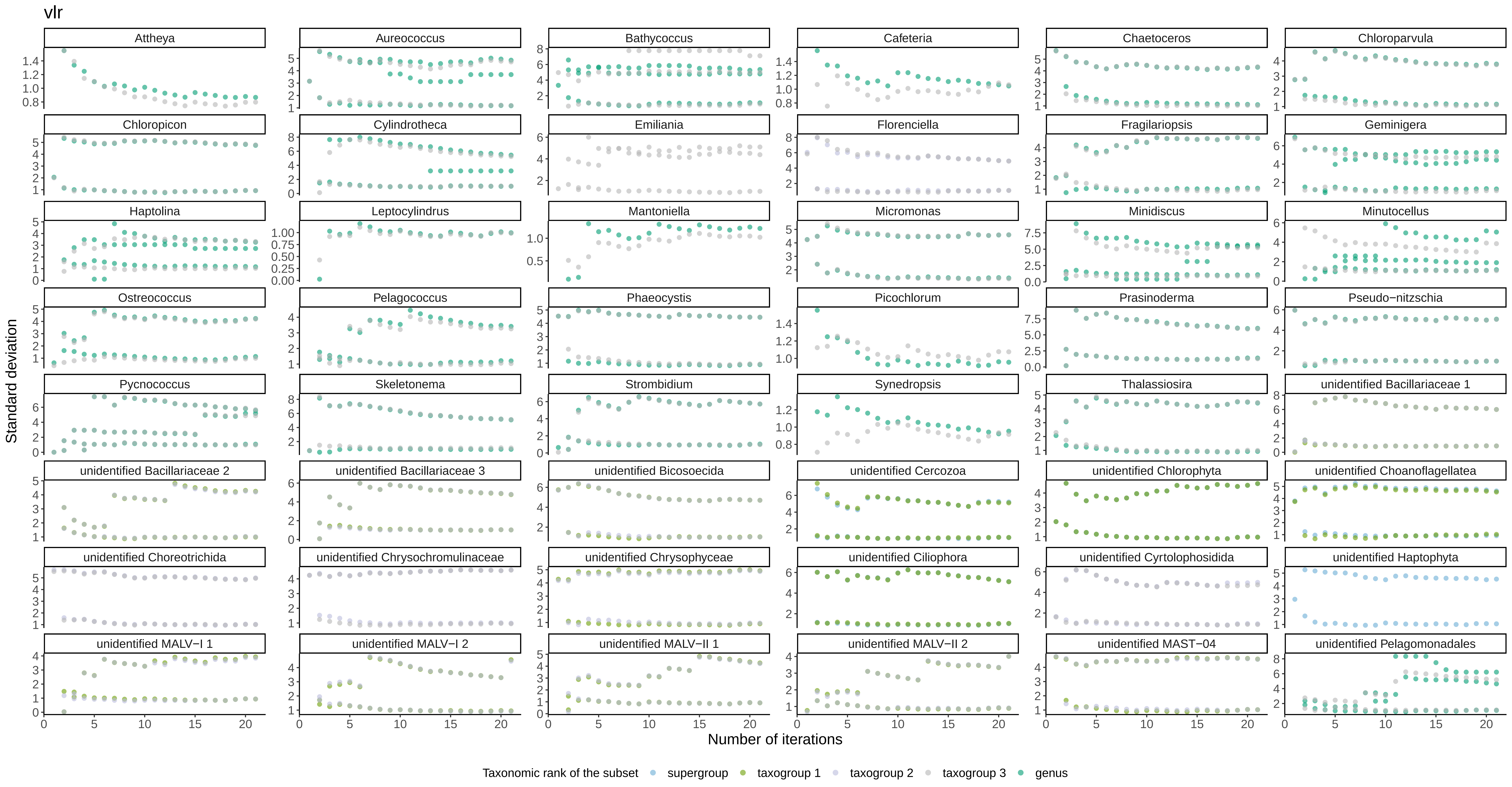

### sim_uniq_numbers_cor.pdf

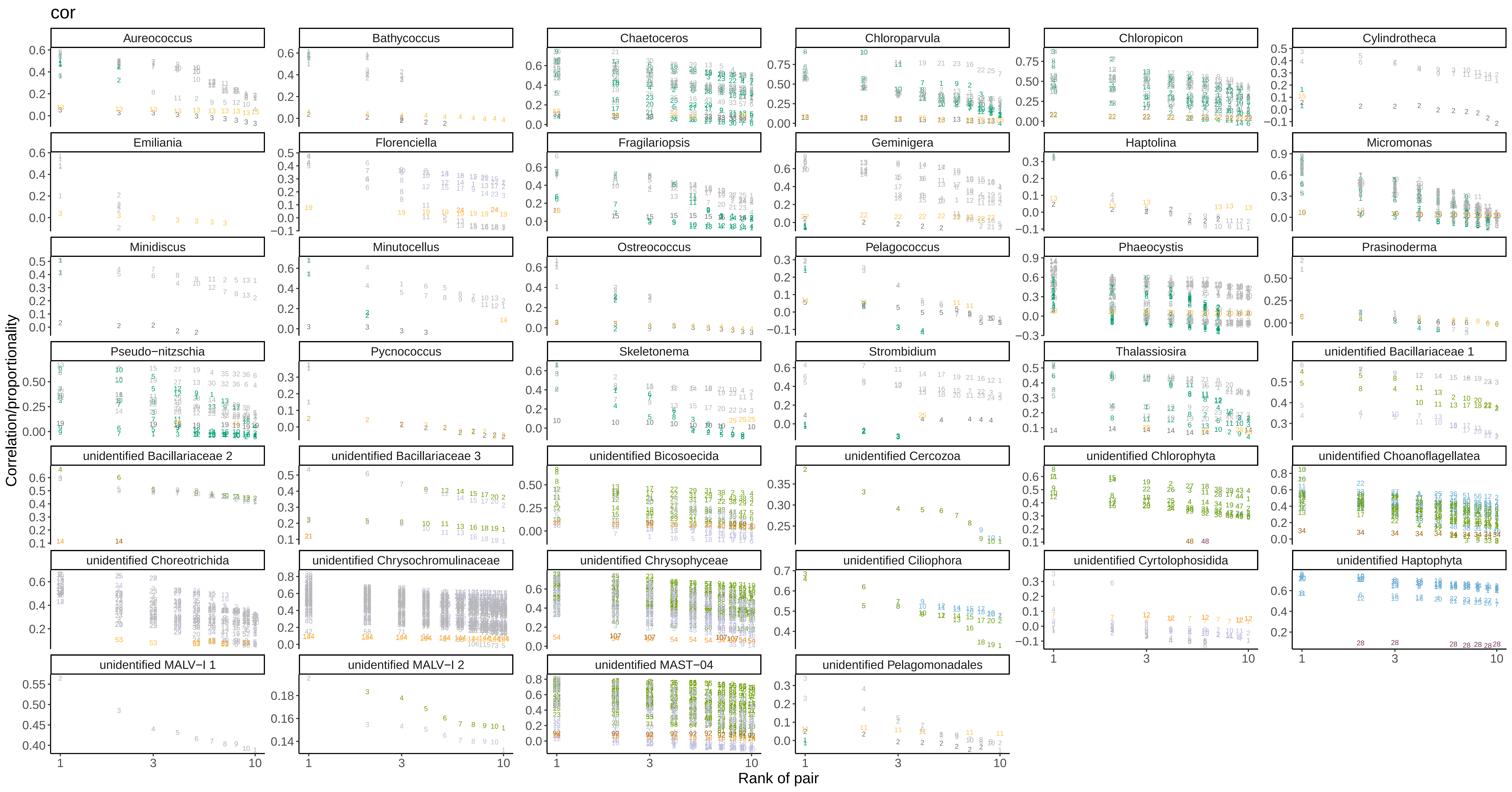

### sim_uniq_numbers_phi.pdf

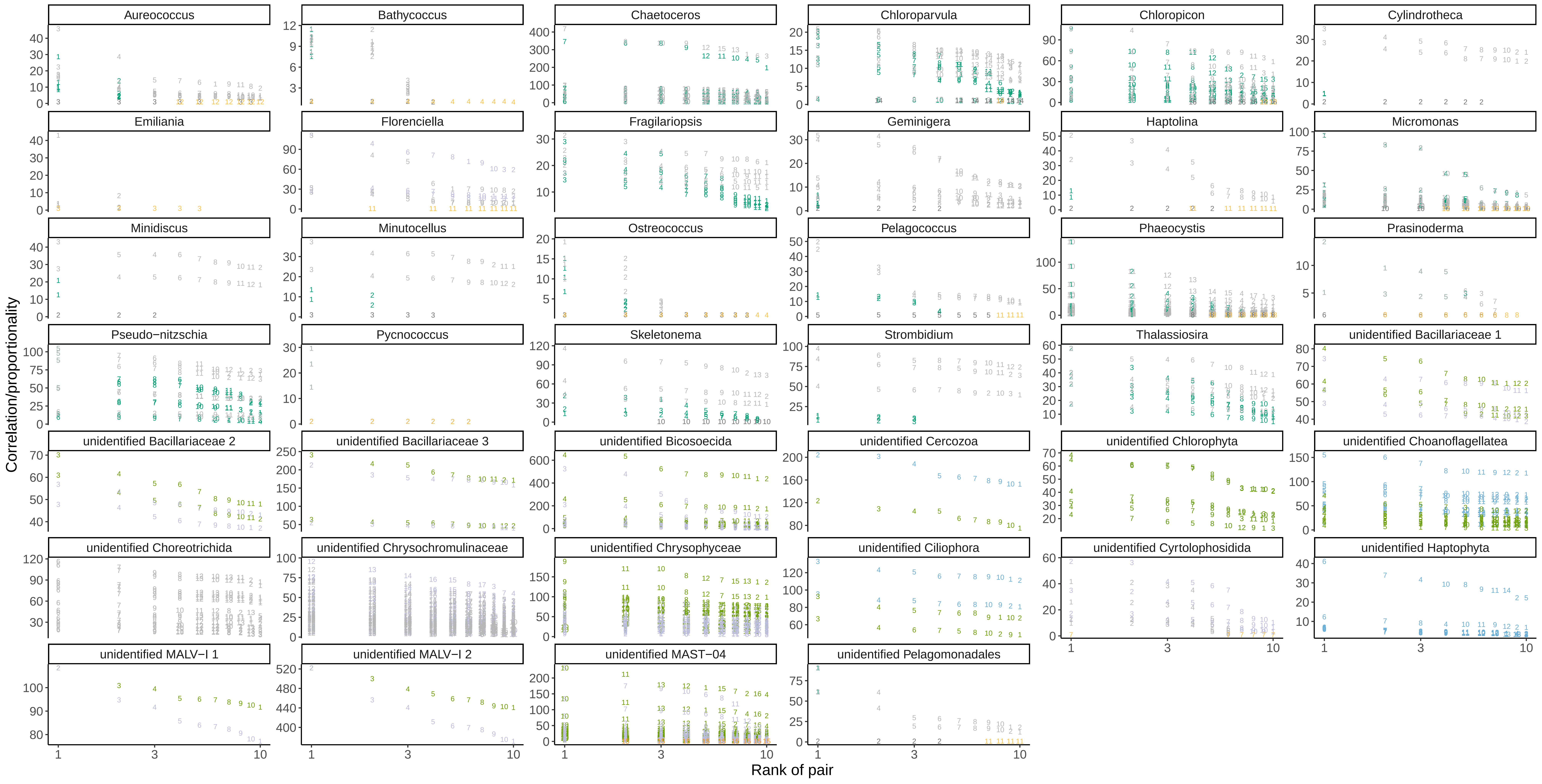

### sim_uniq_numbers_phs.pdf

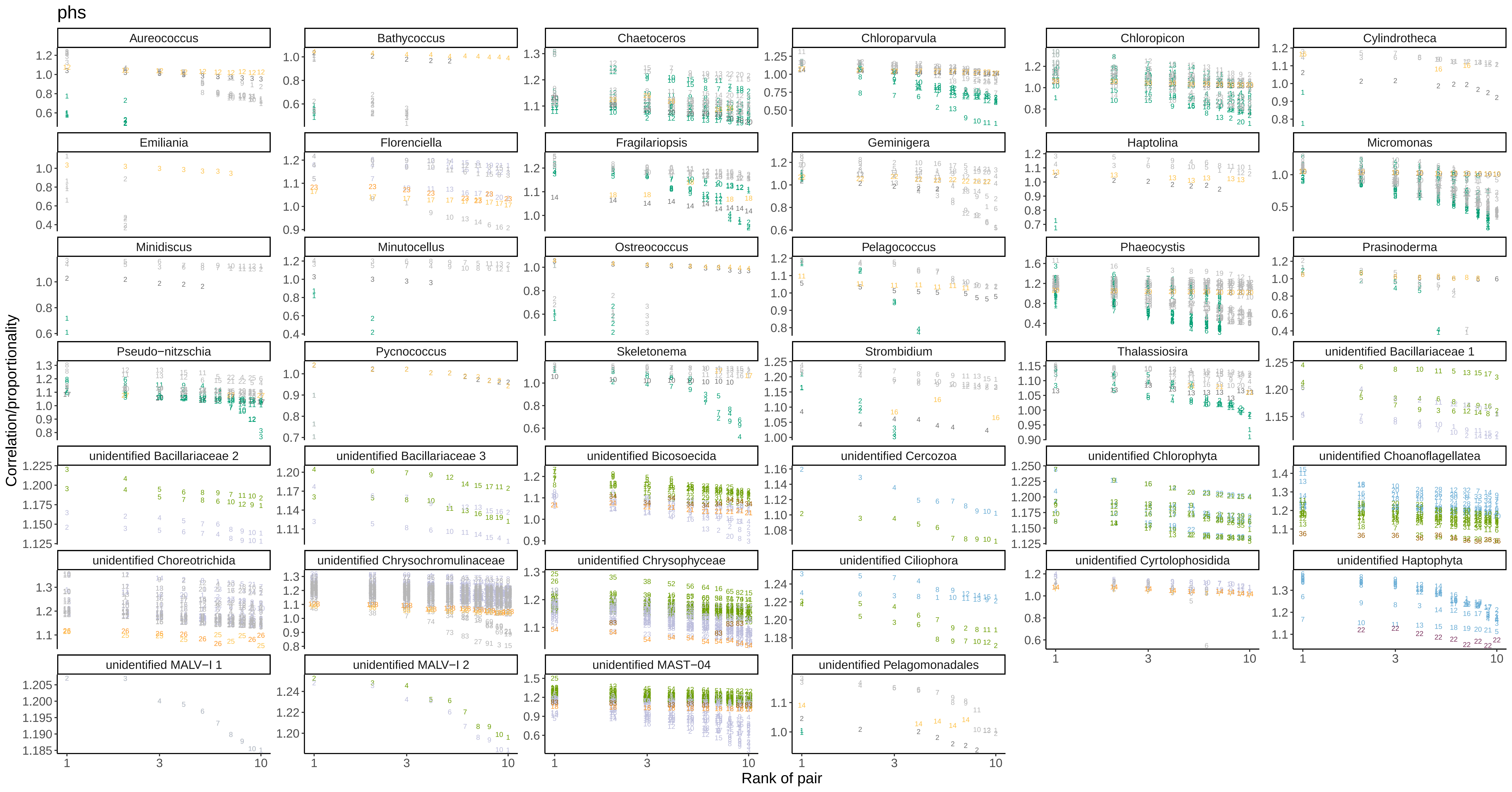

### sim_uniq_numbers_rho.pdf

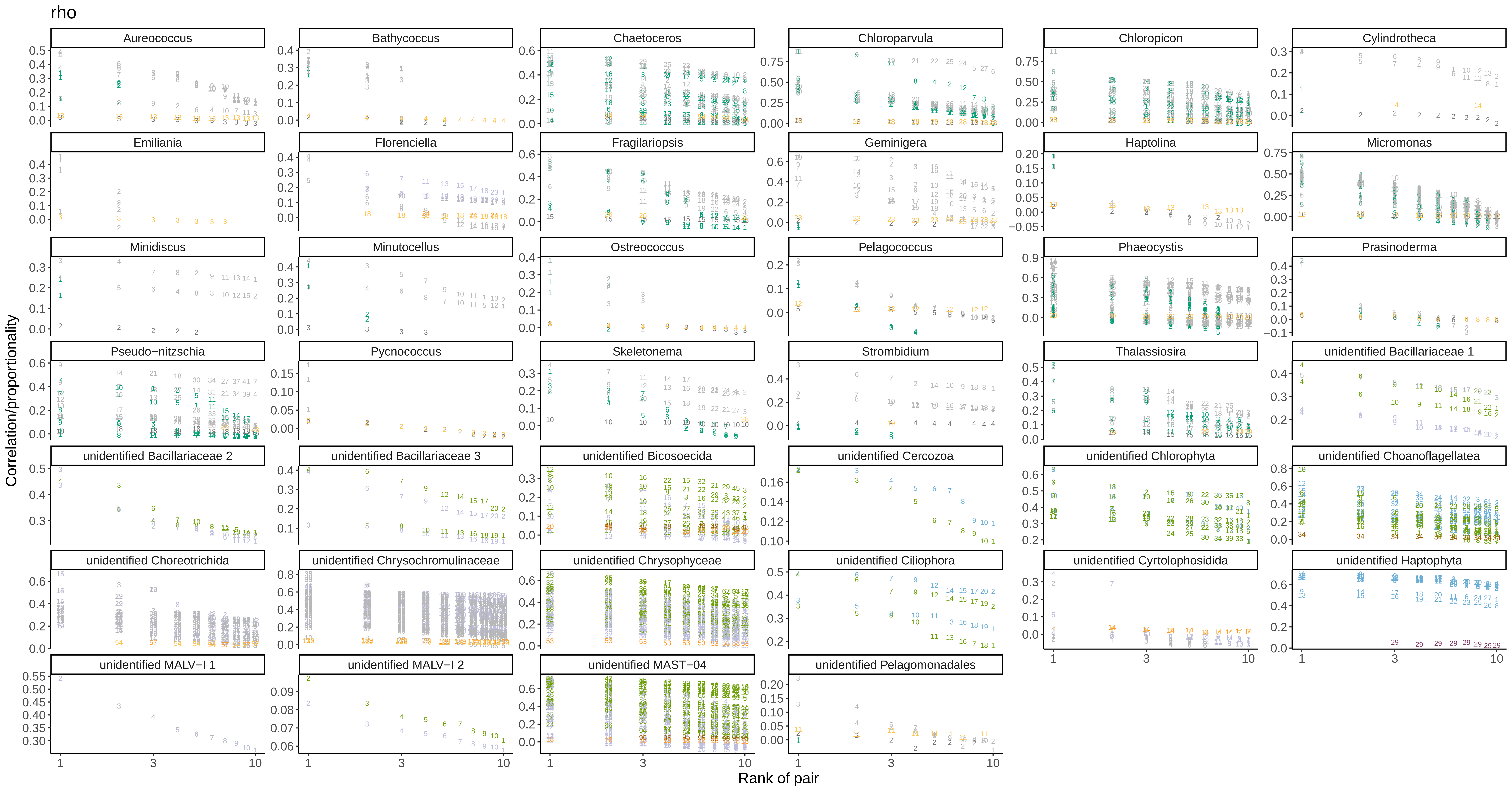

### sim_uniq_numbers_vlr.pdf

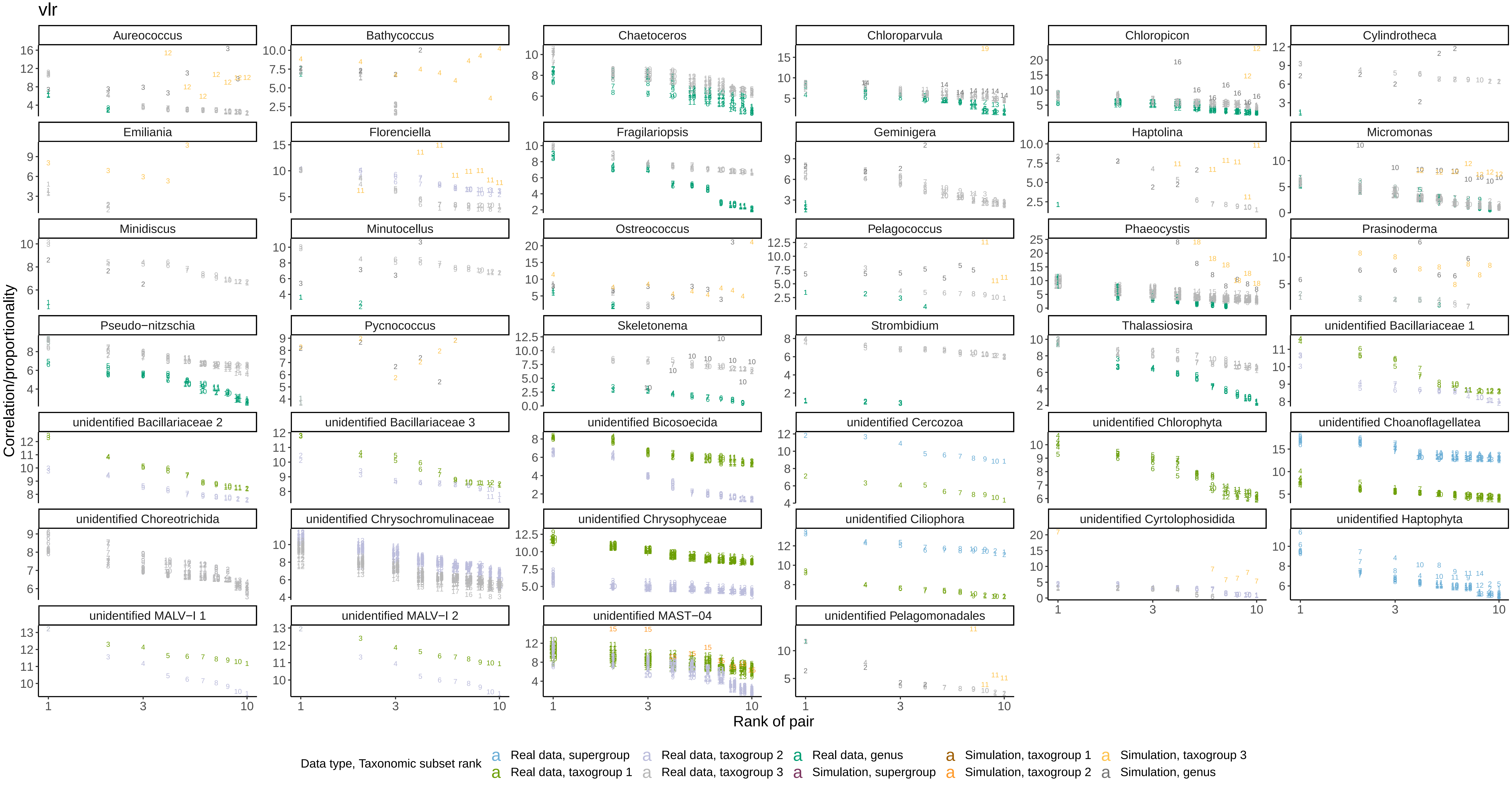

### SMAG_control_scatterplots.pdf

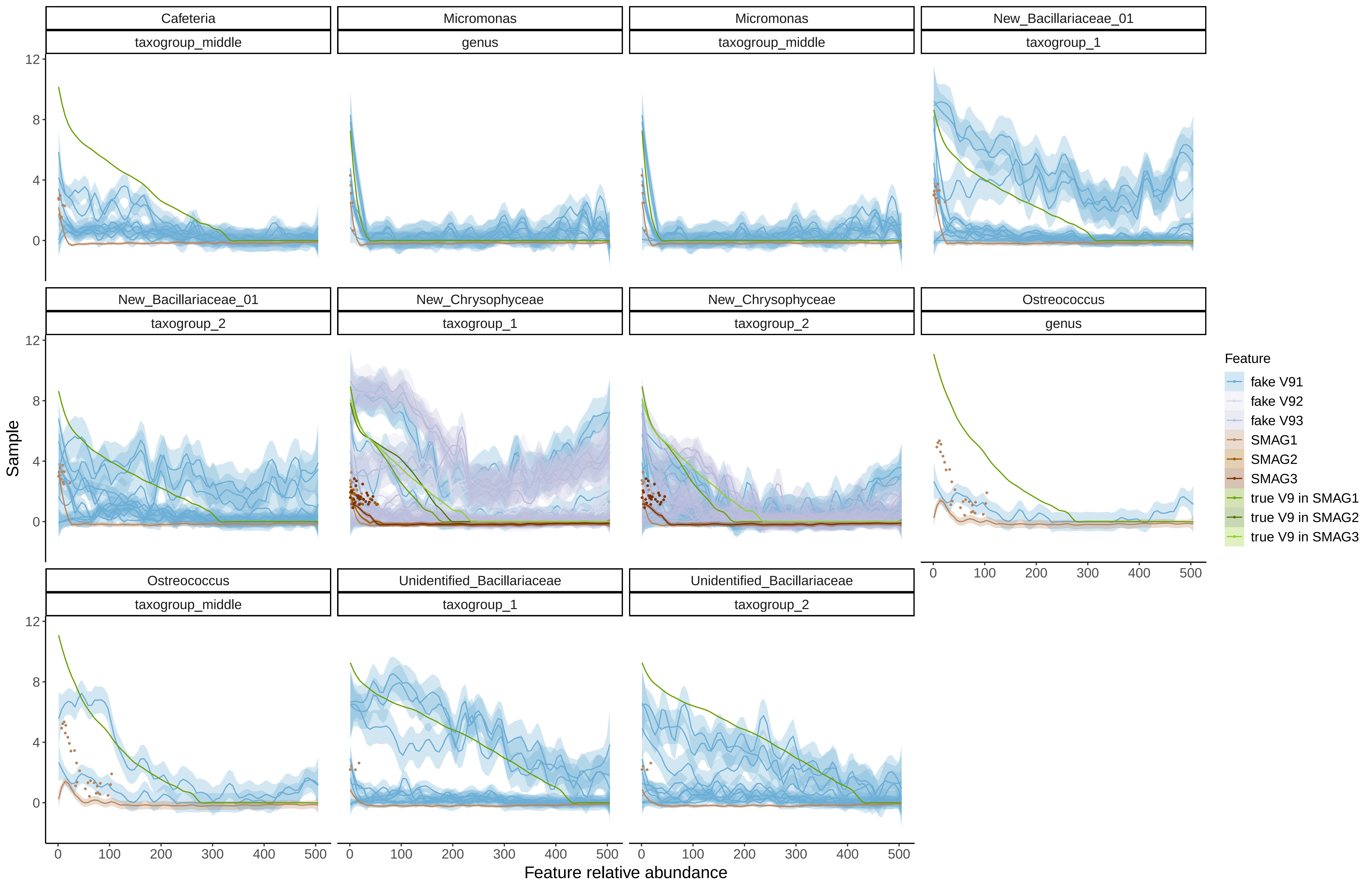

### vlr_calc_plots.pdf

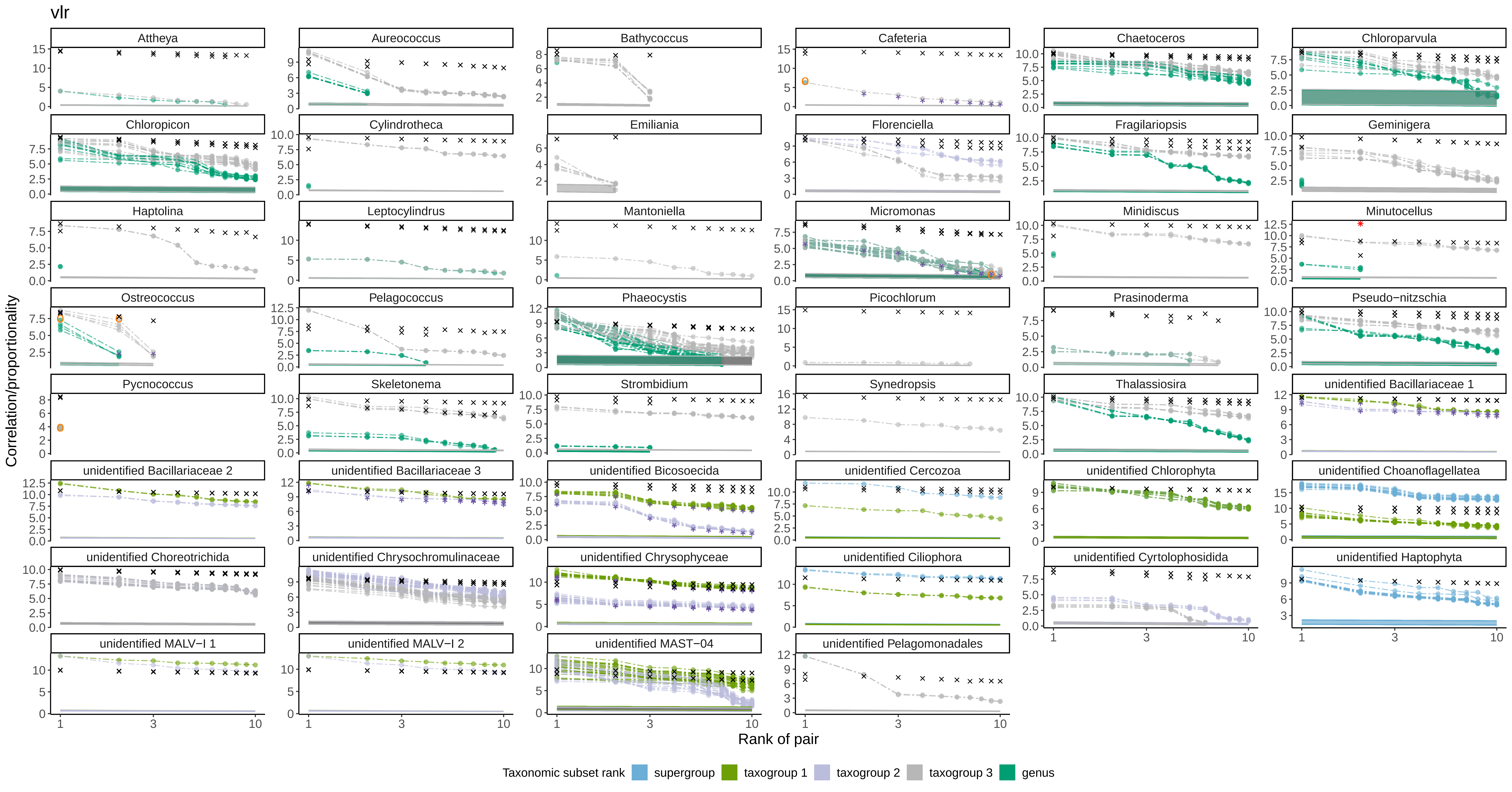

### vlr_uniq_facet_plots2.pdf

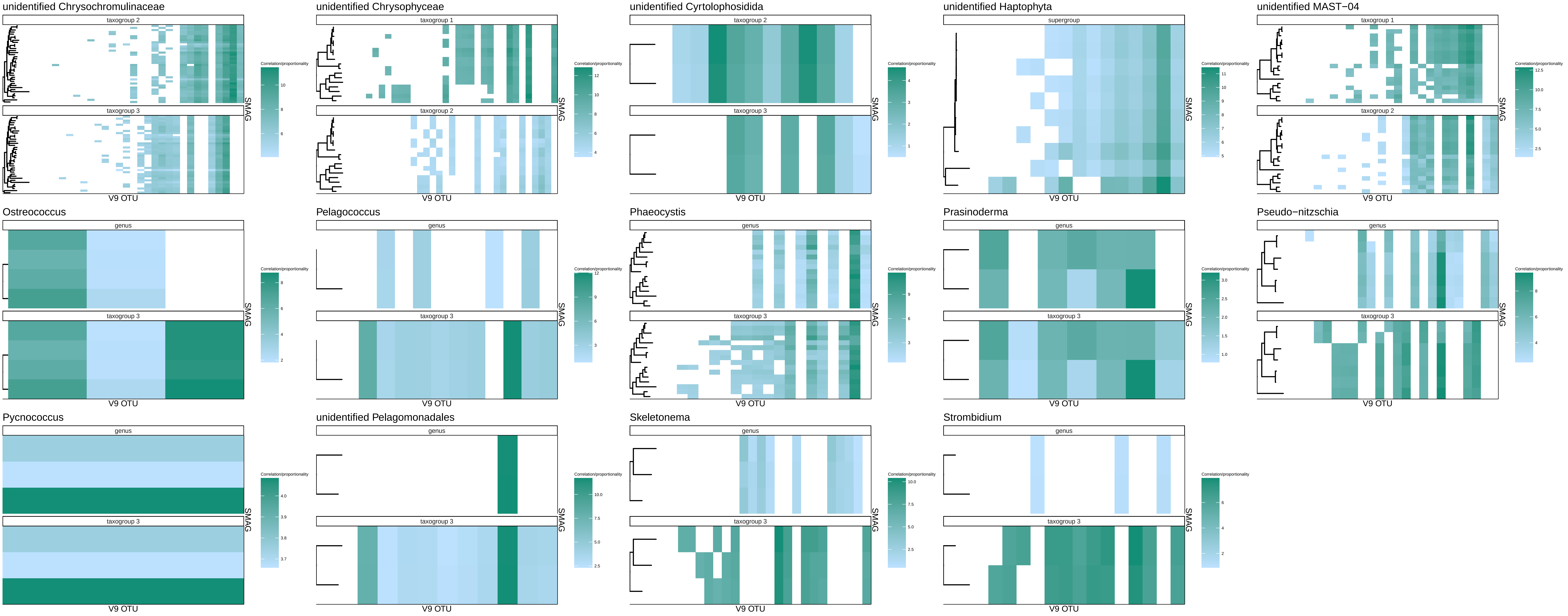

### vlr_uniq_facet_plots.pdf

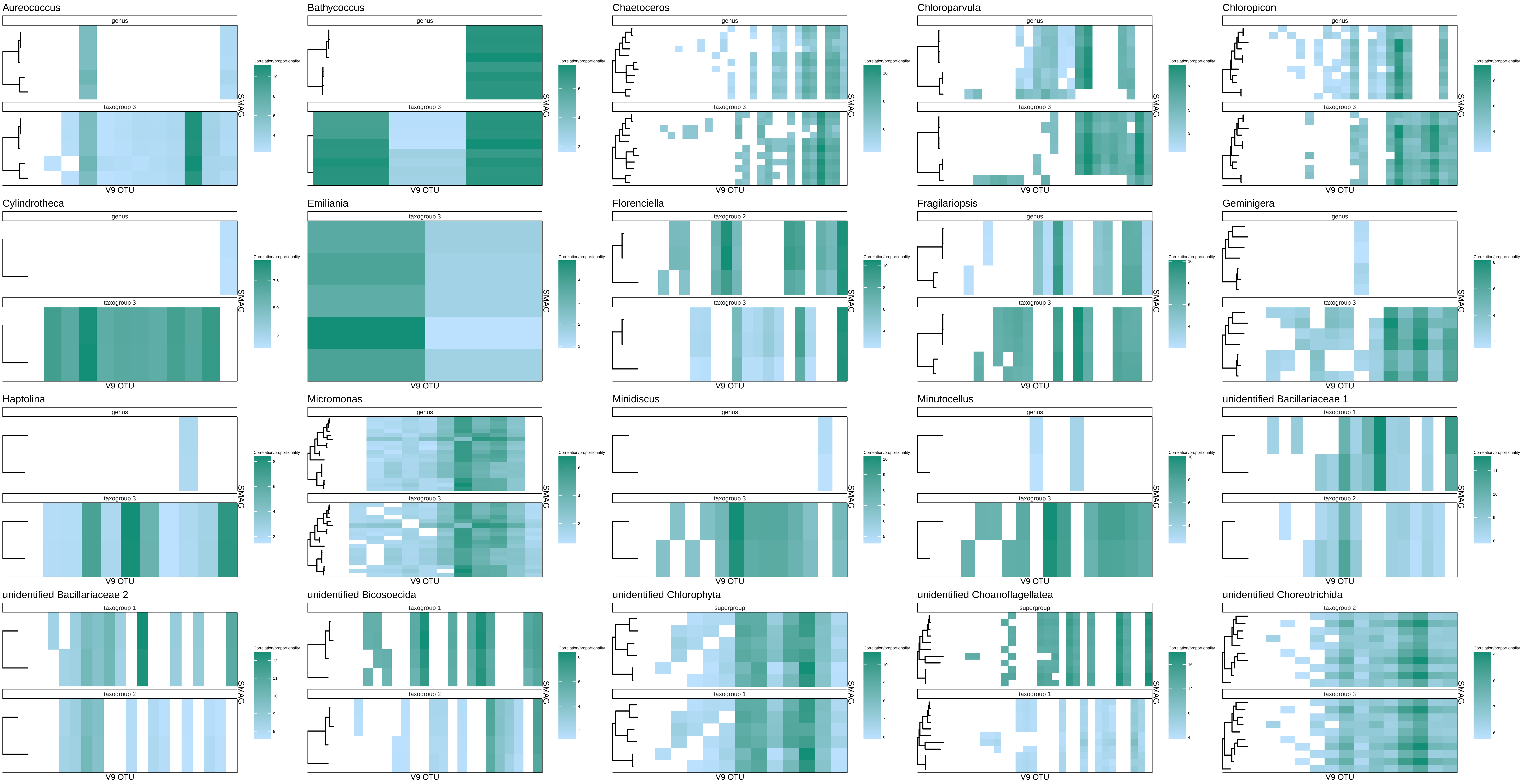
